# CAYSS: package for automatic Cytometry Analysis of Yeast Spore Segregation

**DOI:** 10.1101/2024.09.27.615352

**Authors:** Xavier Raffoux, Matthieu Falque

## Abstract

Meiotic recombination is a powerful source of haplotypic diversity, and thus plays an important role in the dynamics of short-term adaptation. However, high-throughput quantitative measurement of recombination parameters is challenging because of the large size of offspring to be genotyped. One of the most efficient approaches for large-scale recombination measurement is to study the segregation of fluorescent markers in gametes. Applying this to yeast spores by flow cytometry has already been proved to be highly efficient, but manual analyses of density distributions of signal intensities is time-consuming and produces non-perfectly reproducible results. Such analyses are required to identify events corresponding to spores and to assign each of them to a genotypic class depending on their fluorescence intensity. The CAYSS package automatically reproduces the manual process that we’ve been developing to analyze yeast recombination for years, including Maximum-Likelihood estimation of fluorescence extinction (Raffoux et al. 2018a). When comparing the results of manual *vs* CAYSS automatic analyses of the same cytometry data, recombination rates and interference were on average very similar, with less than 3% differences on average and strong correlations (R^2^>0.9). In conclusion, as compared to manual analysis, CAYSS allows to save a lot of human time and produces totally reproducible results.

The CAYSS software is freely available under GPL license from https://forgemia.inra.fr/gqe-base/cayss/-/releases. The package is also provided as Supplementary Material 1 and a tutorial may be found as Supplementary Material 2.

**TAKE AWAY:** The CAYSS R package measures recombination rate and crossover interference from flow cytometry data of isolated yeast spores segregating for fluorescent markers. The analysis is fully automatic and reproducible. It includes spore identification from FSC and SSC data, genotype determination from fluorescence intensities, and Maximum Likelihood estimation of recombination parameters.

## INTRODUCTION

Meiosis is a particular type of cellular division which produces four haploid cells from a diploid mother cell. Among others, meiosis plays a key role in short-term evolution from standing genetic variation because it can produce new combinations of alleles which already exist in the population. This can be achieved by reassorting alleles of loci located on different chromsomes during homologs segregation at anaphase I, but also by reassorting alleles of loci located on the same chromosome through recombination due to crossover and gene conversions occurring during prophase I.

Meiotic recombination rate can be measured by several approaches, including (1) cytological observation of chromosomes and visualization of crossover (CO) sites by immunofluorescence of proteins that form foci at CO positions (Chelysheva et al. 2012) or by EM observation of late recombination nodules (Anderson et al. 1999), (2) analysis of DNA polymorphisms in segregating populations obtained after crossing parents, or (3) direct genotyping of gametes produced by an heterozygous individual. When high throughput is required, the latter method is often the most appropriate, but depending on the organism studied, it is not always feasible. Even though single-cell sequencing or genotyping is now achievable in more and more organisms, phenotypic markers determined by a single (or major) genetic locus can still be very efficient to proxy the genotype of segregating loci, similarly to what was initially done by the pioneers of genetics (Morgan 1910).

Some phenotypic traits very efficient for that purpose are fluorescent phenotypes produced by the expression of genes coding for fluorescent proteins. This has for instance been developed and widely used in the plant *Arabidopsis thaliana* (Berchowitz and Copenhaver 2008; Yelina et al. 2013) and in the yeast *S. cerevisiae* (Thacker et al. 2011a; Raffoux et al. 2018a). In yeast, one such approach is tetrad analysis, carried out by observing fluorescent tetrads under the microscope to determine the fluorescence status of each of the four spore for three colors (Thacker et al. 2011a). The other approach is random spores analysis, which consists of disrupting ascii and analyzing fluorescence status of individual spores, for instance by flow cytometry (Raffoux et al. 2018a). In both cases, diploid strains hemizygous for three fluorescent markers linked on the same chromosome will produce spores segregating for the markers, and the numbers of tetrads or spores in each of the expected genotypic classes allow to calculate recombination rates in both intervals between markers, as well as the coefficient of coiencidence (CoC), which is a measurement of crossover interference in a pair of intervals between markers (Weinstein 1918). The precision of these results tightly depends on the numbers of observations, particularly for CoC which is based on double recombination events, that can be very rare if recombination is low in the region, and/or if interference is strong. Thus, the number of tetrads or spores analyzed must be as high as possible. To analyze fluorescent tetrads automatically, an AI-driven software tool has recently been proposed (Szücs et al. 2024), but most works use manual observation of tetrads by eye (Vincenten et al. 2015; Rogers et al. 2018). Alternatively, one may use an imaging flow cytometer (Basiji 2016), but such equipments and associated data treatment servers are very expensive and difficult to calibrate.

On the other hand, random spore data acquisition is performed at very high throughput (more than 10^6^ spores per minute) with a standard flow cytometer (Raffoux et al. 2018a), but each sample requires a careful analysis of distributions of fluorescence intensities at the wavelengths of each marker, to obtain an accurate estimation of numbers of spores in each genotypic class expected from the segregation. Moreover, not all events detected by the flow cytometer are isolated spores, there can be some cellular debris, some remaining vegetative cells, and spores can sometimes form doublets if the spore isolation protocol was not efficient enough. So, before computing recombination rates, it is necessary to (1) select events as putative spores and distinguish them from vegetative cells based on both their level of light forward scattering (FSC) — which reveals the overall size of the objects — and side-scattering (SSC) — which depends on the granularity of their surface, then (2) select spore singlets from spore doublets or bigger cell aggregates, based on the ratio between the height of a SSC peak over time produced by a cell passing through the detector and the area of that peak, and (3) count the numbers of spores having each of the different genotypes — that is different combinations of fluorescent markers — allowed by the segregation.

This is not always straight-forward: for instance, when a mixture of diploid cells without any fluorescent marker (thus producing only non-fluorescent spores) and diploid cells hemizygous for the fluorescent markers are sporulated together in a population, then, depending on the promoter used with the fluorescent markers, remaining fluorescent proteins expressed in the cytoplasm of diploid vegetative cells may be passed onto the spores, thus producing more than two distinct levels of fluorescence.

Altogether, these successive cytometry analysis steps are usually carried out manually by experts, based on drawing gates in 1-D or 2-D plots produced by computer programs provided with the flow cytometers. However, when large numbers of samples have to be analyzed and/or one wants the results to be perfectly reproducible, an automated analysis pipeline of raw cytometry data is necessary. A number of sophisticated tools have been developed to automate different types of health-related cytometry analyses of various biological samples, see reviews in (Cheung et al. 2021, Cheung et al. 2022), but we do not know of any completely automated pipe-line to calculate recombination rates and CoC from raw flow cytometry results of fluorescent isolated yeast spores. Thus we developed the CAYSS R package to be able to analyze in a high-throughput and completely reproducible fashion large amounts of flow cytometry raw data obtained with the method and the fluorescent tester strains (available upon request) previously developed in our lab (Raffoux et al. 2018a). With the hope of further facilitating the use of that method by the community, the CAYSS package is freely available as Supplementary Material 1 and as a release of the project at https://forgemia.inra.fr/gqe-base/cayss/-/releases. A quick-start tutorial is available as Supplementary Material 2.

## MATERIALS AND METHODS

### 1. Yeast strains and populations

The spores analyzed by flow cytometry to produce the raw data used in this work were obtained from two different types of diploids hemizygous for fluorescent markers: (1) diploids obtained in a previous work (Raffoux et al. 2018b) by crossing each of the 26 strains of the collection of the *Saccharomyces* Genome Resequencing Project (Cubillos et al. 2009; Liti et al. 2009) with the eight tri-fluorescent SK1 tester strains described in (Raffoux et al. 2018a), and (2) diploids obtained by crossing a population of non-fluorescent spores of diverse genotypes with the fluorescent testers described above. To do so, the SGRP-4x population (Cubillos et al. 2013) was first crossed with the SK1-VI-C1Y2R3 (Raffoux et al. 2018a) strain, and 1 to 8 generations of panmictic sexual reproduction were carried out by sporulation, tetrad disruption, and mixing of isolated spores. At each of the 8 generations, 50 000 non-fluorescent spores were FACS sorted and crossed on solid YPD Petri dishes containing Chloramphenicol with each of the eight tri-fluorescent tester strains to obtain diploids hemizygous for the fluorescent markers.

### 2. Culture Media and drugs

Solid YPD contained 1% yeast extract, 2% peptone, 2% glucose, 2% Agar. Solid SPOR medium contained 0,25% yeast extract, 0,1% glucose, 1% potassium acetate, 2% Agar. Chloramphenicol was used at a final concentration of 50µg/mL, and Hygromicin B at a final concentration of 200µg/mL.

### 3. Isolation of spores

After incubation at 30°C 2 days, a quarter of each Petri dishe was scrapped using a tip, and cells were resuspended in a 96 well plate containing 500 µL ddH20. We then diluted 100 µL of each well with 700 µL ddH20 in a second 96 well plate, and again 100 µL of each well of the second plate with 700 µL ddH20 in a third plate. Patches of 50 µL cell suspension were then spotted on SPOR Petri dishes (9 patches per Petri dish) with hygromycin to select diploids against residual haploid tester cells. After ten days of incubation at 30°C, cells were picked up from the lawn by scraping patches with a bended pipette tip, and suspended in tubes in a rack of 96 tube strips (Macherey-Nagel Ref 740637) containing 375 µL of ddH20 with 5mg/mL 20T zymolyase (Euromedex, Souffelweyersheim, France) and 100 µL glass beads suspension (0.5 mm diameter, Dutscher 67172 Brumath Ref 1606106). To disrupt tetrads and enrich spores (Rockmill et al. 1991), plates were shaken 1 min at a frequency of 23 Hz in a TissueLyser (Qiagen), incubated 30 min at 30°C, shaken again (1 min, 23 Hz), incubated 30 min at 30°C, shaken (1 min, 23 Hz), then centrifuged 5 min at 4500 rpm. Supernatant was discarded by pipetting and the pellets were resuspended by shaking (1 min, 23 Hz) in 200µL ddH2O. The suspension, mostly containing vegetative cells, was discarded. Spores, which adhere to the plastic of the tube, were stripped by shaking 2 min (23 Hz) the tubes containing 400µL ddH20 with 0.01% NONIDET NP40 (Sigma-Aldrich, Saint Quentin Fallavier, France). The spore suspensions were then analyzed by flow cytometry.

### 4. Manual flow cytometry analyses

The spore suspensions were analyzed either with a single-tube MoFlo ASTRIOS flow cytometer, in which case acquisition and manual analyses were carried out with the associated software ‘Summit 6.3’, or with a 96-wells plate CytoFLEX flow cytometer, in which case acquisition and manual analyses were carried out with the associated software ‘CytExpert 2.5’.

For manual analyses, we first selected events corresponding to the size and granularity of spores using a gate in the SSC-Height-Log *vs* FSC-Height-Log plot (Top left of Figure S1 in Supplementary Material 3), then we discarded events containing more than one cell using a gate in the SSC-Height-Log *vs* SSC-Area-Log graph (Top right of Figure S1 in Supplementary Material 3). Fluorescence intensity of each spore in mCherry, yECerulean, and Venus channels (excitation at 561, 405, and 488 nm respectively, emission at 614/20, 448/59, and 526/52 nm respectively) was then analyzed: gates were manually drawn on the three 2-D fluorescence intensity scatter plots (mCherry-Height-Log *vs* CFP-Height-Log), (mCherry-Height-Log *vs* YFP-Height-Log), (CFP-Height-Log *vs* YFP-Height-Log), and spores were assigned to one of the four clouds of points in each graph (Bottom panel of Figure S1 in Supplementary Material 3). Finally, we used combinations of the twelve clouds decsribed above to assign spores to one of the eight possible genotypic classes (Figure S2 in Supplementary Material 4).

### 5. CAYSS package working principle

The workflow implemented in CAYSS was designed to execute successively in an automatic and perfectly reproducible fashion the same steps that we have been developing and improving for several years in our manual flow cytometry data analyses to study meiotic recombination (Raffoux et al. 2018a, 2018b). The main steps of this workflow are the following:

First, spores are pre-selected based on fixed FSC and SSC thresholds defined by parameters *minSporeFscHLog, maxSporeFscHLog, minSporeSscHLog*, and *maxSporeSscHLog*. This allows to discard most of the cellular debris (mostly smaller than spores) and remaining vegetative cells (mostly larger than spores).

Second, the population of events considered as spores is delimited by the following procedure: (1) define a grid (e.g. of 100×100 bins defining 10000 elements) in the SSC-Height-Log *vs* FSC-Height-Log 2-D plot within the given X and Y range defined by the FSC and SSC thresholds, (2) set the initial selected gate as the element of the grid containing the largest number of events *N* _*max*_, (3) iteratively enlarge the selected gate by including for each iteration *i* all other elements of the grid containing *N* ≥ *p*_*i*_×*N*_*max*_ events, with *p*_*i*_ decreasing from 0.95 to 0.1 by step of 0.05, and calculate the ratio 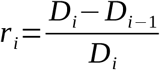 where *D*_*i*_ is the average distance between all events selected at iteration *i* and the center of the initial gate, and *D*_*i*−1_ is the same at the previous iteration *i*-1. (4) Stop iterating when *r*_*i*_ becomes lower than the threshold given by the parameter *slopeFindPeak* of the main function. Events belonging to any of the selected gates are then considered as being spores.

Third, to distinguish single spores from possible spore doublets, or other types of cell aggregates, the usual method is to discriminate based on the width/length ratio of the objects. The events to be selected are then expected to follow a straight line in the graph SSC-Height-Log *vs* SSC-Area-Log. The procedure to select spores along the main linear cloud is very similar to the previous one, except that one expects the selected cloud to be linearly shaped: (1) define a grid in the graph, as previously, (2) set the initial selected gate as the 10 elements of the grid containing the highest numbers of events, (3) compute a linear regression from the 10 centers of these squares, (4) enlarge the gate iteratively as previously except that the distance used is not the distance to the center of the initial gate, but the distance to the regression line. The threshold to stop iterating is the same parameter *slopeFindPeak* as previously. Events selected by both this gate and the previous one are considered as spore singlets, and all the subsequent fluorescence intensity analyses will be restricted to those events. To save computation time in case of large data sets, CAYSS can take random subsamples of these single spores, as specified by the parameter *nbUsedForGating*.

Then, analyses of fluorescence intensity are carried out successively for each of the two or three fluorescent markers used. For each of them, the procedure is the following: (1) the density distribution of log-transformed fluorescence intensities of the single spores is smoothed, and peaks and valleys are detected, (2) the peaks whose height is lower than the height of the highest peak times the parameter *cutOff* are discarded. (3) If no or only one peak is detected, the analysis stops and proceeds to the next sample. Otherwise, the two peaks corresponding to the highest fluorescence intensities are selected and will be considered as containing the spores which segregate the presence and the absence of the considered fluorescent marker, as opposed to other classes of events which may produce peaks of lower fluorescence intensities (on the left side of the graph, e.g. spores derived from non-fluorescent diploids), (4) for each selected peak, the spores with fluorescence intensities ranging within the mode of the peak plus or minus the standard deviation of the peak times the parameter *widthAroundPeak* are selected and assigned to class 1 (marker present) or class 2 (marker absent). All other spores are discarded, so each of the two or three markers further contributes to eliminating events that are not spores.

Finally, each selected spore is assigned to a two-loci (4 classes) or three-loci (8 classes) genotypic class as explained before (Raffoux et al. 2018a), and recombination rates are computed in each interval between markers. When three markers are used, the CoC is computed, and a different estimation of recombination rates and CoC is also provided, based on Maximum-Likelihood (ML) estimation when considering probabilities of fluorescence extinction for each of the three markers, as described before (Raffoux et al. 2018a). We observed in that previous study that such fluorescence extinction does happen in some fluorescent strains, and this can be easily identified from the CAYSS output data because in that case, ML-inferred values of the fluorescence extinction parameters are significantly different from zero.

## RESULTS AND DISCUSSION

### 1. Interface and launching parameters

CAYSS is designed to automatically analyze batches of FCS raw output files from flow cytometers. This is particularly useful when working in 96-well sample plate format, but can also be used with single-tube devices. All FCS files of a batch must be in the same folder and their names must contain a unique sample ID at the end, just before the ‘.fcs’ extension, separated from the beginning of the file name by an underscore. For example, the file ‘Sample1_output-FCS_file25.fcs’ corresponds to the ID ‘file25’. In the case of 96-well sample plates, that ID may be the plate coordinate (e.g. ‘A1’). In addition, a plate design file containing the list of samples of the batch must be prepared as a tab-separated text file with two columns: the complete sample name, and the corresponding unique sample ID which must correspond to that of one of the FCS files. The complete sample name must be composed of several tokens separated by underscores (e.g. ‘Sample1_SK1_ChrVI_2024-08-23_RCY’). Within this name, three tokens indicate (1) the date of the experiment, (2) the chromosome carrying the markers, and (3) a mandatory character string containing three single-letter codes, each of them representing a fluorescent marker in the order of their position on the chromosome. For instance, the fifth token ‘RCY’ of sample name ‘Sample1_SK1_ChrVI_2024-08-23_RCY’f indicates that the markers are placed in the order R-C-Y on the chromosome. It is thus easy to have samples with different marker orders in the same batch, whereas all other parameters are given as arguments of the main function ‘*AnalyzeRecombination()*’ and will be the same for all samples of the batch.

To our experience, the launching arguments of CAYSS almost never need to be changed when using the same markers and flow cytometer hardware configuration. However, in some rare cases, some particular samples may require different values for some of these parameters (for instance ‘*minSporeFscHLog*’, ‘*cutOff*’ or ‘*widthAroundPeak*’). Therefore, after a first run of the batch and a visual check of the output graphs, if some samples fail to be analyzed, one can specify different values of some parameters for each of these samples, by filling an additional column in a copy of the plate design file which is generated at each run. After one or two such iterations, the last run usualy gives optimal results for all samples, and the analysis is completely reproducible. More details may be found in the documentation of the function ‘*AnalyzeRecombination()*’.

For each fluorescent marker used, the argument ‘*fluo1H*’, ‘*fluo2H*’, or ‘*fluo3H*’ of the main function ‘*AnalyzeRecombination()*’ provides the corresponding single-letter code (e.g. ‘R’), the complete marker name (e.g. ‘mCherry’), and the name of the corresponding channel of the cytometer (e.g. ‘FL8.H’). Additional arguments of that function indicate also the channels used for FSC.Height, SSC.Height, and SSC.Area, the number of spores used to analyze distributions of intensities and set the gates, and some parameters used to tune the algorithms of automatic peak detection and gating.

During the analysis, three types of output files are generated: (1) a one-page-per-sample combination of graphics showing the shapes of the different distributions of FSC, SSC, and fluorescence intensities, together with numerical values of the recombination parameters obtained (Figure S3 in Supplementary Material 5). This makes it very quick to visually check if the results are faithful. (2) A tabulated text file with one line per sample, and all numeric results as columns, and (3) in the case of three markers, a one-page-per-sample combination of graphics showing the quality of parameters convergence for the Maximum-Likelihood estimation of recombination and fluorescence extinction parameters (Figure S4 in Supplementary Material 6).

Detailed information about all input parameters or files and the output files produced is available in the documentation of the main function ‘*AnalyzeRecombination()*’ of the CAYSS package.

### 2. Selection of spores

Random spores segregation analysis by flow cytometry requires to disrupt tetrads into isolated spores, and to get rid as much as possible of remaining vegetative cells (Raffoux et al. 2018a) which can be non-sporulated diploids or even remaining haploid cells if no proper diploid selection was applied. Even though spores are enriched by the wet-lab protocol, it is necessary to use the flow cytometer to select spores based on their morphological differences with the vegetative cells. This is automatically performed by CAYSS in the three following steps (see details in Materials and Methods).

First, fixed thresholds are applied to the FSH and SSC peak height values. Our repeated experiments over years showed that fixed threshold values, once determined by looking at a few samples manually, almost never need to be changed if one uses always the same flow cytometer hardware configuration and the same channels with the same fluorochromes.

Second, the main central cloud of spores is automatically localized (see Materials and Methods) and events within that region are gated, as indicated by green dots in Figure 1. The events selected as spores have thus similar sizes (FSC, X-axis) and granularities (SSC, Y axis). We showed previously (Raffoux et al. 2018a) that one can interpret the different populations of events as cellular debris on the bottom-left side, then a more or less circular cloud of events which are spores in the middle, and then a more diffused cloud of points on the top-right side, which mostly contain vegetative cells and/or aggregates.

**Figure 1.**
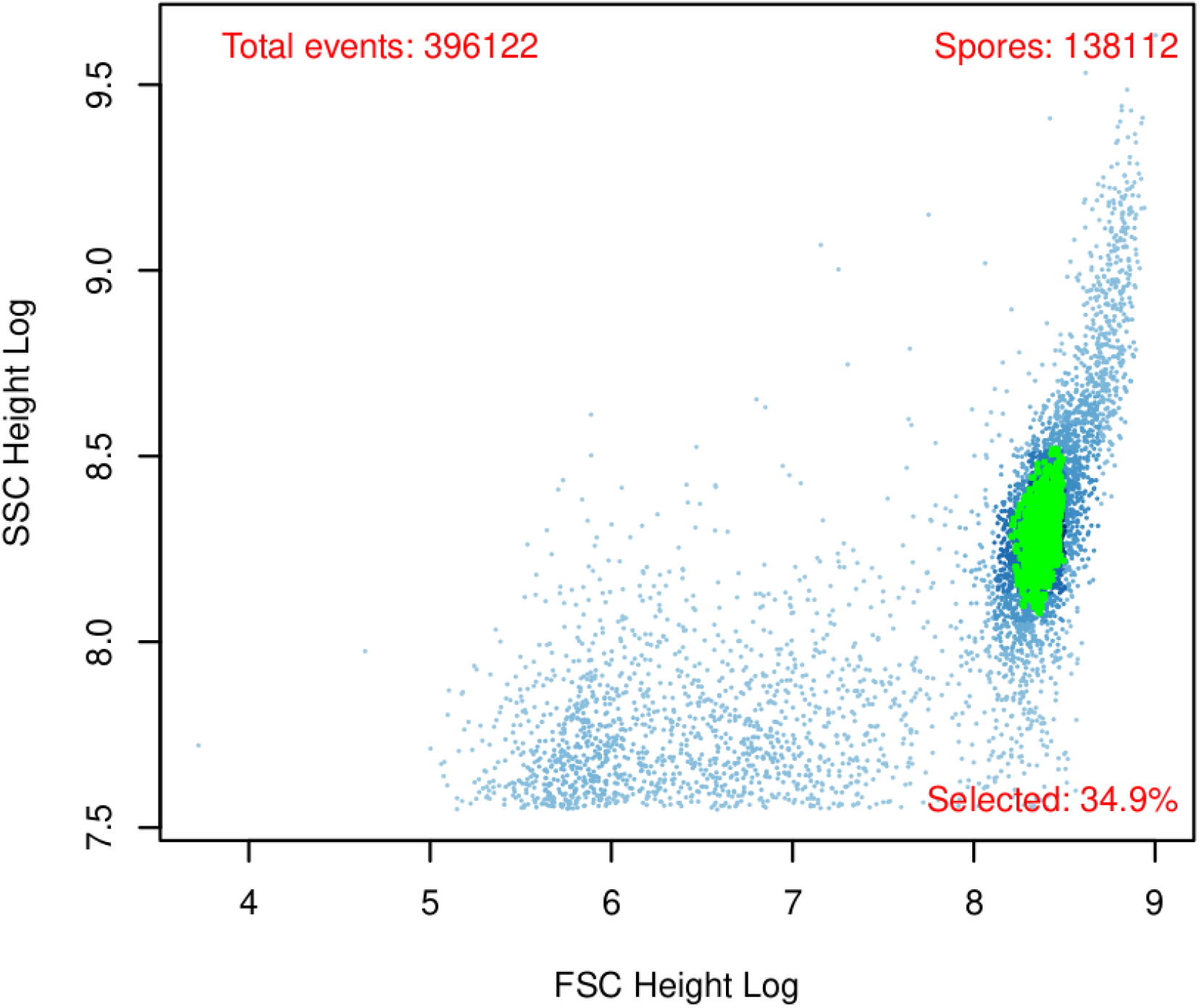
SSC-Height-Log (Y-axis) *vs* FSC-Height-Log (X-axis) plot showing the result of the procedure used by CAYSS to fine-select spores among pre-selected events, based on automatic identification of the main cloud. Putative spores (in grid elements colored in green) are distinguished from cellular debris (on the bottom-left side) or larger types of cells (e.g. vegetative cells, on the top-right side).

Third, to discard spore doublets from isolated spores, the SSC-Height *vs* SSC-Area graph (Figure 2) is used to distinguish the top line-shaped cloud of isolated spore (singlets) from the cloud of doublets below the first one. Indeed, SSC peak height indicates the width of the object, and SSC peak area proxies the length of the object. The algorithm implemented in CAYSS then identifies the main linear cloud and selects as spore singlets the events following that cloud (see details in Materials and Methods).

**Figure 2.**
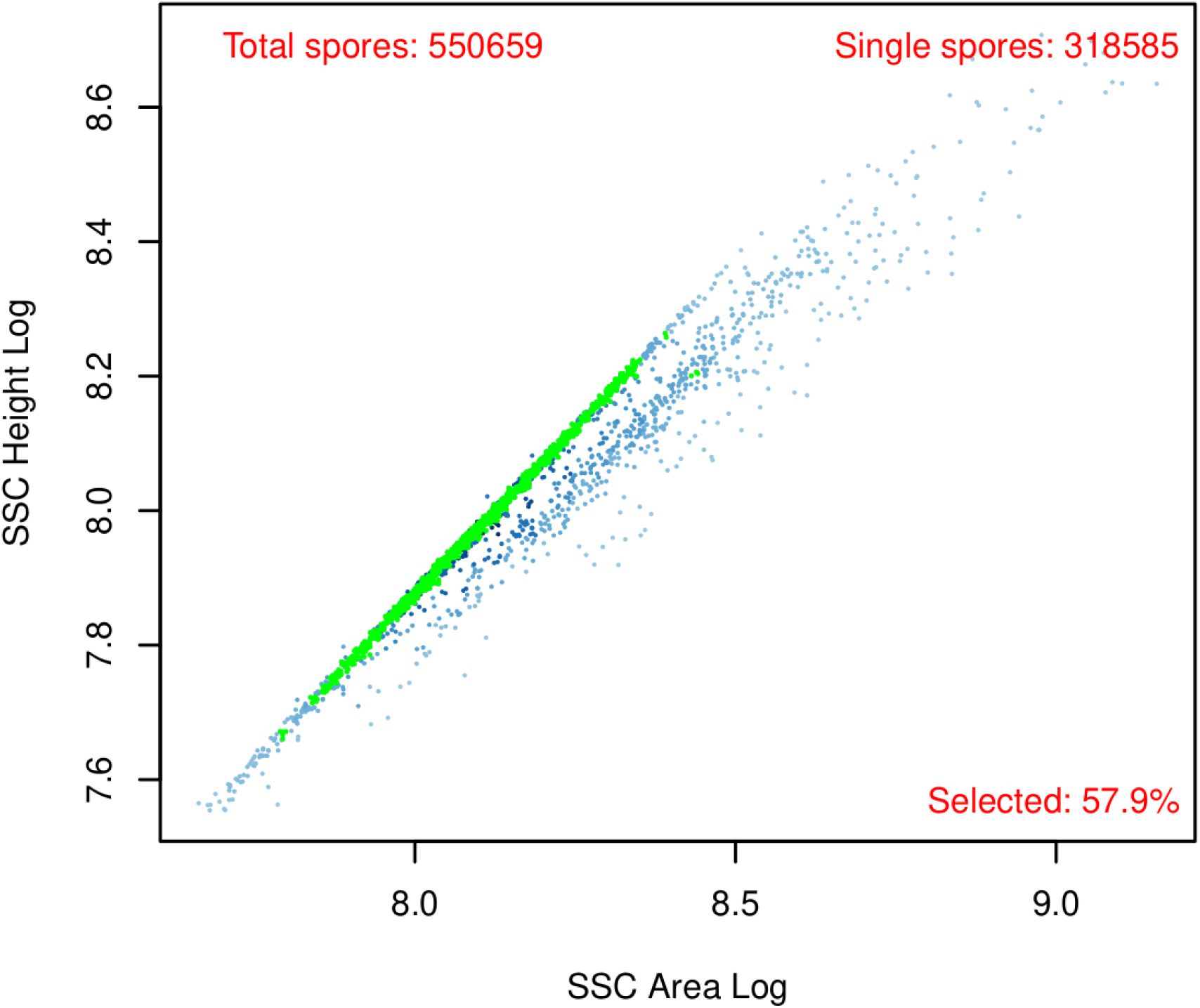
SSC-Height-Log (Y-axis) *vs* SSC-Area-Log (X-axis) plot showing the result of the procedure used by CAYSS to select spore singlets (in grid elements colored in green) which form the main linear cloud, and discard spore doublets or larger cell aggregates which usually form a second elongated cloud below the main one.

After each of these steps, if the number of selected events is less than the parameter ‘*minNbSpores*’, then the process is stopped for that sample, and CAYSS continues with the next one.

### 3. Analysis of fluorescence intensity of markers

Once a sufficient number of spore singlets have been identified and selected, fluorescence intensity density distributions are analyzed for the two or three markers successively. For each of them, the two peaks corresponding to ‘fluorescent spores’ and ‘non-fluorescent spores’ are identified (Figure 3), and each spore is assigned to one of these peaks, or discarded (see details in Materials and Methods). Above each peak is printed the frequency of spores assigned to that peak and the corresponding number of spores counted. This allows to check for possible Mendelian segregation biases, which is a very important and useful indicator of the quality of the results (Raffoux et al. 2018a).

**Figure 3.**
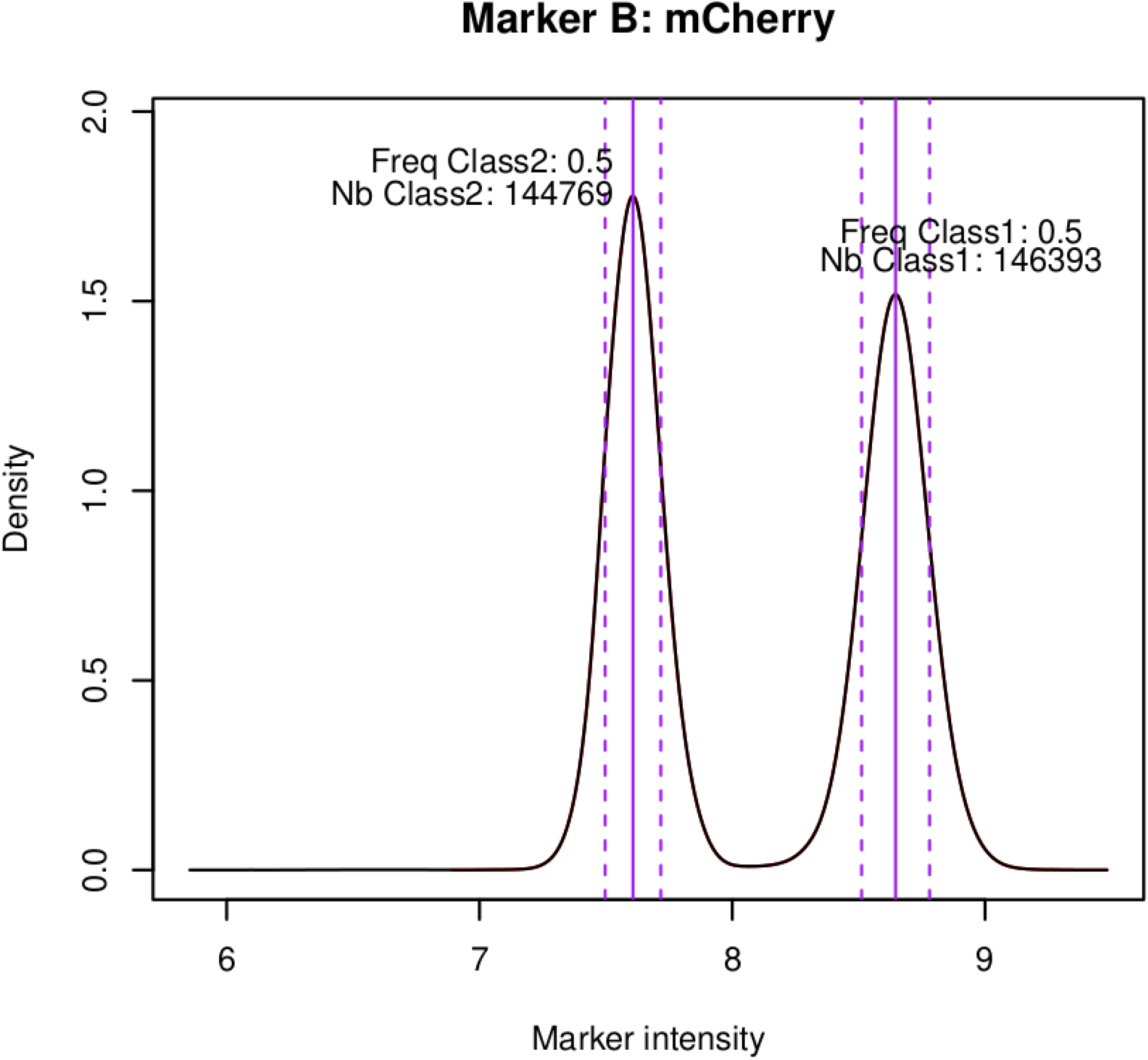
Density (Y-axis) distribution of fluorescence intensity (mCherry-Log, X-axis) of spore singlets showing two peaks, one on the left side corresponding to spores which do not express the fluorescent protein, and one on the right side corresponding to spores which do express the protein. Vertical solid bars indicate the modes of the distribution, and the dotted vertical lines indicate the interval within which the spores are assigned to the peak for segregation analysis.

If for any of the markers, only zero or one peak is detected (e.g. see Figure S5 in Supplementary Material 7), its density curve is plotted in the graphics output file and the rest of the analysis is skipped for that sample.

For each fluorescent marker, the density distributions of fluorescence intensity is plotted as a 1-D density graph in the one-page graphical output file (Figure S3 in Supplementary Material 5). In the case of only two markers, a 2-D scatter plot is also generated (Figure S6 in Supplementary Material 8), showing the four expected clouds of points, two for the parental (non-fluorescent and bi-fluorescent) spores, and two for the recombinant (mono-fluorescent) spores.

### 4. Possible additional fluorescence classes

If the parental diploid strain is hemizygous for a fluorescent marker, the distribution of fluorescence intensity for that marker should have two peaks, corresponding to the spores with and without the marker. However, we observed that when analyzing a population of diploid cells containing a mixture of cells hemizygous for the fluorescent markers, and cells homozygous for the absence of the markers, then a third peak is observed on the left of the graph (Figure 4). This peak corresponds to a fluorescence intensity even lower than that of the non-fluorescent spores produced after meiosis of hemizygous diploids. We checked that this peak corresponds to spores produced by non-fluorescent diploid cells. This suggests that the fluorescent proteins are not completely degraded during meiosis, and are still found in the cytomplasm of spores which do not have the marker gene. We observed this behaviour with our tester strains in which the fluorescent marker genes are placed under a strong ubiquitous TDH3 promoter, but it should be different when using a spore-specific promoter as proposed by (Thacker et al. 2011b). The possibility to select spores descending from non-fluorescent diploids, and to discard them for the analysis is very helpful when such spores are present in the sample, because measuring recombination can be done only with spores produced by hemizygous diploids. In such cases, CAYSS discards spores belonging to the left peak for each marker. Interestingly, when that peak is less clearly separated from the middle peak for one of the markers (e.g. for yECerulean in Figure 4), the other two markers help selecting the adequate subset of spores (produced by hemizygous diploids), so the frequencies estimated even for yECerulean are correct.

**Figure 4.**
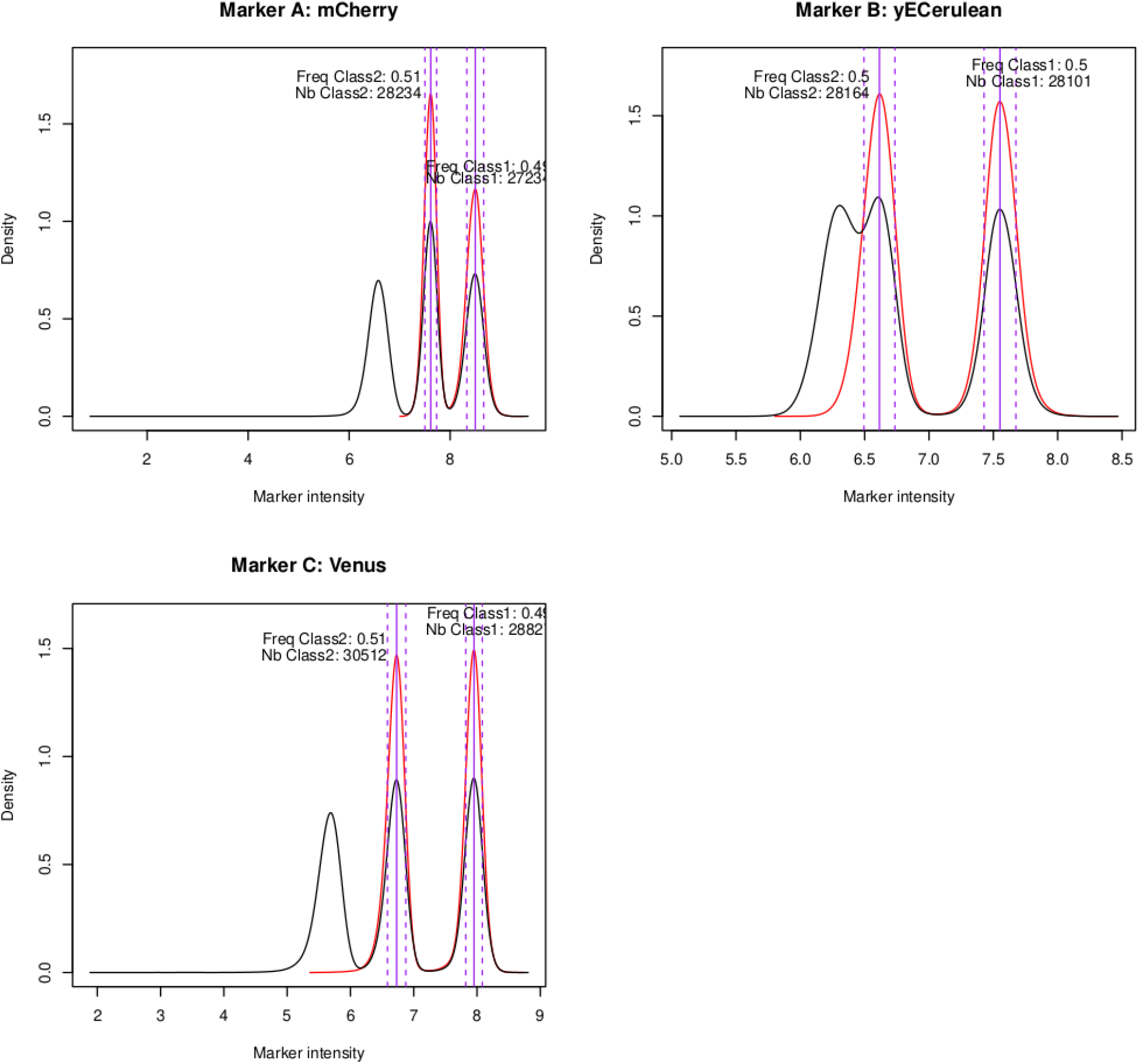
Density (Y-axis) distribution of fluorescence intensity of spores at three markers, in a sample showing three peaks: (1) a peak on the left corresponding to spores which do not produce the fluorescent protein and derive from a non-fluorescent diploid which does not contain any marker gene, (2) a peak in the middle corresponding to spores which do not produce the protein but descend from fluorescent diploids hemizygous for the fluorescent marker genes and which still contain some remainings of fluorescent proteins, and (3) a peak on the right, which corresponds to spores producing the fluorescent proteins. Black curves indicate distributions for all spore singlets. Red curves indicate the distributions for the subset sof spore singlets that belong to the two most right-positionned peaks for all three markers, which corresponds to spores coming from hemizygous diploids (the only ones that may be used to study recombination).

### 5. Maximum-likelihood inference of fluorescence extinction

We observed previously that with some transformed strains, the fluorescent marker shows some probability of extinction of the fluorescence after sporulation. We showed that this is due to a homologous recombination-mediated change in the sequence coding for the fluorescent protein, so one of the markers changes its colour for that of a neighbouring marker (Raffoux et al. 2018a). Using an algorithm already published in that paper, the CAYSS package infers by Maximum-Likelihood the probability of extinction for each of the three markers, as well as the recombination rates in the two intervals, and the CoC. The latter three values are most of the time very close to their naive estimates, except when some of the extinction probabilities are high. In most of the constructs, extinction is very low or even not detectable (e.g. zero value inferred for Aext and Cext in Figure S4 of Supplementary Material 6).

### 6. Comparison with manual analysis

Flow cytometry analyses are usually carried out manually by expert users who draw gates visually around clouds of points in 2-D scatter plots. To assess the consistency of the results automatically produced by CAYSS with our manual analyses, we used CAYSS to re-analyze FCS files from 579 samples that we considered as valid, that is for which recombination rates were lower than 0.5 and, and CoC was lower than 1.3. All these samples had already been analyzed manually to compute recombination rates and CoC (Raffoux et al. 2018b). Figure 5 (top panel) shows that the correlation between automatic and manual values is good, with R^2^ close to 0.99 for recombination rates rAB and rBC in the two intervals between adjacent markers, and 0.93 for the CoC. The distributions of differences between automatic and manual values are shown in the bottom panel of Figure 5. The averages of absolute values of differences between automatic and manual results were 0.0034, 0.0041, and 0.028 for rAB, rBC, and CoC respectively, which relatively to the average of the manual values, corresponds to respectively 2.4%, 2.0%, and 2.7%, showing that automatic results are very close to manual ones. Now, when looking directly at the differences between automatic and manual results instead of their absolute values, the averages for rAB, rBC, and CoC were respectively -0.0002, -0.0005, and -0.016, which, relatively to the average of the manual values, corresponds to respectively -0.3%, -0.1%, and -1.4% differences. Although these values of systematic bias are significantly different from zero (Student t-test, p-value<10^−16^) they are probably very small compared to the intrinsic uncertainty of such cytometry analyses. So we consider that once the parameters have been adapted for each fluorescent marker and flow cytometer, the quality of the analyses done automatically with CAYSS is equivalent to what we did manually.

**Figure 5.**
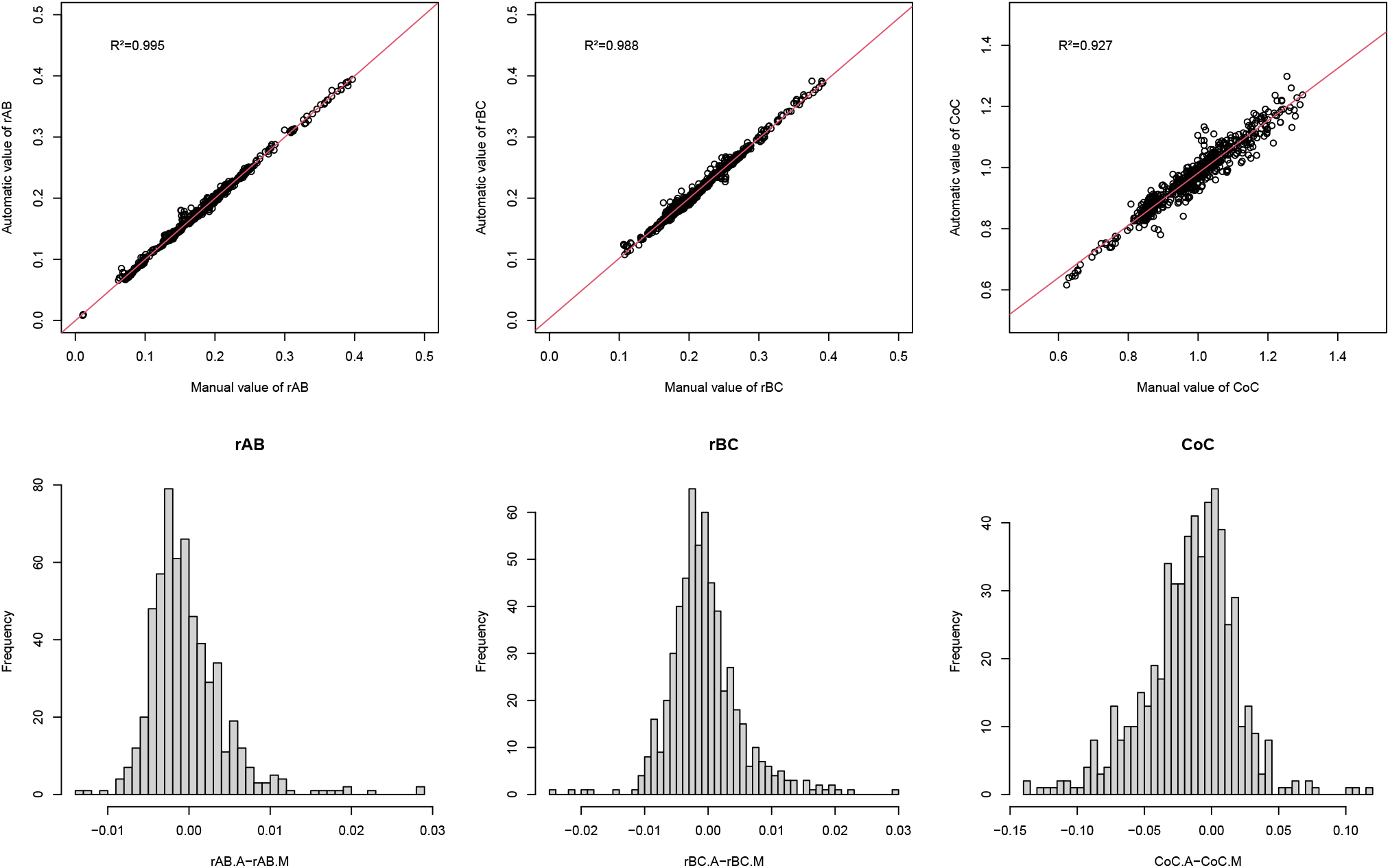
Comparison between manual and automatic recombination analysis for 579 samples, each corresponding to spores from a different cross from (Raffoux et al. 2018b). The recombination parameters were first determined by manual analysis and visual gating using the software ‘Summit’, and then the same FCS files were re-analyzed using CAYSS with the default parameters. The values of rAB.M, rBC.M, and CoC.M indicate respectively the manual estimations of recombination rates in the first and second interval between adjacent markers, and of the coefficient of coincidence. The values of rAB.A, rBC.A, and CoC.A are the corresponding parameters automatically computed with CAYSS. Top panel: correlation between automatic and manual values. Bottom panel: distribution of differences between automatic and manual values.

### 7. Conclusions

Flow cytometry is a very efficient tool for high-throughput random spore-based analysis of recombination in yeast, but the time required for manual gating of raw fluorescence data is very significant. Moreover, because of the unavoidable subjectivity of such analyses, they should be performed always by the same person to allow faithful comparisons, and even though, it is very difficult to keep exactly the same analysis criteria over long periods of time. While being fully automatic and reproducible, CAYSS faithfully mimmics the manual analysis process that we developed over years while analyzing thousands of samples, among which some from very diverse strains (Raffoux et al. 2018b), and the results are very close to those obtained manually for the same raw data. CAYSS is freely available under GPL licence, so we hope it can be useful to other research groups, as it is to us.

## Supporting information

CAYSS tutorial (PDF)

CAYSS R package

## ACKNOWLEDGEMENTS

Authors contributions: XR developed part of the R code and carried out all wet-lab experiments as well as manual cytometry data analyses. MF developed part of the R code, re-analyzed previous manually-processed cytometry data with CAYSS, and wrote the manuscript. Both authors read and approved the manuscript.

The authors thank Mickael Bourge for many years of sharing with us his valuable experience in cytometry analysis, Franck Gauthier for his help to check the R package, and both for their advice on the manuscript. The present work has benefited from the Cytometry / Electronic Microscopy / Light Microscopy facility of Imagerie-Gif, (http://www.i2bc.paris-saclay.fr), member of IBiSA (http://www.ibisa.net), supported by “France-BioImaging” (ANR-10-INBS-04-01), and the Labex “Saclay Plant Science” (ANR-11-IDEX-0003-02). The present work has also benefited from funding from the EVOLREC (ANR-20-CE13-0010) and CO-PATT (ANR-20-CE12-0006) projects. GQE—Le Moulon benefits from the support of the LabEx Saclay Plant Sciences-SPS (ANR-10-LABX-0040-SPS).

## Notes

https://forgemia.inra.fr/gqe-base/cayss/-/releases

### Competing Interest Statement

The authors have declared no competing interest.

