## Supplementary material for "CAYSS: package for automatic Cytometry Analysis of Yeast Spore Segregation": CAYSS tutorial (PDF)

### Tutorial of the CAYSS R Package

2024-09-26

This tutorial does not present all possibilities of the CAYSS package. It only shows the most standard examples of use to compute recombination rates and crossover interference from raw cytometry data.

For a complete list of arguments and details about the algorithms, rather use the package documentation and type '? AnalyzeRecombination'

#### 1. Two-markers data set example of analysis

Since there are only two markers, only recombination rate rAB can be computed. Crossover interference cannot be analyzed, and Maximum-Likelihood estimations of fluorescence extinction cannot be performed either. Thus, a lot of columns in the res output file will contain NA.

```
library(CAYSS)
exampleFolder <- CAYSS::ExampleFile(type="exampleFolder")
plateDesignFileName1 <- CAYSS::ExampleFile("PlateDesign1")
read.table(plateDesignFileName1, header=TRUE)
```

```
##                   Sample fcs_ID
## 1 Exp1_20230118_VI_C1Y2     C1
```

```
res <- AnalyzeRecombination(
            outDir=exampleFolder, # or outputFolder <- file.path("/xxx/yyy")
            fcsDir=exampleFolder,
            plateDesignFile=plateDesignFileName1,
            tokensDateChrMarkers=c(2,3,4),
            pdfGraphs="none" # plots graphs in a window and not in a pdf file
       )
```

```
## Exp1_20230118_VI_C1Y2_C1 => Exp1_C1.fcs
## Type of cytometer detected: CytoFLEX after 2021
## Removed 7783 duplicated event IDs (TIME)
## Removed 28568 invalid raw data in channels
## Valid events imported: 348541
```

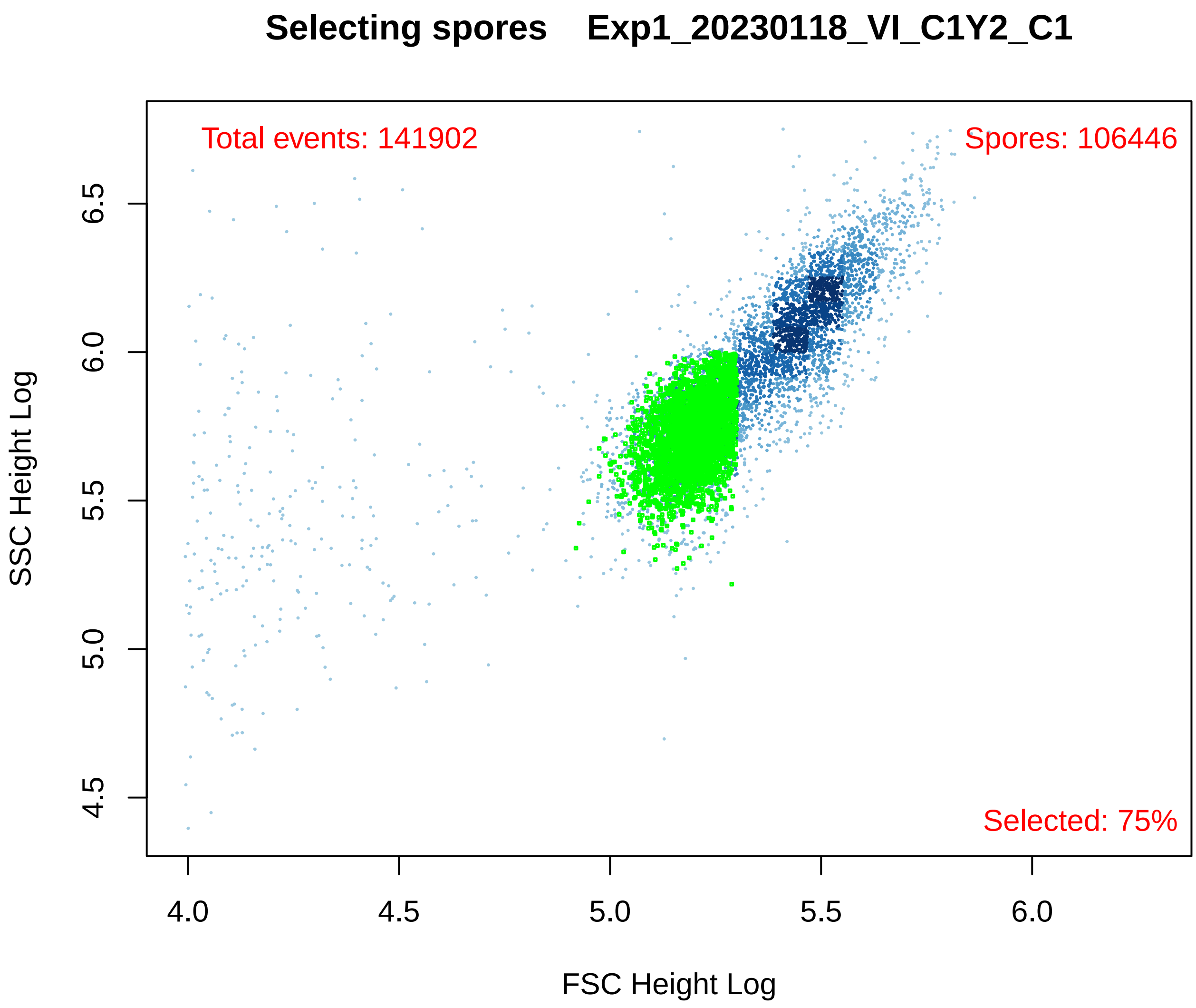
Selecting spores Exp1_20230118_VI_C1Y2_C1
Total events: 141902
Spores: 106446
Selected: 75%
SSC Height Log
FSC Height Log
4.5
5.0
5.5
6.0
6.5
4.0
4.5
5.0
5.5
6.0

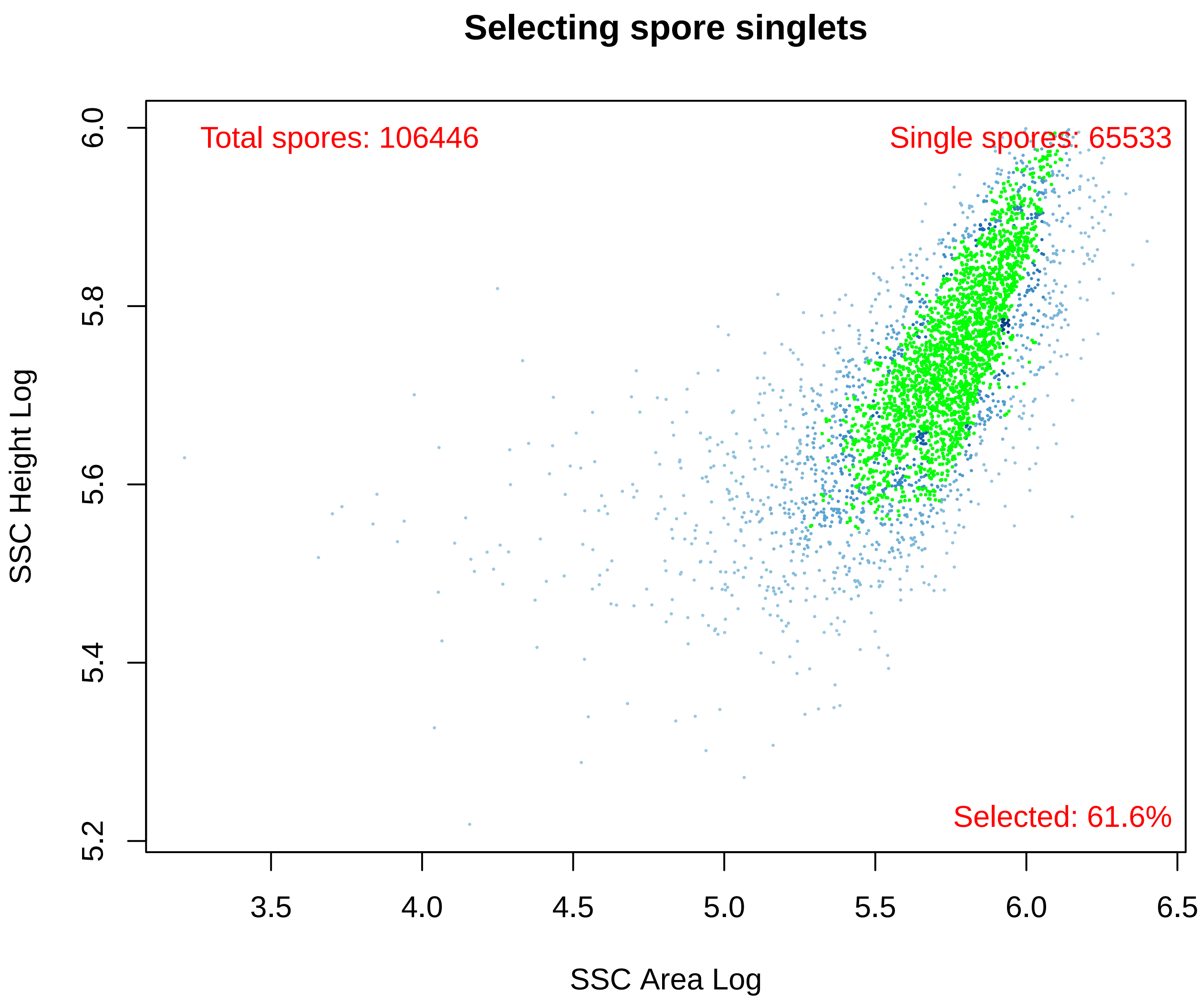
Selecting spore singlets
Total spores: 106446
Single spores: 65533
Selected: 61.6%
SSC Height Log
6.0
5.8
5.6
5.4
5.2
3.5
4.0
4.5
5.0
5.5
6.0
6.5
SSC Area Log

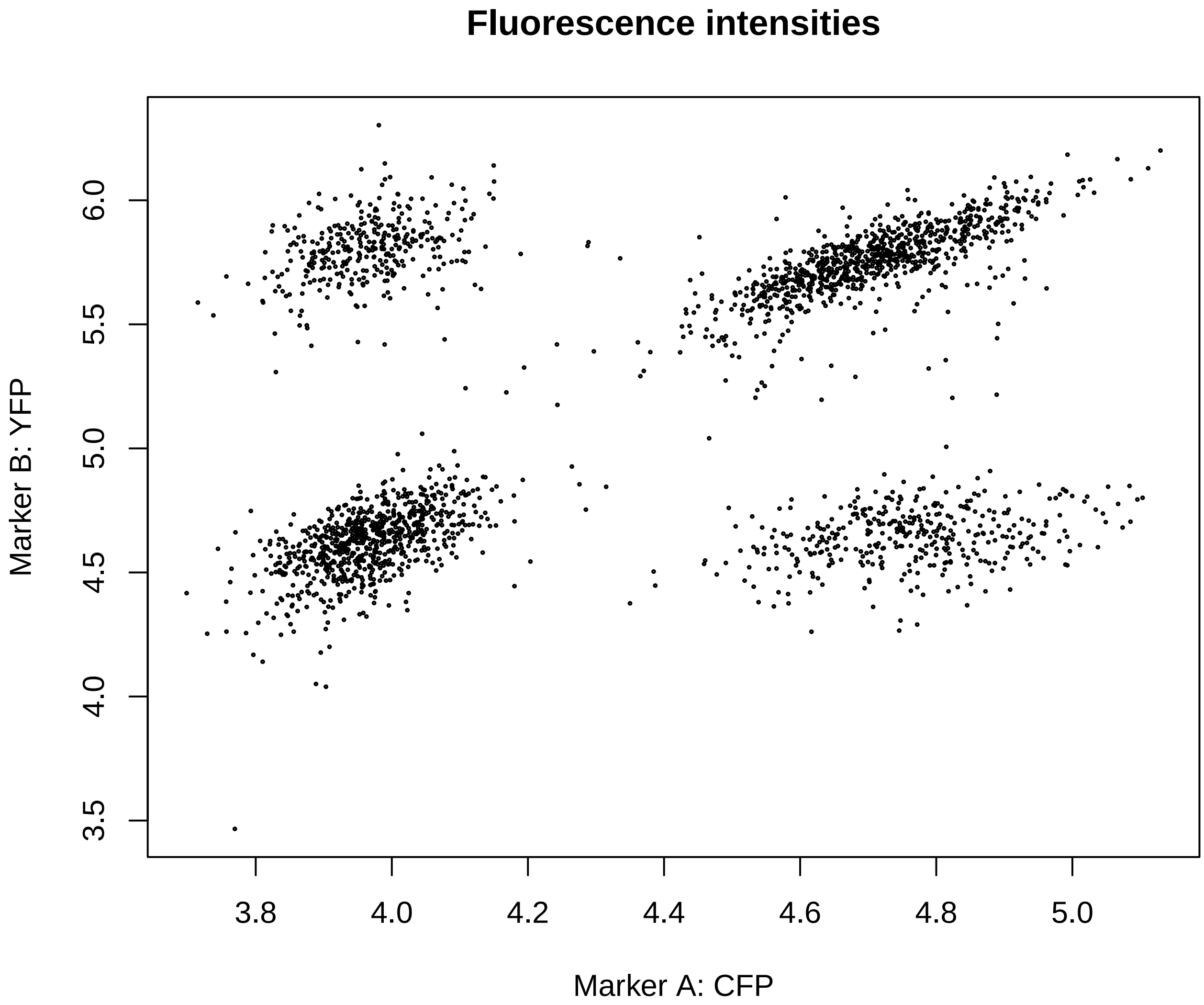


```
## Marker: CFPHeightLog
```

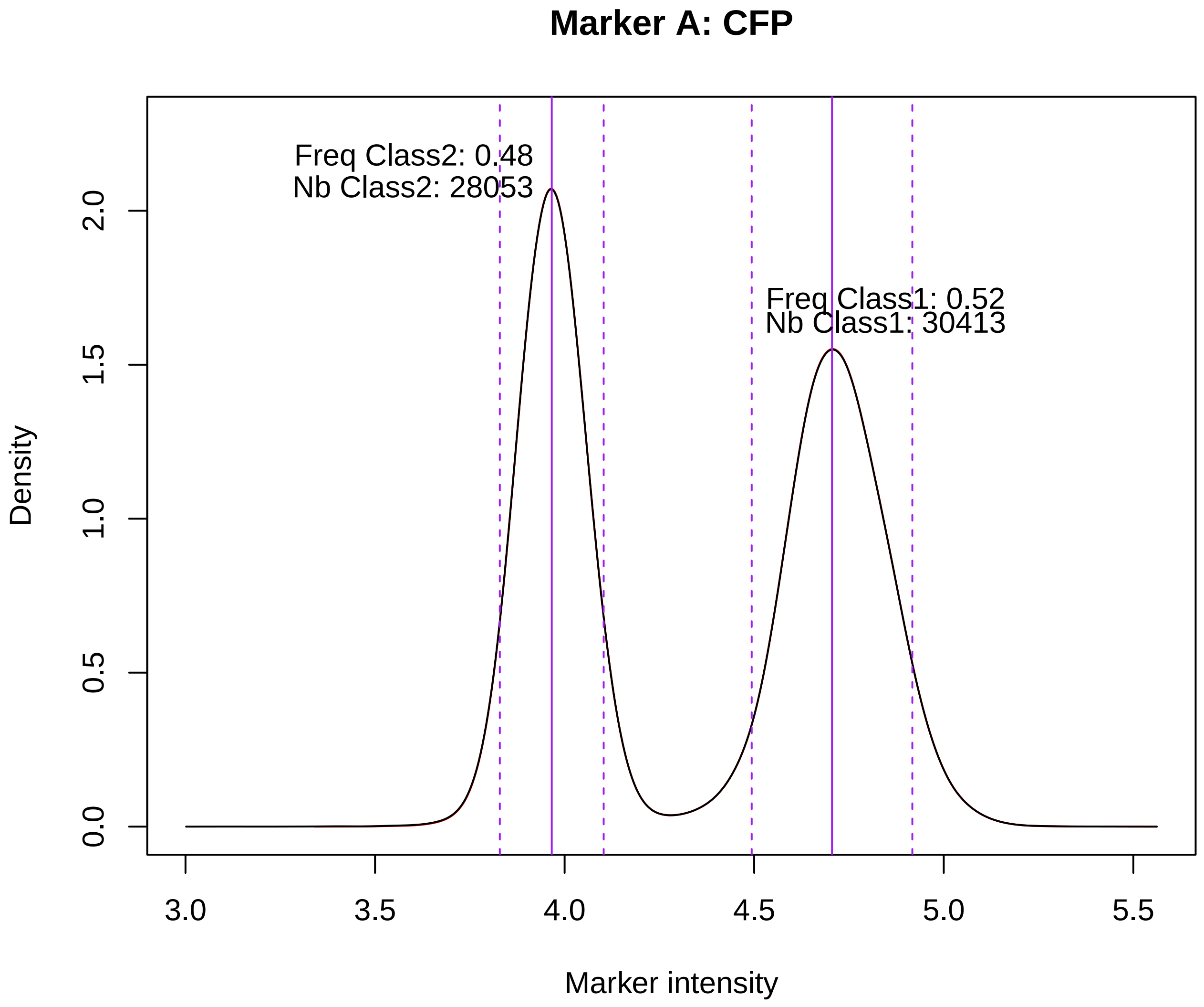


```
## Marker: YFPHeightLog
```

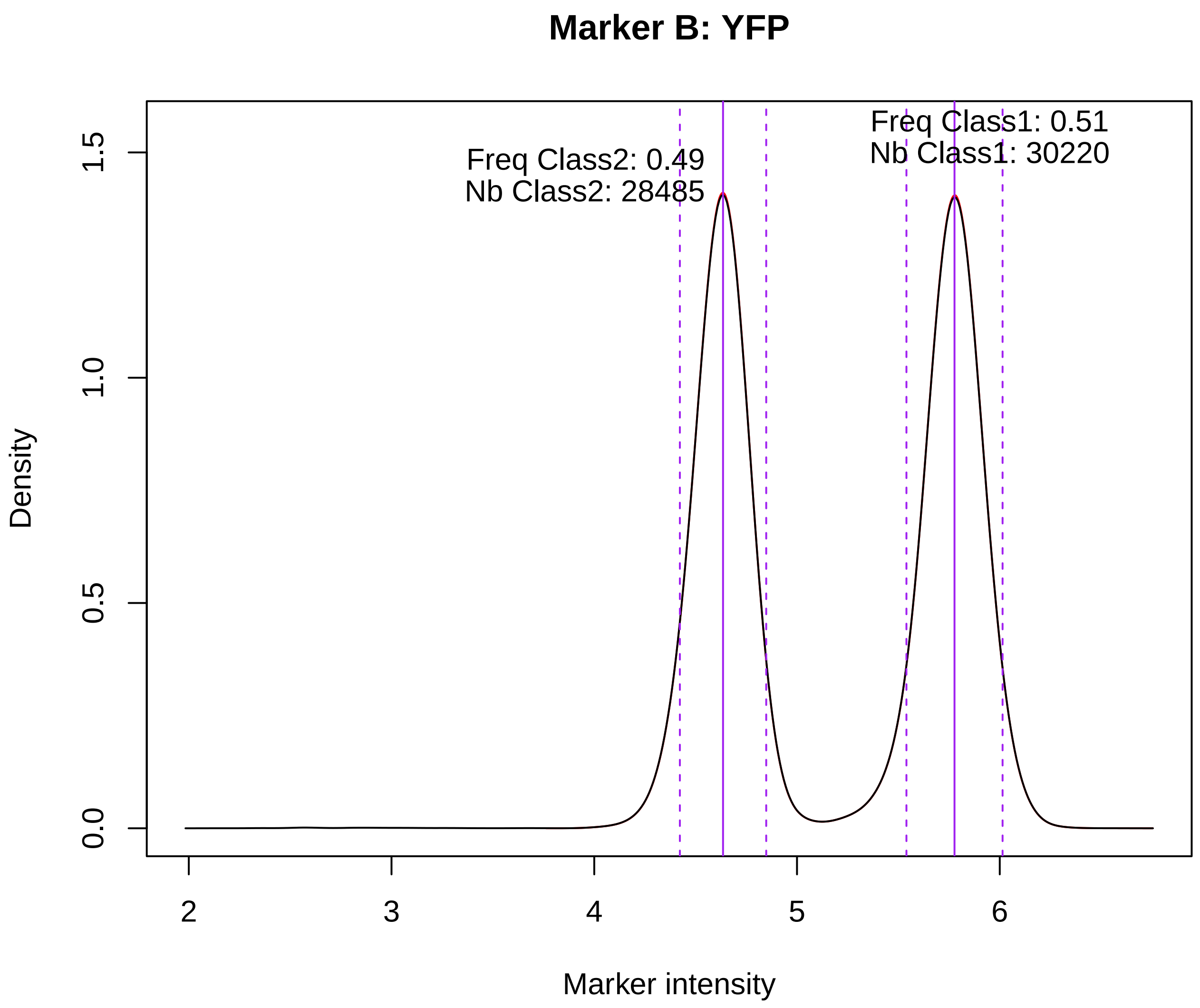


#### Computing naive recombination rates for 2 markers

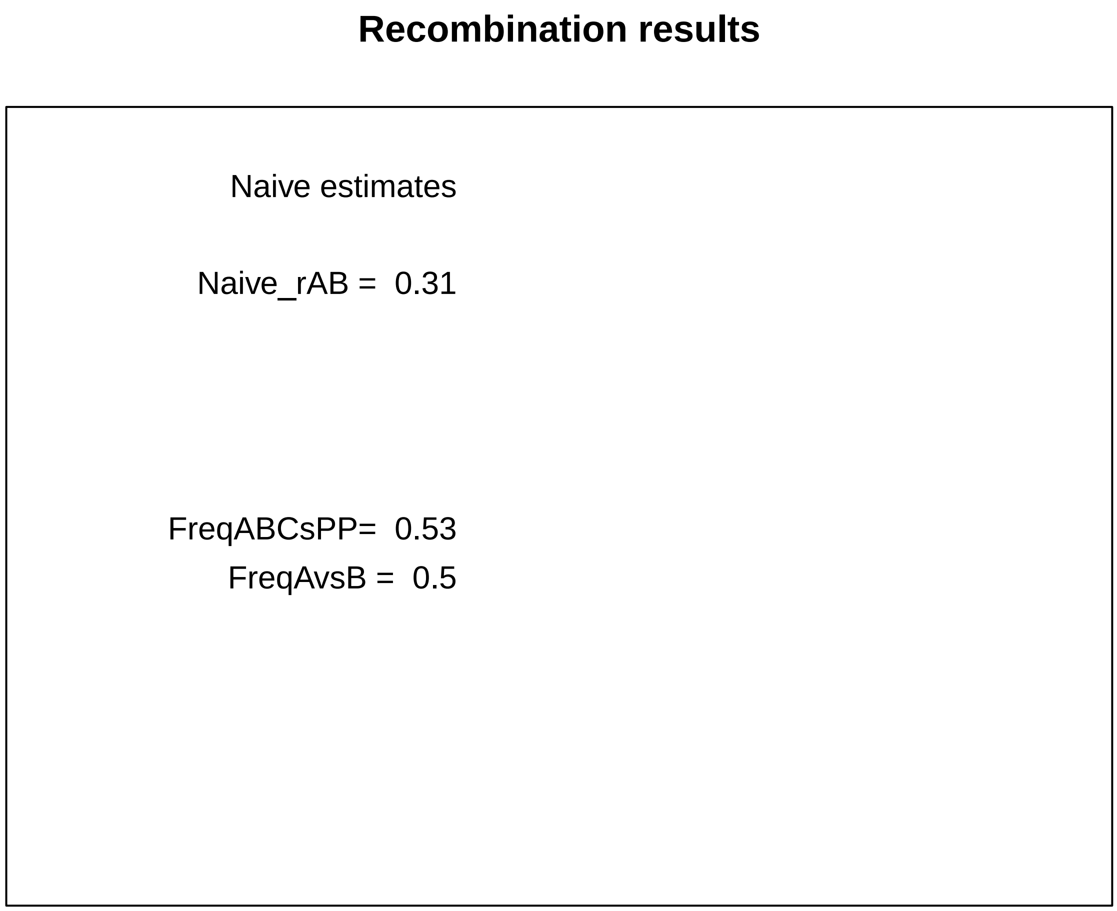


All numerical results in 'res' are also written in file YYYY-MM-DD_hhmmss_ALL_RESULTS.txt located in folder YYYY-MM-DD_hhmmss_RESULTS. For details about the columns of this table, use '? AnalyzeRecombination'

```
print(res)  # prints the numerical results that are written in the RESULTS output folder
```

```
##                  SampleName    DateExp Chr Markers NbEvents NbPreselected
## 1 Exp1_20230118_VI_C1Y2_C1 2023-01-18  VI    C1Y2   348541        141902
##   NbSpores NbSingletSpores Freq_CFP Freq_RFP Freq_YFP FreqABCvsPPP FreqBCvsA
## 1   106446           65533     0.52       NA     0.51           NA        NA
##   FreqABvsC FreqACvsB Naive_rAB Naive_rBC Naive_rDouble Naive_CoC StatusFcs
## 1        NA        NA 0.3050317        NA            NA        NA        OK
##     StatusGating  App   ABp ABC ApC  pBp pBC ppC   ppp TotSpores ML_rAB ML_rBC
## 1 Data number OK 8273 20048  NA  NA 8210  NA  NA 17506        NA     NA     NA
##   ML_CoC ML_Aext ML_Bext ML_Cext StatusML ChangedParams
## 1     NA      NA      NA      NA       NA            NA
```

#### 2. Three-markers data set example of analysis

Since there are three markers (A,B,C in the order of their position on the chromosome), recombination rates rAB and rBC can be computed, as well as the coefficient of coincidence (CoC). In addition, Maximum-Likelihood analysis will be performed to take into account possible fluorescence extinction (see Raffoux et al., 2018, Yeast 35:731)

```
library(CAYSS)
exampleFolder <- CAYSS::ExampleFile(type="exampleFolder")
```

```r
plateDesignFileName2 <- CAYSS::ExampleFile("PlateDesign2")
read.table(plateDesignFileName2, header=TRUE)
```

```
##                     Sample fcs_ID
## 1 Exp2_20230321_VI_R3Y4C5     E1
```

```r
res <- AnalyzeRecombination(
  outDir=exampleFolder, # or outputFolder <- file.path("/xxx/yyy")
  fcsDir=exampleFolder,
  plateDesignFile=plateDesignFileName2,
  tokensDateChrMarkers=c(2,3,4),
  pdfGraphs="none" # plots graphs in a window and not in a pdf file
)
```

```
## Exp2_20230321_VI_R3Y4C5_E1 => Exp2_E1.fcs
## Type of cytometer detected: CytoFLEX after 2021
## Removed 6730 duplicated event IDs (TIME)
## Removed 4112 invalid raw data in channels
## Valid events imported: 375815
```

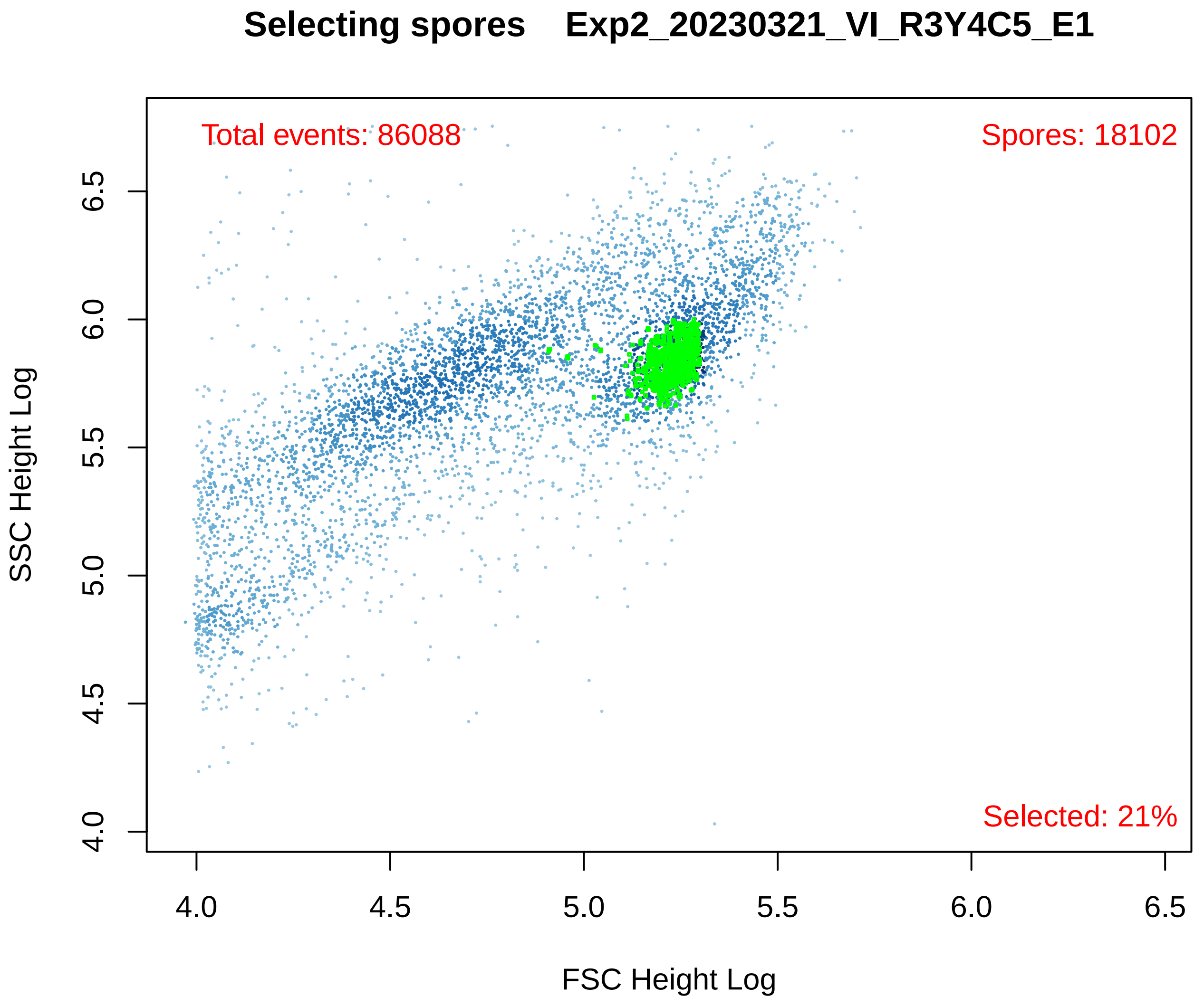

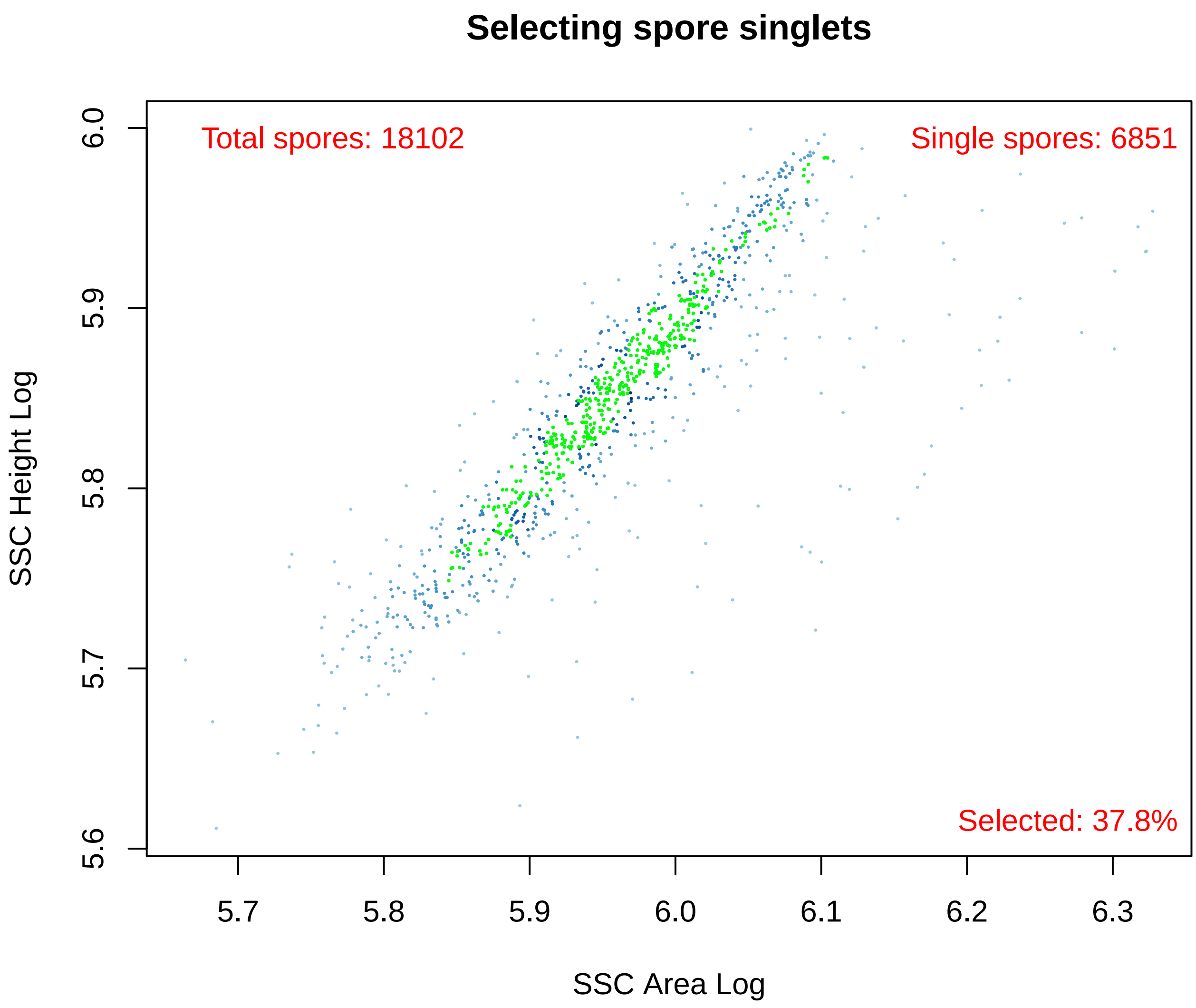


```
## Marker: RFPHeightLog
```

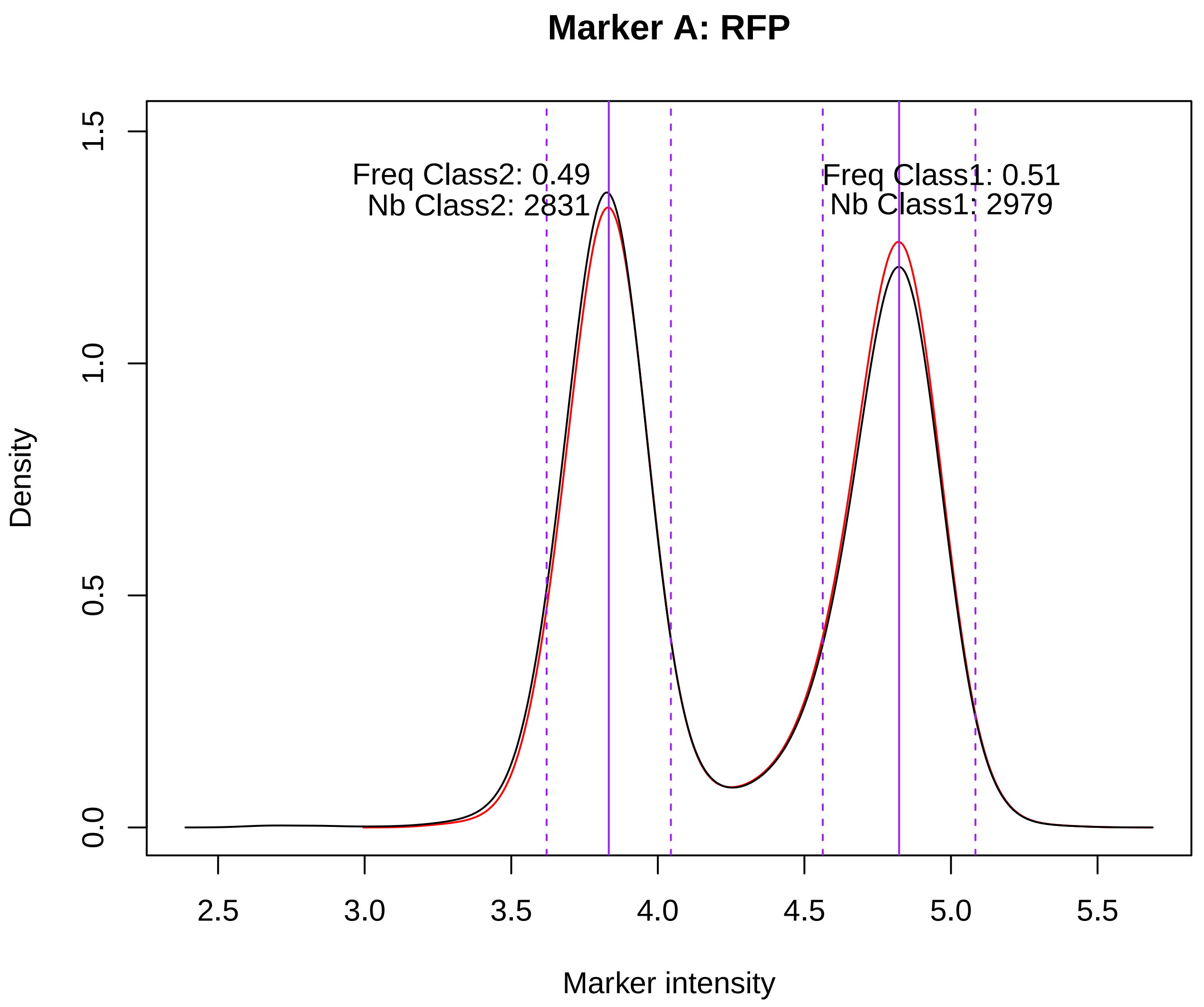


```
## Marker: YFPHeightLog
```

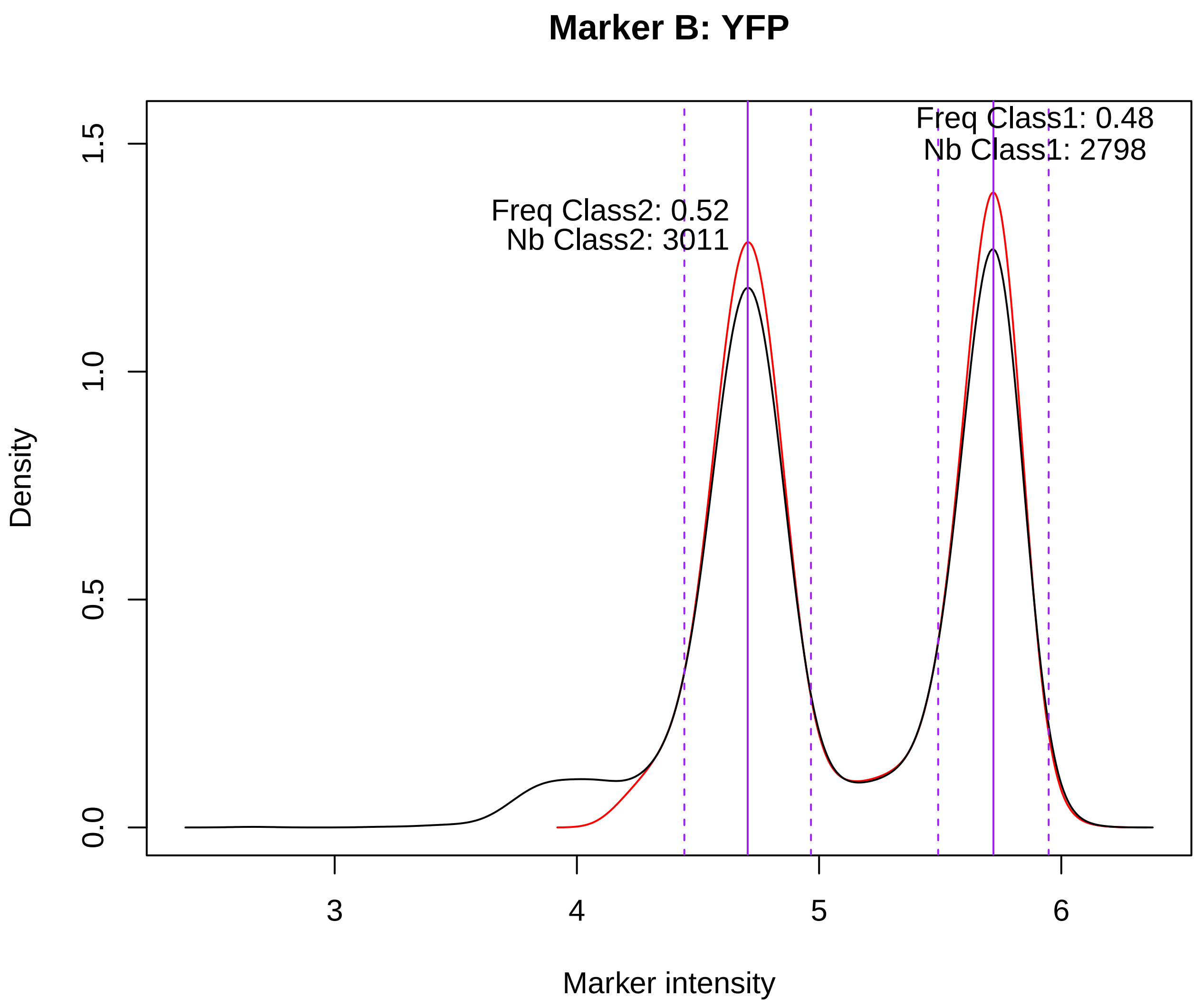


```
## Marker: CFPHeightLog
```

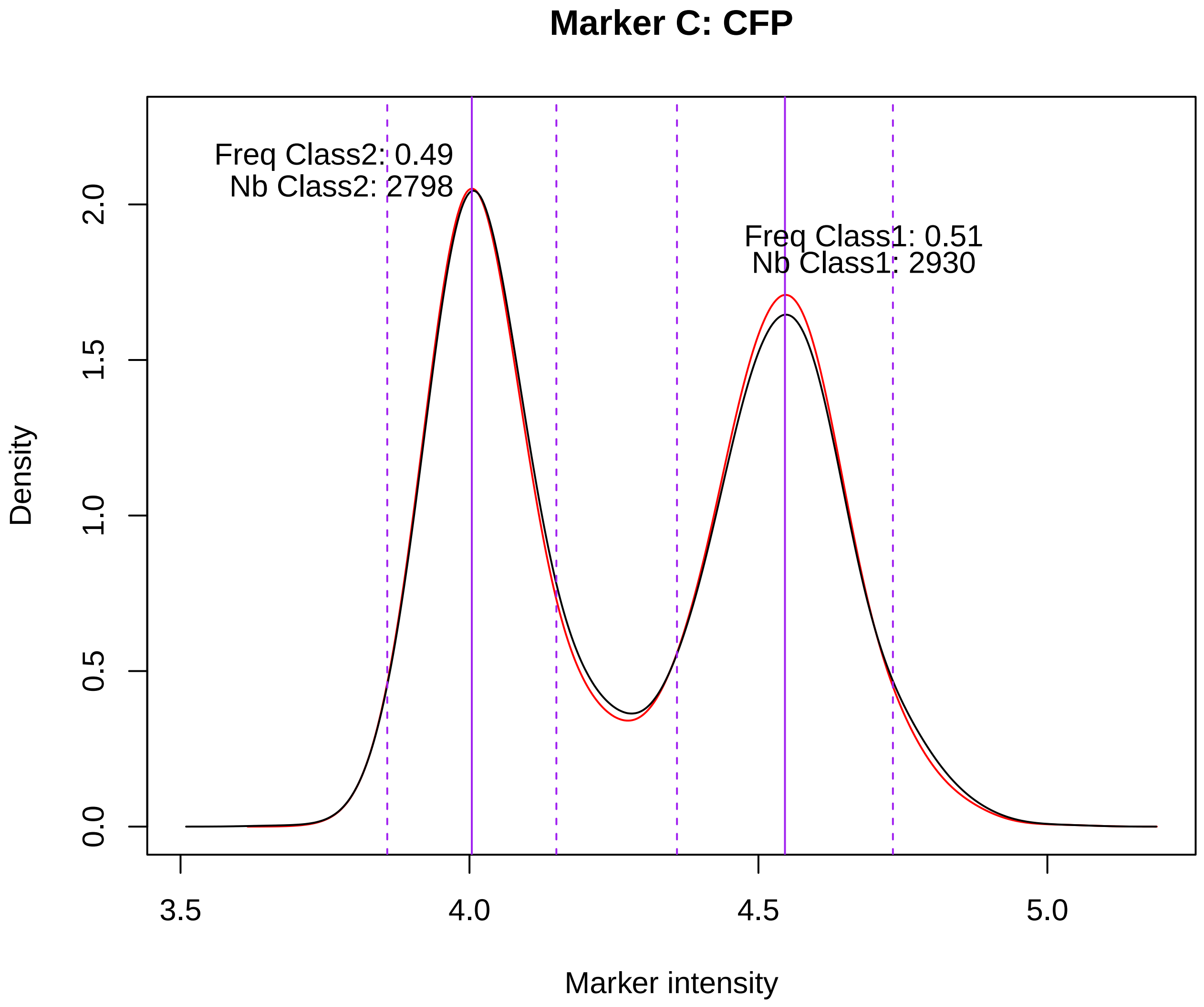


```
## Computing naive recombination rates for 3 markers

## Computing Max Likelihood recombination rates with 3 markers
```

#### Recombination results

| Naive estimates | ML estimates |
|---|---|
| Naive_rAB: 0.12 | ML_Naive_rAB: 0.11 |
| Naive_rBC: 0.39 | ML_Naive_rBC: 0.38 |
| Naive_CoC: 1.02 | ML_CoC: 0.91 |
| DoubleRec: 0.05 | |
| FreqABCvsPPP: 0.52 | ML_extA: NA |
| FreqBCvsA: 0.45 | ML_extB: 0.04 |
| FreqABvsC: 0.48 | ML_extC: NA |
| FreqACvsB: 0.66 | |

```
##
## Warning: Naive_rAB=0.123832724319935 but ML_rAB=0.105780363903273
## Warning: Naive_CoC=1.01790880286541 but ML_CoC=0.909195675990886
```

All numerical results in 'res' are also written in file YYYY-MM-DD_hhmmss_ALL_RESULTS.txt located in folder YYYY-MM-DD_hhmmss_RESULTS For details about the columns of this table, use '?AnalyzeRecombination'

```
print(res)  # prints the numerical results that are written in the RESULTS output folder
```

```
##                    SampleName    DateExp Chr Markers NbEvents NbPreselected
## 1 Exp2_20230321_VI_R3Y4C5_E1 2023-03-21  VI  R3Y4C5   375815         86088
##   NbSpores NbSingletSpores Freq_CFP Freq_RFP Freq_YFP FreqABCvsPPP FreqBCvsA
## 1    18102            6851     0.51     0.51     0.48    0.5169683 0.4486486
##   FreqABvsC FreqACvsB Naive_rAB Naive_rBC Naive_rDouble Naive_CoC StatusFcs
## 1 0.4777644    0.6625 0.1238327 0.3865205    0.04872107  1.017909        OK
##     StatusGating App ABp  ABC ApC pBp pBC ppC  ppp TotSpores    ML_rAB
## 1 Data number OK 204 795 1371 159  81 166 869 1281      4926 0.1057804
##      ML_rBC    ML_CoC ML_Aext    ML_Bext ML_Cext
## 1 0.3802785 0.9091957      NA 0.04428487      NA
##                           StatusML ChangedParams
## 1 Warning: Naive-ML difference > 10%            NA
```

#### 3. Complex fluorescence patterns.

Example of spores from a mixture of diploids without fluorescent marker and diploids hemizygous for three markers. In that case, three peaks of fluorescence intensity are observed, corresponding (from left to right on the plots) to:

- spores which come from non-fluorescent diploids, and which do not express fluorescence
- spores which come from fluorescent (hemizygous) diploids, and which do not express fluorescence
- spores which come from fluorescent (hemizygous) diploids, and which do express fluorescence

```
library(CAYSS)
exampleFolder <- CAYSS::ExampleFile(type="exampleFolder")
plateDesignFileName3 <- CAYSS::ExampleFile("PlateDesign3")
read.table(plateDesignFileName3, header=TRUE)
```

```
##                    Sample fcs_ID
## 1 Exp3_20211027_I_R2C3Y4     G7
```

```
res <- AnalyzeRecombination(
  outDir=exampleFolder, # or outputFolder <- file.path("/xxx/yyy")
  fcsDir=exampleFolder,
  plateDesignFile=plateDesignFileName3,
  tokensDateChrMarkers=c(2,3,4),
  pdfGraphs="none" # plots graphs in a window and not in a pdf file
)
```

```
## Exp3_20211027_I_R2C3Y4_G7 => Exp3_G7.fcs
## Type of cytometer detected: CytoFLEX before 2021
## Removed 7116 duplicated event IDs (TIME)
## Removed 69 invalid raw data in channels
## Valid events imported: 246188
```

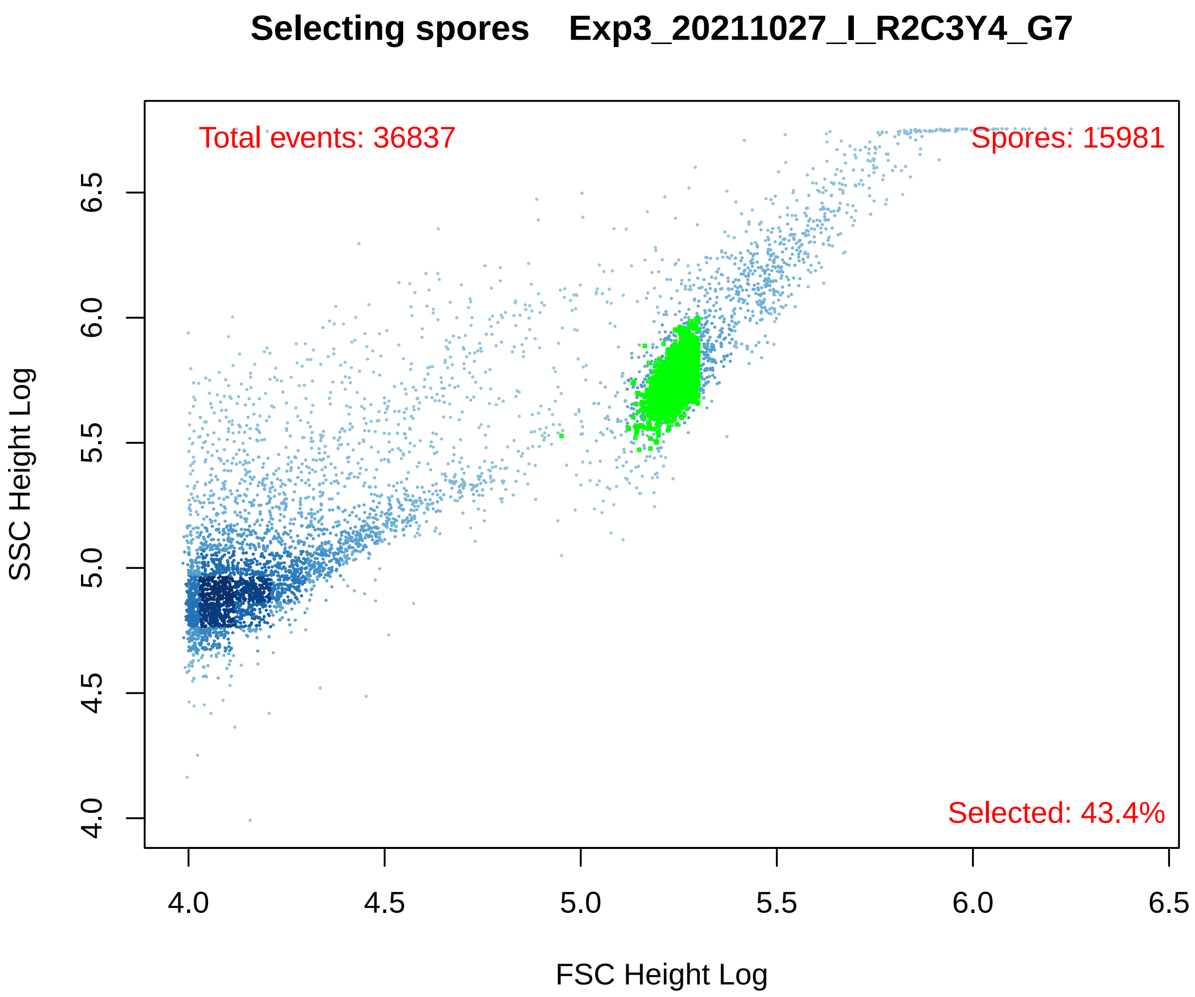
Selecting spores Exp3_20211027_I_R2C3Y4_G7
Total events: 36837
Spores: 15981
Selected: 43.4%
SSC Height Log
FSC Height Log
4.0
4.5
5.0
5.5
6.0
6.5

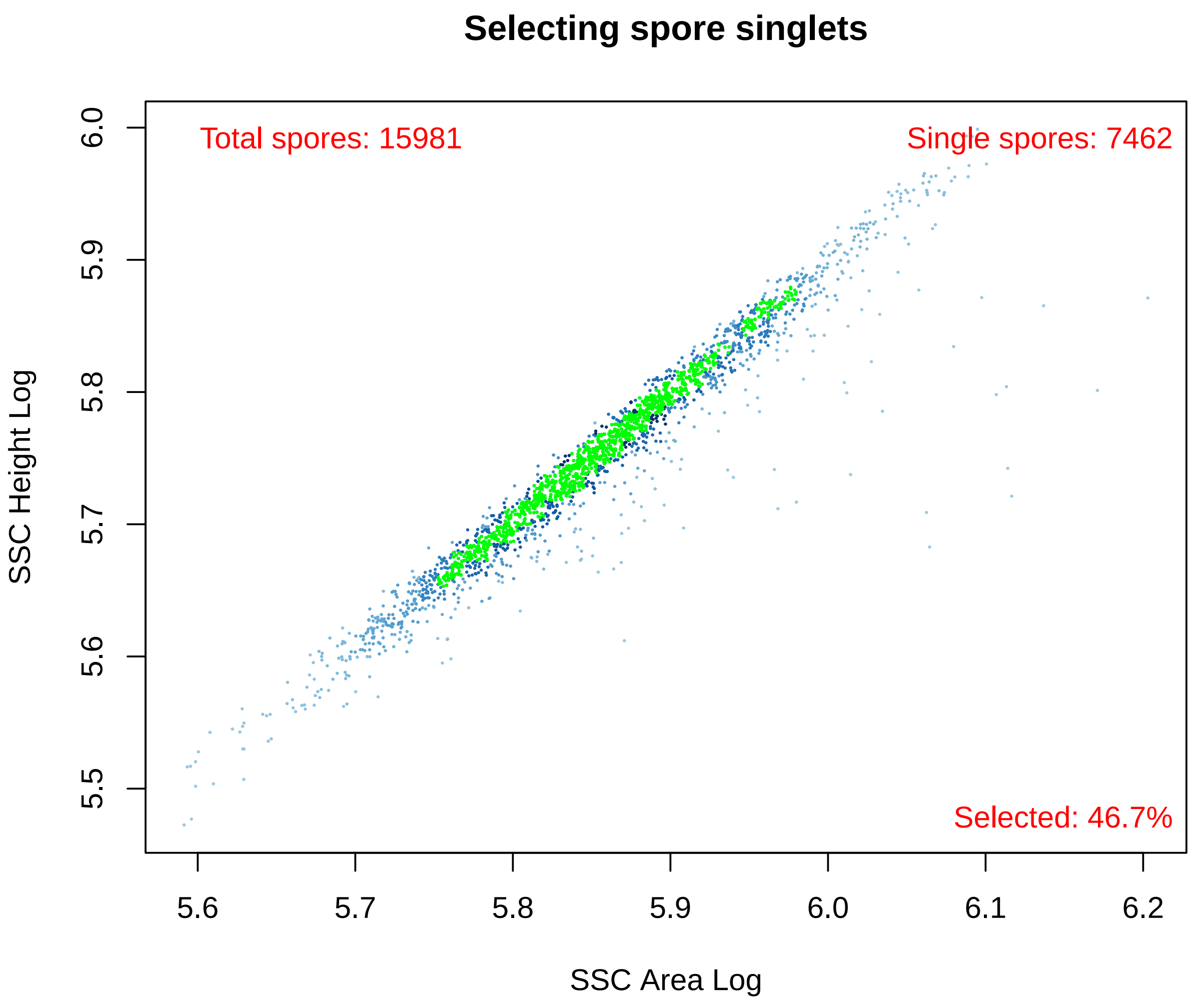


```
## Marker: RFPHeightLog
```

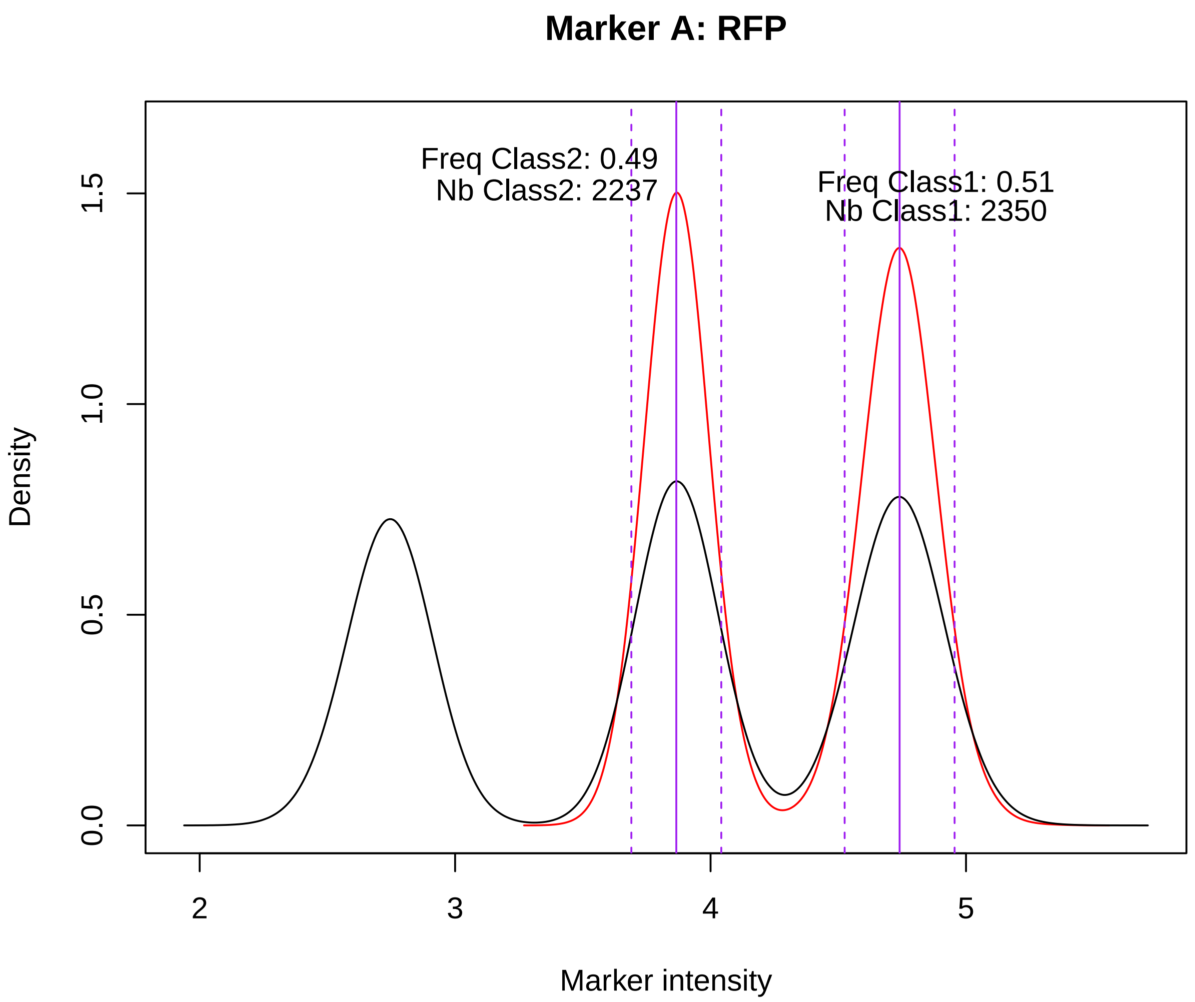


```
## Marker: CFPHeightLog
```

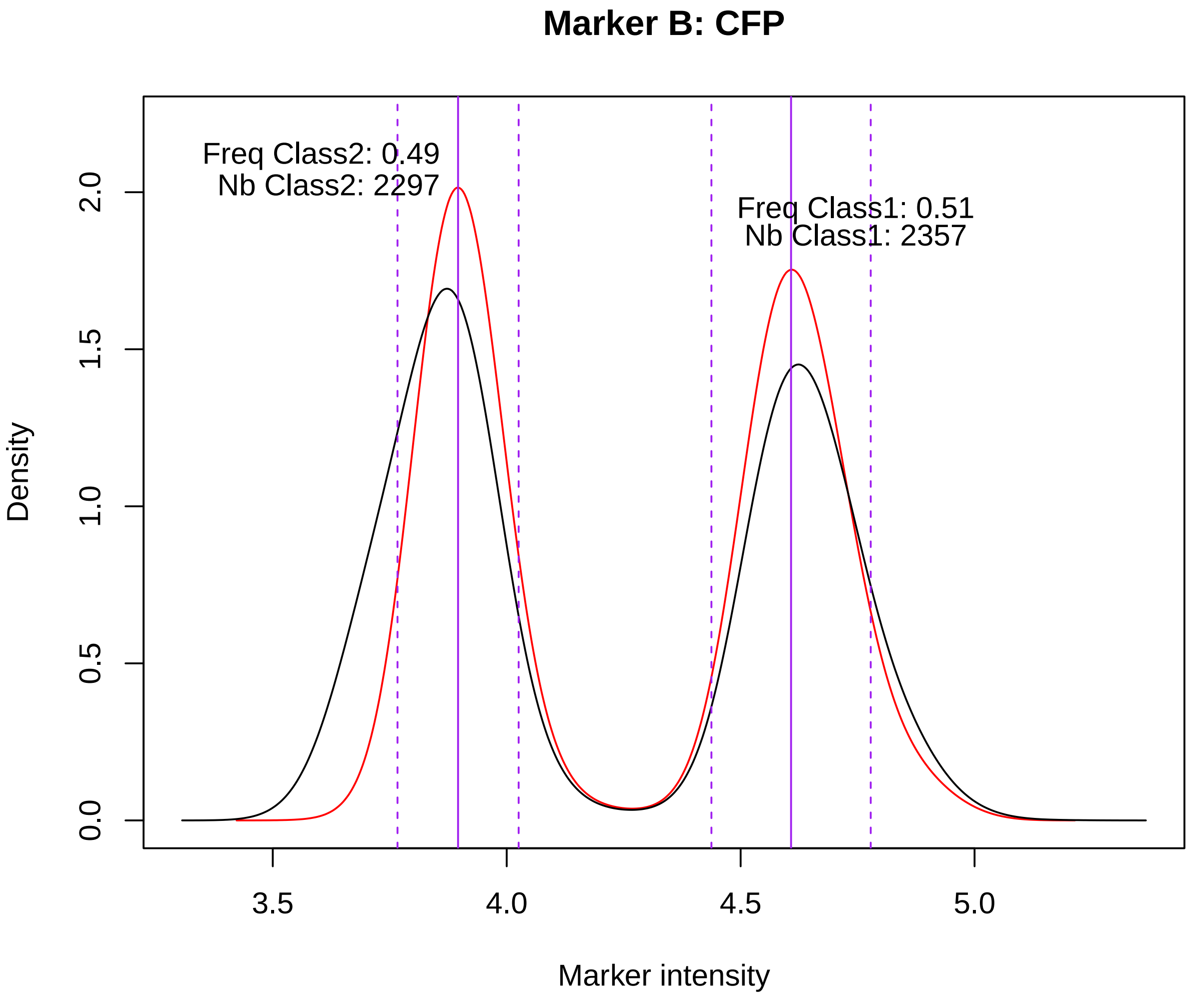


```
## Marker: YFPHeightLog
```

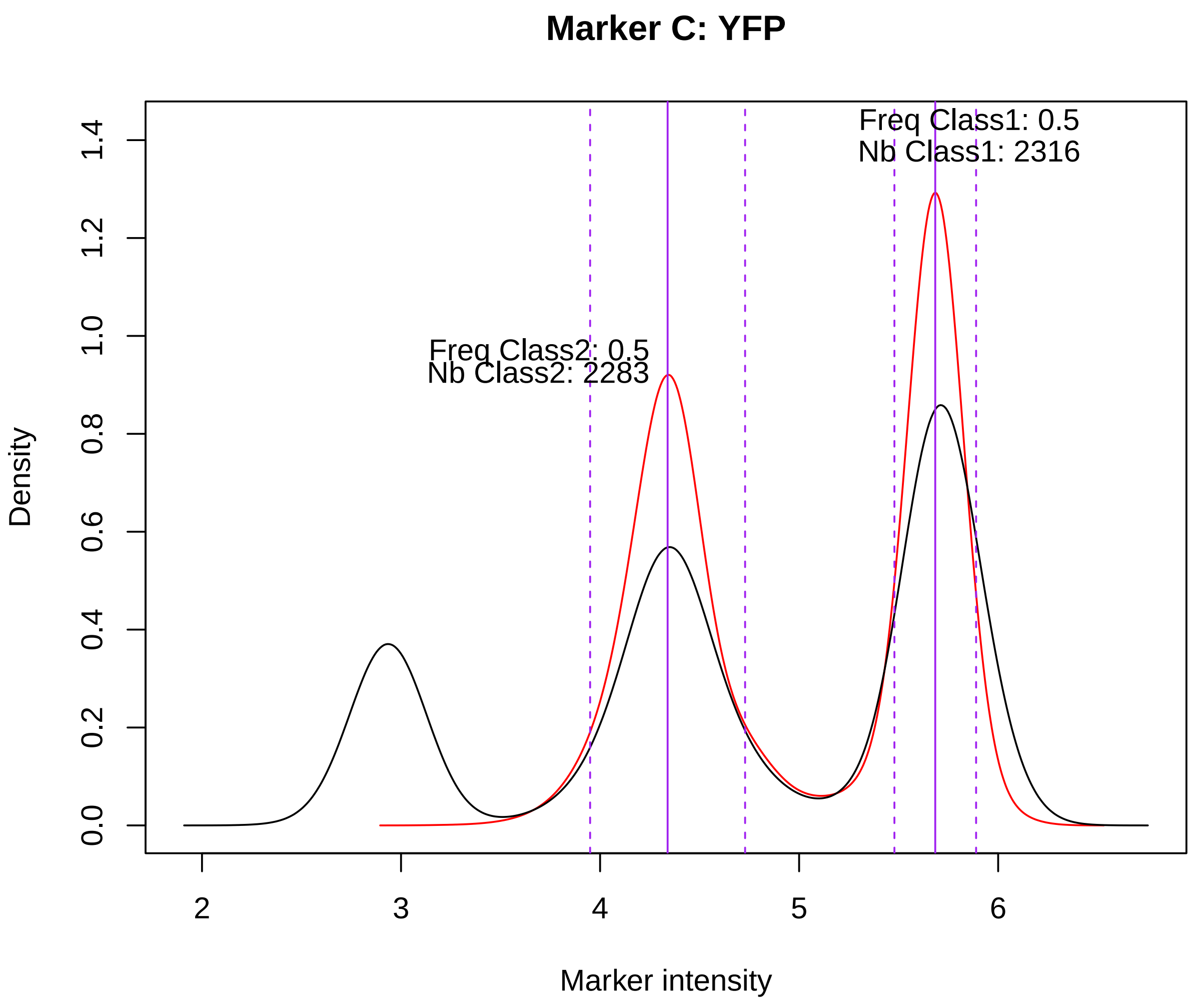


```
## Computing naive recombination rates for 3 markers

## Computing Max Likelihood recombination rates with 3 markers
```

##### Recombination results

| Naive estimates | ML estimates |
|---|---|
| Naive_rAB: 0.2 | ML_Naive_rAB: 0.19 |
| Naive_rBC: 0.29 | ML_Naive_rBC: 0.28 |
| Naive_CoC: 1.04 | ML_CoC: 1.01 |
| DoubleRec: 0.06 | |
| FreqABCvsPPP: 0.52 | ML_extA: NA |
| FreqBCvsA: 0.51 | ML_extB: 0.01 |
| FreqABvsC: 0.52 | ML_extC: NA |
| FreqACvsB: 0.58 | |

```
print(res)  # prints the numerical results that are written in the RESULTS output folder
```

```
##                  SampleName    DateExp Chr Markers NbEvents NbPreselected
## 1 Exp3_20211027_I_R2C3Y4_G7 2021-10-27   I  R2C3Y4   246188         36837
##   NbSpores NbSingletSpores Freq_CFP Freq_RFP Freq_YFP FreqABCvsPPP FreqBCvsA
## 1    15981            7462     0.51     0.51      0.5    0.5230352 0.5121495
##   FreqABvsC FreqACvsB Naive_rAB Naive_rBC Naive_rDouble Naive_CoC StatusFcs
## 1  0.521542 0.5814978 0.1975117 0.2874546    0.05883878  1.036338        OK
##     StatusGating App ABp  ABC ApC pBp pBC ppC  ppp TotSpores    ML_rAB
## 1 Data number OK 261 460 1158 132  95 274 422 1056      3858 0.1938211
##      ML_rBC   ML_CoC ML_Aext    ML_Bext ML_Cext StatusML ChangedParams
## 1 0.2847414 1.008109      NA 0.01095736      NA       Ok            NA
```

### 4. Batch analysis with all examples listed before

This is the usual way to run analyses. All samples listed in the plate Design File will be automatically analyzed, and all plots will be organized in one-page graphical output pdf files (one file per sample). In addition, a multi-page pdf file (one page for each sample) named YYYY-MM-DD_hhmmss_ALL_CYTO_GRAPHS.pdf will be also generated in the YYYY-MM-DD_hhmmss_GRAPHICS folder if the argument 'pdfGraphs' is set to "both". See details with '?AnalyzeRecombination'

```
library(CAYSS)
exampleFolder <- CAYSS::ExampleFile(type="exampleFolder")
plateDesignFileName123 <- CAYSS::ExampleFile("PlateDesign123")
```

```
read.table(plateDesignFileName123, header=TRUE)
```

```
##                    Sample fcs_ID
## 1   Exp1_20230118_VI_C1Y2     C1
## 2 Exp2_20230321_VI_R3Y4C5     E1
## 3  Exp3_20211027_I_R2C3Y4     G7
```

```
res <- AnalyzeRecombination(
  outDir=exampleFolder, # or outputFolder <- file.path("/xxx/yyy")
  fcsDir=exampleFolder,
  plateDesignFile=plateDesignFileName123,
  tokensDateChrMarkers=c(2,3,4),
  pdfGraphs="both" # plots graphs in a window and not in a pdf file
)
```

```
## Exp2_20230321_VI_R3Y4C5_E1 => Exp2_E1.fcs
## Changed param 'minSporeFscHLog' to 5
## Changed param 'widthAroundPeak' to 1
## Type of cytometer detected: CytoFLEX after 2021
## Removed 6730 duplicated event IDs (TIME)
## Removed 4112 invalid raw data in channels
## Valid events imported: 375815

## Warning in KernSmooth::bkde2D(x, bandwidth = bandwidth, gridsize = nbin, : La
## grille de regroupement par classe est trop grossière pour la (faible) largeur
## de fenêtre : essayez en augmentant 'gridsize'

## Marker: RFPHeightLog

## Marker: YFPHeightLog

## Marker: CFPHeightLog

## Computing naive recombination rates for 3 markers

## Computing Max Likelihood recombination rates with 3 markers

##
## Warning: Naive_rAB=0.119809130771215 but ML_rAB=0.107632808711549
##
## Exp3_20211027_I_R2C3Y4_G7 => Exp3_G7.fcs
## Type of cytometer detected: CytoFLEX before 2021
## Removed 7116 duplicated event IDs (TIME)
## Removed 69 invalid raw data in channels
## Valid events imported: 246188

## Marker: RFPHeightLog

## Marker: CFPHeightLog

## Marker: YFPHeightLog

## Computing naive recombination rates for 3 markers

## Computing Max Likelihood recombination rates with 3 markers
```

```
print(res)  # prints the numerical results that are written in the RESULTS output folder
```

```
##                   SampleName    DateExp Chr Markers NbEvents NbPreselected
## 1 Exp2_20230321_VI_R3Y4C5_E1 2023-03-21  VI  R3Y4C5   375815         74033
## 2  Exp3_20211027_I_R2C3Y4_G7 2021-10-27   I  R2C3Y4   246188         35192
```

```
##   NbSpores NbSingletSpores Freq_CFP Freq_RFP Freq_YFP FreqABCvsPPP FreqBCvsA
## 1    42709           17847     0.50     0.52     0.48    0.5549604 0.4676871
## 2    14995            8897     0.51     0.51     0.52    0.5811765 0.5490716
##   FreqABvsC FreqACvsB Naive_rAB Naive_rBC Naive_rDouble Naive_CoC StatusFcs
## 1 0.5041486 0.6627566 0.1198091 0.3703895    0.04397730 0.9910145        OK
## 2 0.5466867 0.6129032 0.1748691 0.2750436    0.04328098 0.8998754        OK
##     StatusGating App  ABp  ABC ApC pBp pBC  ppC  ppp TotSpores    ML_rAB
## 1 Data number OK 313 1276 2383 226 115 275 1255 1911      7754 0.1076328
## 2 Data number OK 170  363  988  76  48 207  301  712      2865 0.1748691
##      ML_rBC    ML_CoC ML_Aext ML_Bext ML_Cext
## 1 0.3655851 0.9036817      NA      NA      NA
## 2 0.2750431 0.8998680      NA      NA      NA
##                             StatusML                      ChangedParams
## 1 Warning: Naive-ML difference > 10% minSporeFscHLog=5.0_widthAroundPeak=1
## 2                                 Ok                               <NA>
```

### 5. Changing specific parameters for some samples

This is the recommended way to optimize analyses if the default parameters are not appropriate for some of the samples. After a first pass of batch analysis, a new Plate Design File named XXXX_PARAMS.txt is generated, with two additional columns. After visual check of the graphical output file, these columns may be manually edited to specify for each sample, if it should be re-run, and which changes should be made in the launching parameters for that specific sample. The other launching arguments will be the same as specified when calling the function AnalyzeRecombination(). The red curves indicate the distributions for the subset of spores that belong to the two most right-positionned peaks, which correspond to the spores coming from hemizygous diploids (the only ones that may be used to study recombination)

```
library(CAYSS)
exampleFolder <- CAYSS::ExampleFile(type="exampleFolder")
plateDesignFileNameParams <- CAYSS::ExampleFile("PlateDesignParams")
read.table(plateDesignFileNameParams, header=TRUE)
```

```
##                     Sample fcs_ID Re_Do                       Changed_Params
## 1   Exp1_20230118_VI_C1Y2     C1     0                                 <NA>
## 2 Exp2_20230321_VI_R3Y4C5     E1     1 minSporeFscHLog=5.0_widthAroundPeak=1
## 3  Exp3_20211027_I_R2C3Y4     G7     1                                 <NA>
```

```
res <- AnalyzeRecombination(
  outDir=exampleFolder, # or outputFolder <- file.path("/xxx/yyy")
  fcsDir=exampleFolder,
  plateDesignFile=plateDesignFileNameParams,
  tokensDateChrMarkers=c(2,3,4),
  pdfGraphs="both" # plots graphs in a window and not in a pdf file
)
```

```
## Exp2_20230321_VI_R3Y4C5_E1 => Exp2_E1.fcs
## Changed param 'minSporeFscHLog' to 5
## Changed param 'widthAroundPeak' to 1
## Type of cytometer detected: CytoFLEX after 2021
## Removed 6730 duplicated event IDs (TIME)
## Removed 4112 invalid raw data in channels
## Valid events imported: 375815
```

```
## Warning in KernSmooth::bkde2D(x, bandwidth = bandwidth, gridsize = nbin, : La
```

```
## grille de regroupement par classe est trop grossière pour la (faible) largeur
## de fenêtre : essayez en augmentant 'gridsize'

## Marker: RFPHeightLog

## Marker: YFPHeightLog

## Marker: CFPHeightLog

## Computing naive recombination rates for 3 markers

## Computing Max Likelihood recombination rates with 3 markers

##
## Warning: Naive_rAB=0.119809130771215 but ML_rAB=0.107632808711549
##
## Exp3_20211027_I_R2C3Y4_G7 => Exp3_G7.fcs
## Type of cytometer detected: CytoFLEX before 2021
## Removed 7116 duplicated event IDs (TIME)
## Removed 69 invalid raw data in channels
## Valid events imported: 246188

## Marker: RFPHeightLog

## Marker: CFPHeightLog

## Marker: YFPHeightLog

## Computing naive recombination rates for 3 markers

## Computing Max Likelihood recombination rates with 3 markers
```

```r
print(res)  # prints the numerical results that are written in the RESULTS output folder
```

```
##                  SampleName    DateExp Chr Markers NbEvents NbPreselected
## 1 Exp2_20230321_VI_R3Y4C5_E1 2023-03-21  VI  R3Y4C5   375815         74033
## 2  Exp3_20211027_I_R2C3Y4_G7 2021-10-27   I  R2C3Y4   246188         35192
##   NbSpores NbSingletSpores Freq_CFP Freq_RFP Freq_YFP FreqABCvsPPP FreqBCvsA
## 1    42709           17847     0.50     0.52     0.48    0.5549604 0.4676871
## 2    14995            8897     0.51     0.51     0.52    0.5811765 0.5490716
##   FreqABvsC FreqACvsB Naive_rAB Naive_rBC Naive_rDouble Naive_CoC StatusFcs
## 1 0.5041486 0.6627566 0.1198091 0.3703895    0.04397730 0.9910145        OK
## 2 0.5466867 0.6129032 0.1748691 0.2750436    0.04328098 0.8998754        OK
##     StatusGating App  ABp  ABC ApC pBp pBC  ppC  ppp TotSpores    ML_rAB
## 1 Data number OK 313 1276 2383 226 115 275 1255 1911      7754 0.1076328
## 2 Data number OK 170  363  988  76  48 207  301  712      2865 0.1748691
##      ML_rBC    ML_CoC ML_Aext ML_Bext ML_Cext
## 1 0.3655851 0.9036817      NA      NA      NA
## 2 0.2750431 0.8998680      NA      NA      NA
##                              StatusML                         ChangedParams
## 1 Warning: Naive-ML difference > 10% minSporeFscHLog=5.0_widthAroundPeak=1
## 2                                  Ok                                  <NA>
```
