## Supplementary Figure S1 for "CAYSS: package for automatic Cytometry Analysis of Yeast Spore Segregation"

**Figure S1.** Manual analysis using the software 'Summit'. Top panel: manual gating of spore singlets. Left: SSC-Height-Log (Y-axis) *vs* FSC-Height-Log (X-axis) plot, showing the red circle drawn by the user to select spores from possible cellular debris (on the bottom-left side of the plot) or other types of cells (e.g. vegetative cells on the top-right side). Right: SSC-Height-Log (Y-axis) *vs* SSC-Area-Log (X-axis) plot used for manual gating of spore singlets, showing the polygon drawn by the user to select singlets from possible spore doublets or other cell aggregates. Bottom panel: manual gating of spore fluorescence intensities for the three 2D-scatter plots (left: mCherry-Height-Log *vs* CFP-Height-Log, middle: mCherry-Height-Log *vs* YFP-Height-Log, right: CFP-Height-Log *vs* YFP-Height-Log), showing the polygons drawn by the user to assign each spore singlet to a fluorescent classe.

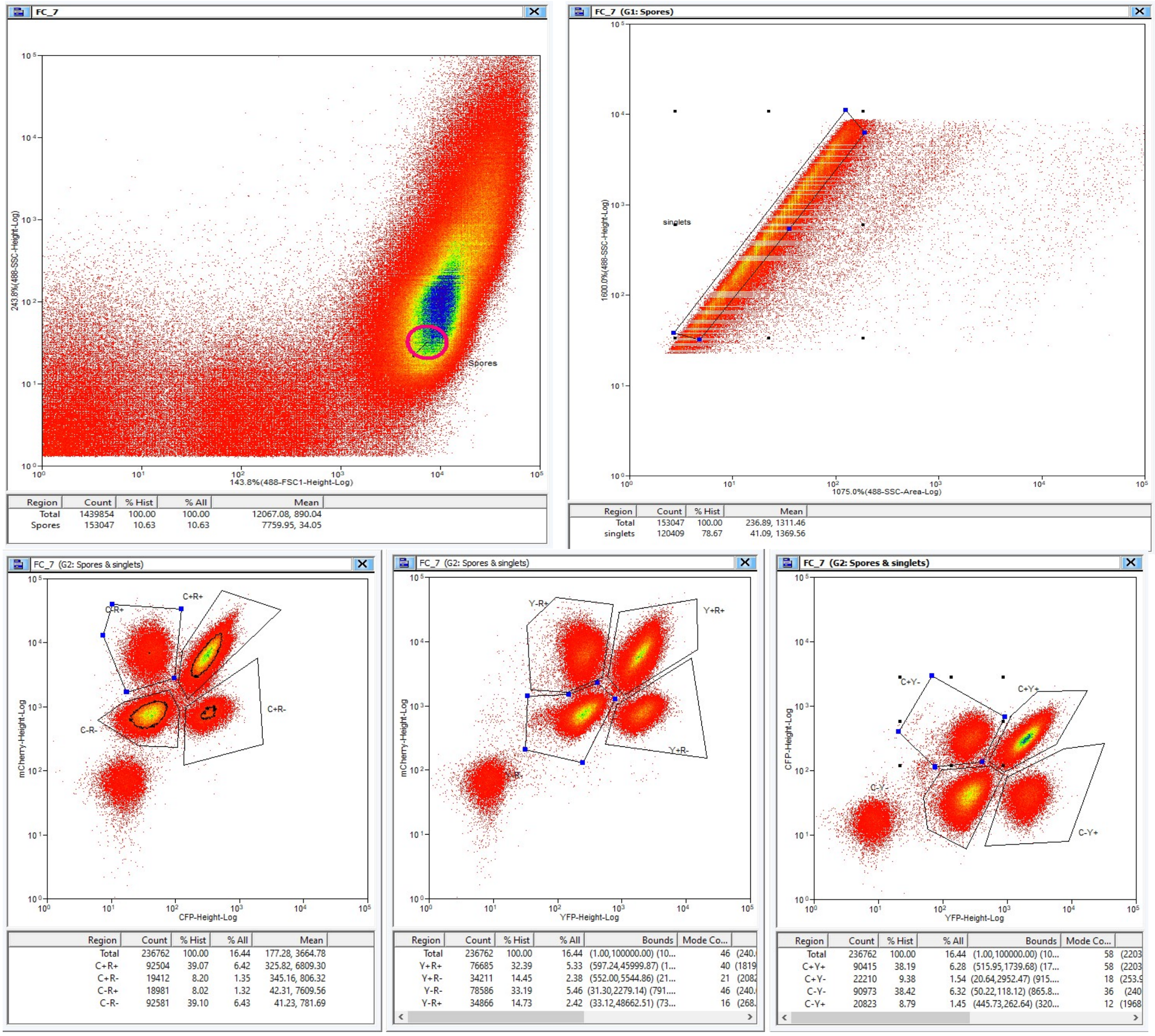
