## Supplementary Figure S2 for "CAYSS: package for automatic Cytometry Analysis of Yeast Spore Segregation"

**Figure S2.** Logical combinations of gates used in the manual analysis with the software "Summit" to assign events identified as spore singlets to each of the eight three-locus genotypic classes N (no marker present), C, R, Y, CR, CY, RY, and CRY corresponding to the fluorescent markers C (yECerulean), R (mCHerry), and Y (Venus).

Gate Logic Builder

| Name | Expression |
|---|---|
| G1 | Spores |
| G2 | Spores & singlets |
| RCY | C+R+ & C+Y+ & Spores & Y+R+ & singlets |
| N | C-R- & C-Y- & Spores & Y-R- & singlets |
| R | C-R+ & C-Y- & Spores & Y-R+ & singlets |
| CY | C+R- & C+Y+ & Spores & Y+R- & singlets |
| RC | C+R+ & C+Y- & Spores & Y-R+ & singlets |
| Y | C-R- & C-Y+ & Spores & Y+R- & singlets |
| RY | C-R+ & C-Y+ & Spores & Y+R+ & singlets |
| C | C+R- & C+Y- & Spores & Y-R- & singlets |
