## Supplementary Figure S3 for "CAYSS: package for automatic Cytometry Analysis of Yeast Spore Segregation"

**Figure S3.** Graphical summary output of CAYSS analysis of a typical sample with three fluorescent markers. The plots show spore (top left) and spore singlet selection (top right), then the distributions of fluorescence intensities for each marker, and then the recombination values computed naively (left column of bottom-right panel) and by ML inference with a model incorporating fluorescence extinction probabilities for each marker (right column of bottom-right panel)

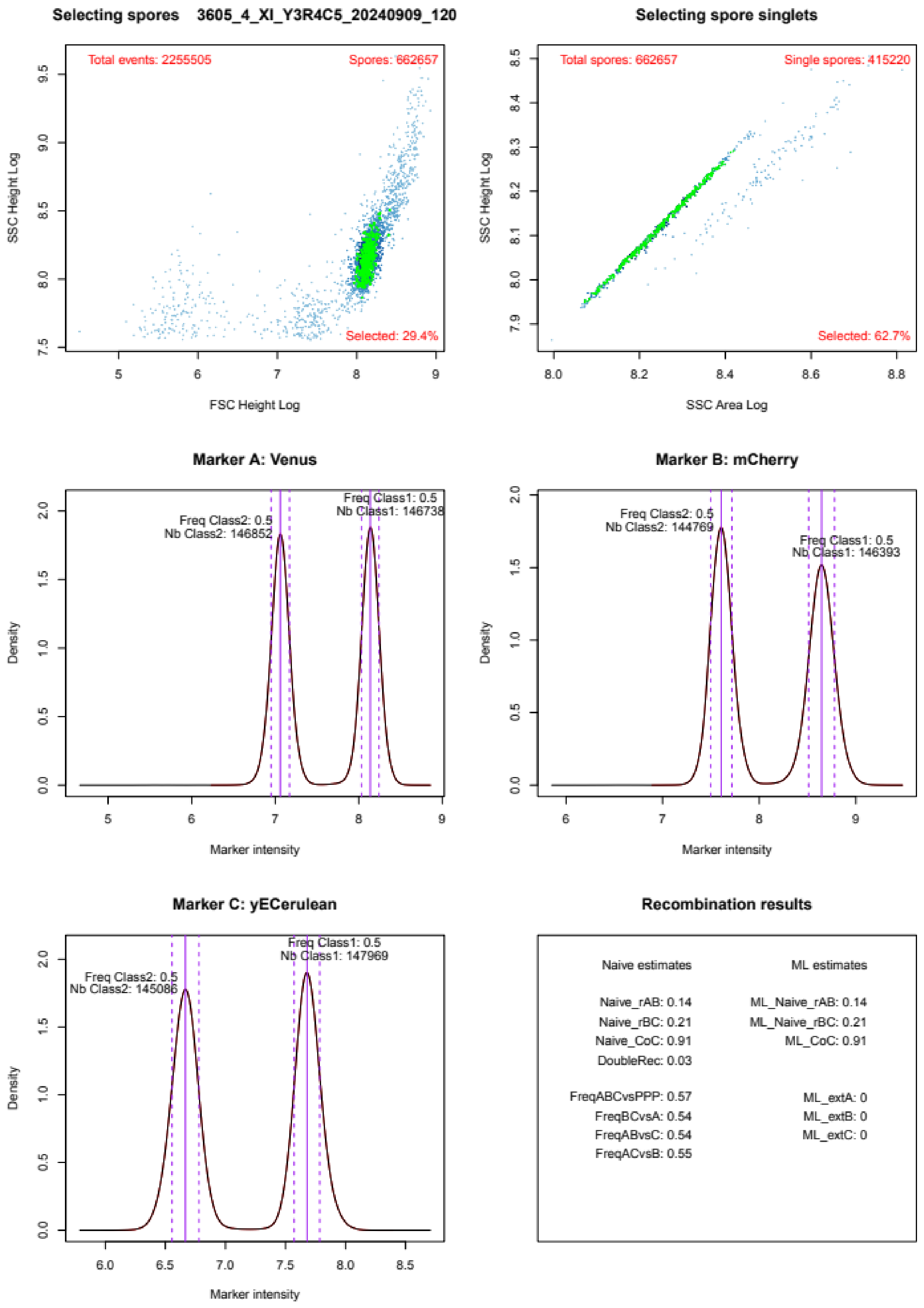
