## Supplementary Figure S4 for "CAYSS: package for automatic Cytometry Analysis of Yeast Spore Segregation"

**Figure S4.** Assessment of fit quality for the Maximum-Likelihood inference. Distributions of the inferred values after Maximum-Likelihood fit of recombination parameters (recombination rates rAB and rBC, and coefficient of coincidence CoC, top panel) across 1000 replicates with different initial values, using a model which incorporates fluorescence extinction for the three markers (extinctions probabilities Aext, Bext, and Cext, bottom panel). In this case, the best fit for Aext and Bext is zero, and all replicates gave the same value, which means that fluorescence extinction was not observed for markers A and C.

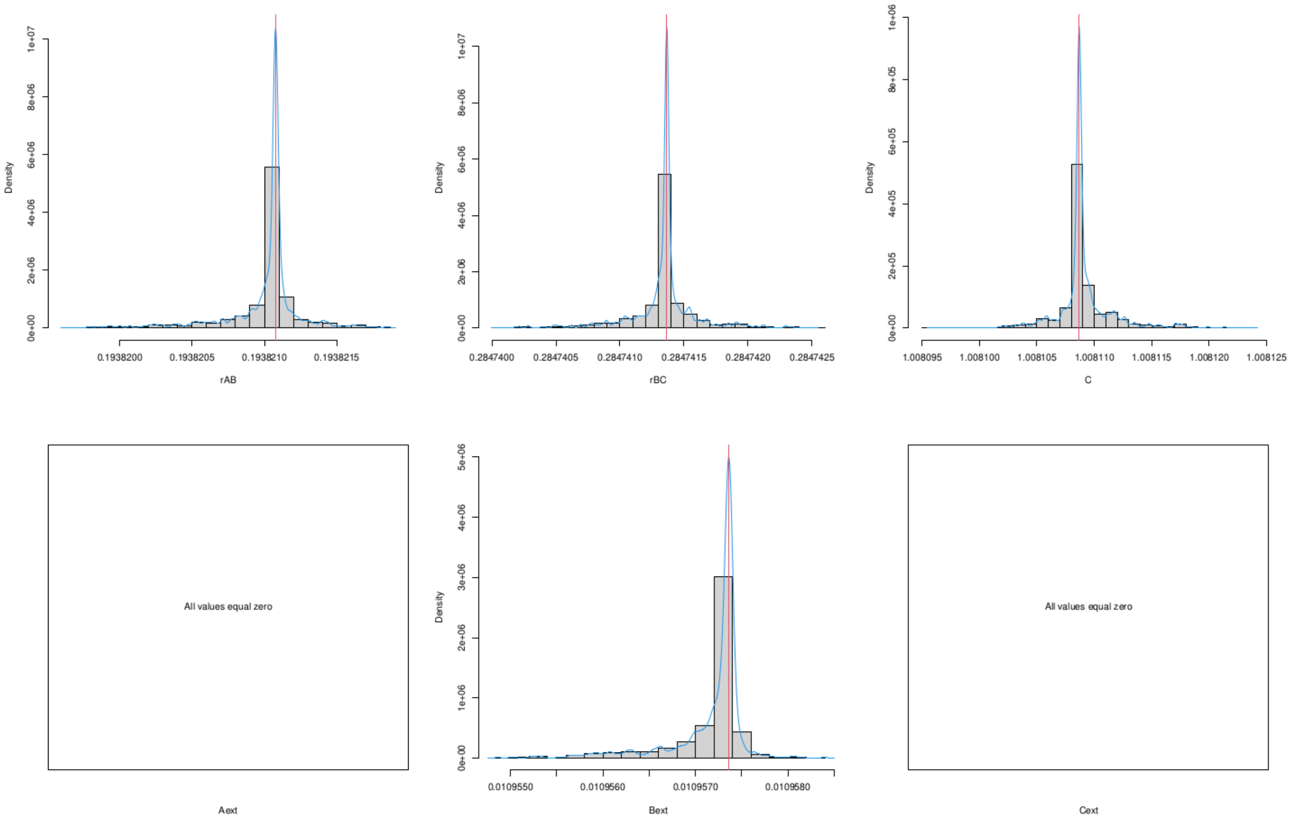
