## Supplementary Figure S5 for "CAYSS: package for automatic Cytometry Analysis of Yeast Spore Segregation"

**Figure S5.** Sample with only one peak in fluorescence intensity distribution. After automatic detection of peaks, when a single peak is detected, the sample is skipped but the density distribution of fluorescence intensity is plotted, so one may assess if different parameters should be tried to rescue a possible second peak.

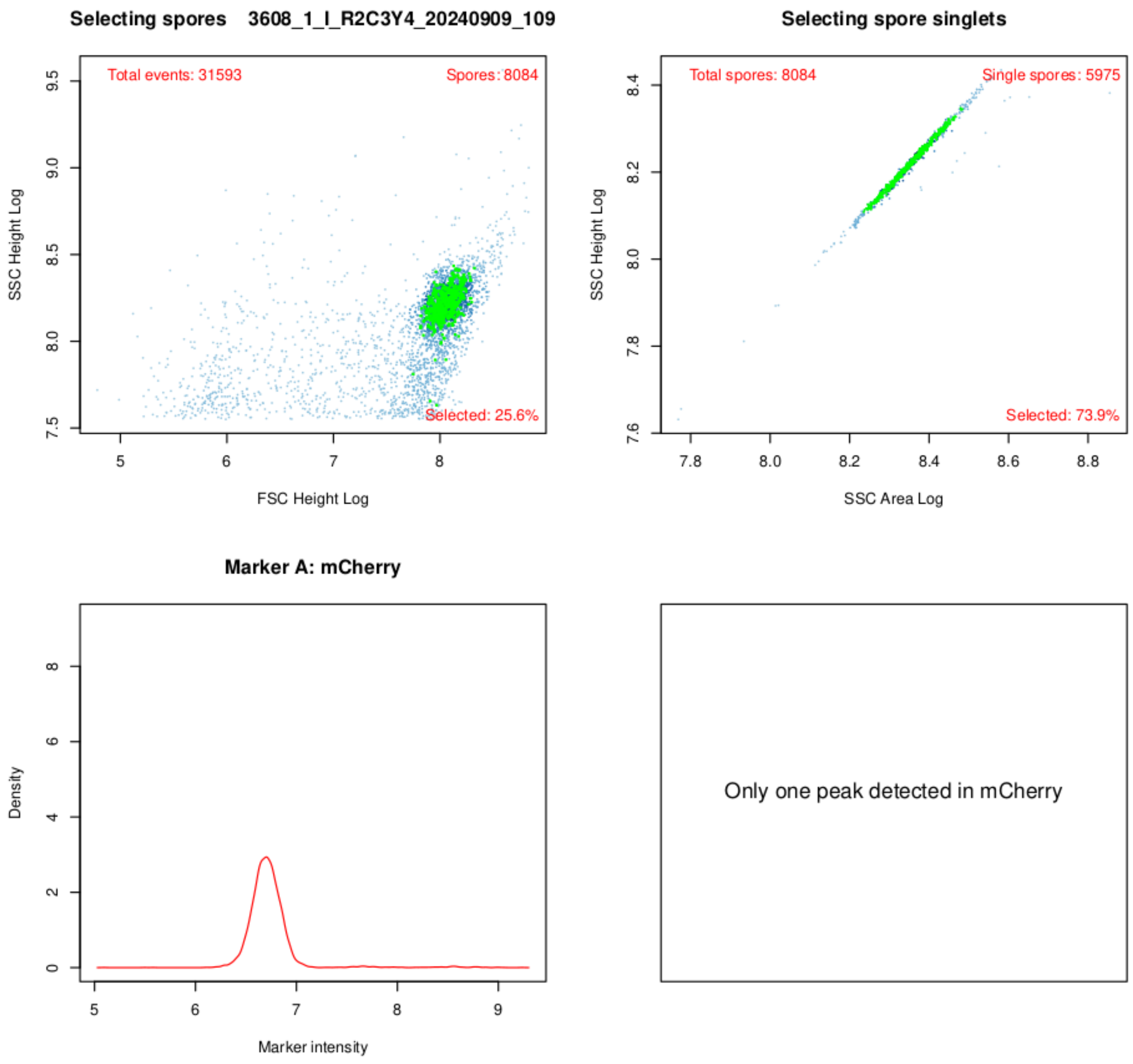
