## Supplementary Figure S6 for "CAYSS: package for automatic Cytometry Analysis of Yeast Spore Segregation"

**Figure S6.** CAYSS graphical output in the case of only two markers. In addition to the distributions drawn for each marker, an additional 2-D plot of CFP (Y-axis) *vs* YFP (X-axis) for spore singlets, shows four clouds of spores: non-fluorescent (parental, bottom-left), bi-fluorescent (parental, top-right), mCherry-only (recombinant, top-left), and Venus-only (recombinant, bottom-right).

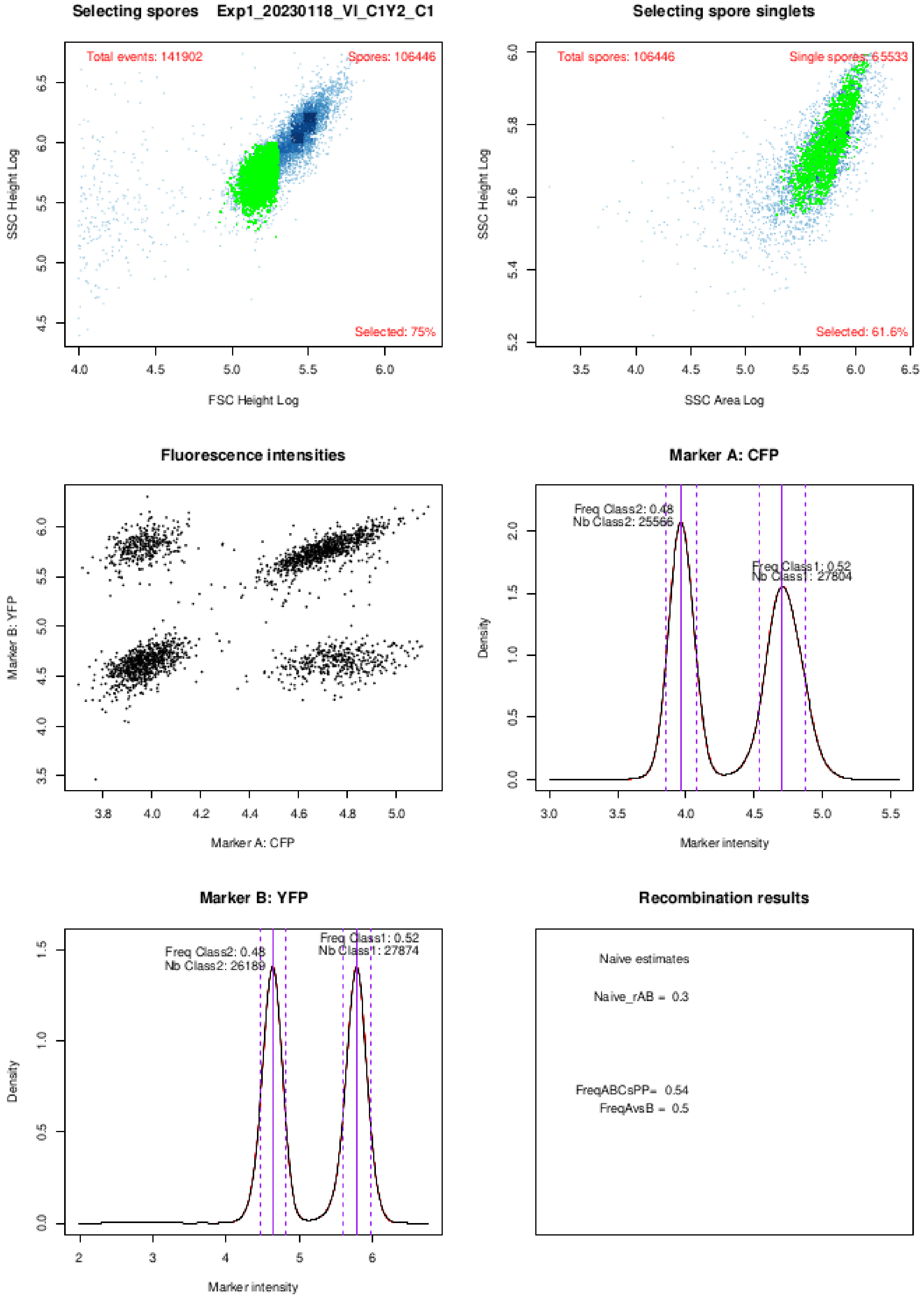
