## Supplementary material for "CAYSS: package for automatic Cytometry Analysis of Yeast Spore Segregation": CAYSS R package: CAYSS.R

```
#***********************************************************************
# Copyright (c) 2024 Matthieu Falque <>.
#
# This file is part of CAYSS
#
#  CAYSS is free software: you can redistribute it and/or modify
#  it under the terms of the GNU General Public License as published by
#  the Free Software Foundation, either version 3 of the License, or
#  (at your option) any later version.
#  CAYSS is distributed in the hope that it will be useful,
#  but WITHOUT ANY WARRANTY; without even the implied warranty of
#  MERCHANTABILITY or FITNESS FOR A PARTICULAR PURPOSE.  See the
#  GNU General Public License for more details.
#
#  You should have received a copy of the GNU General Public License
#  along with CAYSS If not, see <http://www.gnu.org/licenses/>.
#
#***********************************************************************

#' @description
#' The package CAYSS (Cytometry Analysis of Yeast Spores Segregation) allows
#' automatic random spore analysis from flow cytometry data to study the
#' segregation of two or three fluorescent markers and calculate rates of
#' recombination and crossover interference, based on the method described in
#' Raffoux et al. 2018 Yeast(6):431. \cr
#' CAYSS automatically selects events corresponding to spore singlets and
#' analyzes fluorescence intensities for each marker to assign each spore to 
#' one of the classes 'marker present' or 'marker absent'. It then 
#' computes recombination fractions in intervals between adjacent markers,
#' as well as (in the case of three markers) the coefficient of coincidence 
#' (CoC) which indicates the effect of crossover interference. \cr
#' Based on a mathematical model described in the same paper, which incorporates
#' partial fluorescence extinction of some of the markers, CAYSS also infers 
#' recombination, interference, and extinction parameters by 
#' Maximum-Likelihood. \cr
#' The spores must be produced by diploid cells hemizygous for the
#' genes encoding the fluorescent markers, and these genes must be located on 
#' the same chromosome, and genetically linked. Fluorescent strains to be used 
#' to analyze recombination are available upon request to 
#' \cr
"_PACKAGE"

#' @import stringr
#' @import KernSmooth
#' @import cluster
#' @import qpdf
#' @import flowCore
#' @importFrom grDevices colorRampPalette densCols dev.set dev.cur dev.off pdf
#' @importFrom graphics abline hist legend lines par points text
#' @importFrom stats density lm optim runif sd smooth.spline var
#' @importFrom utils install.packages read.table write.table
#' @importFrom methods is

# @name S2c
S2c <- function (string) {
  return(unlist(strsplit(string,"")))
}

#' @name ExampleFile
#' @title Returns path to example files
#' @description Gives path to CAYSS example input files
#' @param type Character string with name of the example file, or NULL. 
#' Example files may be one of the following:
#' - "PlateDesign" complete path to the plate design file that contains the list
#'    of all example samples with the ID corresponding to the FCS file 
#'    (e.g. plate coordinates), as described for the 'plateDesignFile' 
#'    parameter of function \code{\link{AnalyzeRecombination}}
#' - "FCS1" raw output file of the flow cytometer for example with two markers
#' - "FCS2" raw output file of the flow cytometer for example with three markers
#'    without extra peaks
#' - "FCS3" raw output file of the flow cytometer for example with three markers
#'    with extra peaks due to the presence in the sample of non-fluorescent 
#'    spores coming from non-fluorescent diploid mother cells. The third peak 
#'    only appears for markers mCherry and Venus
#' - NULL then the list of files contained in folder "extdata" of the package
#'    is returned as a vector
#' @return Character string or vector of character strings:
#' - if Character string: name with complete path of different types of
#'  example CAYSS input files
#' - if vector of Character strings: list of files contained in the folder 
#' "(/inst/)extdata"
#' @examples
#' ExampleFile("PlateDesign123")
#' @export
ExampleFile <- function(type=NULL) {
  
  userExampleDir <- normalizePath(file.path(Sys.getenv("HOME"),"CAYSS_example"), mustWork=FALSE)
  if( !dir.exists(userExampleDir) ) {
    success = dir.create(userExampleDir, recursive=TRUE, mode="0755")
    if( !success ){
      stop( paste("Could not create directory:", userExampleDir, sep='\n'), domain=NA )
    }
  }
  
  if (is.null(type)) {
    return(system.file("extdata", package="CAYSS") |> list.files())
  } else if (type=="exampleFolder") {
    return(userExampleDir)
  } else if (type=="PlateDesign1") {
    file.copy(from=system.file("extdata", "Example_Plate_Design_1.txt", package="CAYSS"),
              to=userExampleDir, overwrite=TRUE)
    file.copy(from=system.file("extdata", "Exp1_C1.fcs", package="CAYSS"),
              to=userExampleDir, overwrite=TRUE)
    return(file.path(userExampleDir, "Example_Plate_Design_1.txt"))
  } else if (type=="PlateDesign2") {
    file.copy(from=system.file("extdata", "Example_Plate_Design_2.txt", package="CAYSS"),
              to=userExampleDir, overwrite=TRUE)
    file.copy(from=system.file("extdata", "Exp2_E1.fcs", package="CAYSS"),
              to=userExampleDir, overwrite=TRUE)
    return(file.path(userExampleDir, "Example_Plate_Design_2.txt"))
    
  } else if (type=="PlateDesign3") {
    file.copy(from=system.file("extdata", "Example_Plate_Design_3.txt", package="CAYSS"),
              to=userExampleDir, overwrite=TRUE)
    file.copy(from=system.file("extdata", "Exp3_G7.fcs", package="CAYSS"),
              to=userExampleDir, overwrite=TRUE)
    return(file.path(userExampleDir, "Example_Plate_Design_3.txt"))
    
  } else if (type=="PlateDesign123") {
    file.copy(from=system.file("extdata", "Example_Plate_Design_1-2-3.txt", package="CAYSS"),
              to=userExampleDir, overwrite=TRUE)
    file.copy(from=system.file("extdata", "Exp1_C1.fcs", package="CAYSS"),
              to=userExampleDir, overwrite=TRUE)
    file.copy(from=system.file("extdata", "Exp2_E1.fcs", package="CAYSS"),
              to=userExampleDir, overwrite=TRUE)
    file.copy(from=system.file("extdata", "Exp3_G7.fcs", package="CAYSS"),
              to=userExampleDir, overwrite=TRUE)
    return(file.path(userExampleDir, "Example_Plate_Design_1-2-3.txt"))
    
  } else if (type=="PlateDesignParams") {
    file.copy(from=system.file("extdata", "Example_Plate_Design_1-2-3_PARAMS.txt", package="CAYSS"),
              to=userExampleDir, overwrite=TRUE)
    file.copy(from=system.file("extdata", "Exp2_E1.fcs", package="CAYSS"),
              to=userExampleDir, overwrite=TRUE)
    file.copy(from=system.file("extdata", "Exp3_G7.fcs", package="CAYSS"),
              to=userExampleDir, overwrite=TRUE)
    return(file.path(userExampleDir, "Example_Plate_Design_1-2-3_PARAMS.txt"))
    
  } else {
    stop(" in ExampleFile(): argument 'type' must be 'PlateDesignXXX' or 'exampleFolder'")
  }
}


# @name Par2freqs3loci
# @title Computes expected genotype frequencies from model parameters
# @description Computes expected frequencies of fluorescence classes for given
# values of rec, interference, and gene conversion parameters, as described
# in Raffoux et al. 2018 Yeast(6):431 
# @param par Named Vector of Numeric. Elements are:
# \itemize{
#   \item rAB recombination fraction between markers A and B
#   \item rBC recombination fraction between markers B and C
#   \item C Coefficient of Coincidence, defined as observed freq of double 
#   recombinants over expected freq of doubles recombinants in the absence of 
#   crossover interference
#   \item Aext freq of fluorescence extinction for marker A
#   \item Bext freq of fluorescence extinction for marker B
#   \item Cext freq of fluorescence extinction for marker C
# }
# @return segregation Vector of Numeric. Frequencies of the expected 
# frequencies of genotypes (in fact fluorescence phenotypes) for the classes:
# fA, fAB, fABC, fAC, fB, fBC, fC, and fN (non-fluorescent)
# @examples Par2freqs3loci(par=c(rAB=0.2, rBC=0.3, C=0.9, Aext=0, Bext=0, Cext=0.05))
Par2freqs3loci <- function(par) {
  rAB <- par[1]
  rBC <- par[2]
  C <-   par[3]
  Aext <- par[4]
  Bext <- par[5]
  Cext <- par[6]
  #   note that with interference, 
  # one has:	(1-rAB)*(1-rBC)	=>   (1-rAB-rBC+C*rAB*rBC)
  # 											rAB*(1-rBC)		=>   rAB*(1-C*rBC)
  # 											(1-rAB)*rBC		=>   (1-C*rAB)*rBC
  # 											rAB*rBC			=>   C*rAB*rBC
  fN <-	1/2 * (1-rAB-rBC+C*rAB*rBC) +
    1/2 * rAB*(1-C*rBC)*Aext +
    1/2 * (1-C*rAB)*rBC*Cext +
    1/2 * rAB*C*rBC*Bext +
    1/2 * (1-C*rAB)*rBC*Aext*Bext +
    1/2 * rAB*(1-C*rBC)*Bext*Cext +
    1/2 * rAB*C*rBC*Aext*Cext +
    1/2 * (1-rAB-rBC+C*rAB*rBC)*Aext*Bext*Cext

  fABC <-	1/2 *(1-rAB-rBC+C*rAB*rBC)*(1-Aext)*(1-Bext)*(1-Cext)

  fA <-	1/2 * rAB*(1-C*rBC)*(1-Aext) +
    1/2 * (1-C*rAB)*rBC*(1-Aext)*Bext +
    1/2 * rAB*C*rBC*(1-Aext)*Cext +
    1/2 * (1-rAB-rBC+C*rAB*rBC)*(1-Aext)*Bext*Cext

  fB <-	1/2 * rAB*C*rBC*(1-Bext) +
    1/2 * (1-C*rAB)*rBC*(1-Bext)*Aext +
    1/2 * rAB*(1-C*rBC)*(1-Bext)*Cext +
    1/2 * (1-rAB-rBC+C*rAB*rBC)*(1-Bext)*Aext*Cext

  fC <-	1/2 * (1-C*rAB)*rBC*(1-Cext) +
    1/2 * rAB*(1-C*rBC)*(1-Cext)*Bext +
    1/2 * rAB*C*rBC*(1-Cext)*Aext +
    1/2 * (1-rAB-rBC+C*rAB*rBC)*(1-Cext)*Aext*Bext

  fAB <-	1/2 * (1-C*rAB)*rBC*(1-Aext)*(1-Bext) +
    1/2 * (1-rAB-rBC+C*rAB*rBC)*(1-Aext)*(1-Bext)*Cext

  fBC <-	1/2 * rAB*(1-C*rBC)*(1-Bext)*(1-Cext) +
    1/2 * (1-rAB-rBC+C*rAB*rBC)*(1-Bext)*(1-Cext)*Aext

  fAC <-	1/2 * rAB*C*rBC*(1-Aext)*(1-Cext) +
    1/2 * (1-rAB-rBC+C*rAB*rBC)*Bext*(1-Aext)*(1-Cext)

  freqs <- c(fA, fAB, fABC, fAC, fB, fBC, fC, fN)
  names(freqs) <- c("fA", "fAB", "fABC", "fAC", "fB", "fBC", "fC", "fN")

  return(freqs)
}


# @name LogLike3loci
# @title Computes log-likelihood of observations given model parameters
# @description Classical computation of likelihood
# @inheritParams Par2freqs3loci
# @param obs Vector (size 8) of Numeric. Observations of numbers of spores
# for the phenotypic classes fA, fAB, fABC, fAC, fB, fBC, fC, and fN 
# (non-fluorescent)
# @return Numeric. Log-likelihood
# @examples LogLike3loci(par=c(rAB=0.2, rBC=0.3, C=0.9, Aext=0, Bext=0, 
# Cext=0.05), obs=c(fA=126, fAB=519, fABC=1002, fAC=69, fB=54, fBC=114, 
# fC=527, fN=783))
# @export
LogLike3loci <- function(par, obs) {
  if (length(par)==3) par=c(par,Aext=0,Bext=0,Cext=0) #ADD Xext=0 in case of FIT_CONV=FALSE
  freqs <- Par2freqs3loci(par)
  freqs[freqs < 1e-200] <- 1e-200
  ll <- sum( obs * log(freqs) )
  return(ll)
}


# @name MlFitParams3loci
# @title Finds parameters maximizing the log-likelihood of observations
# @description Performs repeated fits from different initial values, and
# plots the distributions of corresponding parameters. Takes the mode of this 
# distribution
MlFitParams3loci <- function(obs, id, NB_REP_FIT=1000, FIT_CONV=TRUE) {

  if (FIT_CONV ) {
    lower <- c(rAB = 1e-6, rBC = 1e-6, C = 0.3, Aext = 0,   Bext = 0,   Cext = 0)
    upper <- c(rAB = 0.5,  rBC = 0.5,  C = 3,   Aext = 0.5, Bext = 0.5, Cext = 0.5)
  } else {
    upper <- c(rAB = 0.5,  rBC = 0.5,  C = 3)
    lower <- c(rAB = 1e-6, rBC = 1e-6, C = 0.3)
  }

  resReps <- NULL
  control <- list(fnscale = -1, factr = 1e3, maxit=100)
  methods <- c("Nelder-Mead", "BFGS", "CG", "L-BFGS-B", "SANN")
  i1 <- i2 <- i3 <- i4 <- i5 <- i6 <- 1
  iRep <- 1
  for (iRep in 1:NB_REP_FIT) {
    thisInitPar <- rep(NA, length(upper))
    for (iParam in 1:length(upper)) {
      thisInitPar[iParam] <- runif(1, lower[iParam], upper[iParam])
    }
    res <- optim(par=thisInitPar, lower=lower, upper=upper, fn=LogLike3loci,
                 method=methods[4], control=control, obs=obs)
    res$par
    resReps <- rbind(resReps, res$par)
  }
  #par(mfrow=c(1,length(upper)))
  par(mfrow=c(2,(length(upper)/2)))
  parMode <- rep(NA,length(upper))
  for (i in 1:length(upper))	{
    dens <- density(resReps[,i])
    if ( ! (all(is.numeric(resReps[,i]), na.rm=FALSE) 
            && any(resReps[,i]==0, na.rm=FALSE))) {
      hist(resReps[,i], proba=T, nclass=NB_REP_FIT/50, main=id, 
           xlab=names(lower)[i], ylim=c(0,max(dens$y)))
      lines(dens, col=4)
      parMode[i] <- dens$x[which.max(dens$y)]
      abline(v=parMode[i], col=2)
    } else {
      plot(0,0, type='n', xaxt='n', yaxt='n', xlab=names(lower)[i], ylab="")
      text(0,0,"All values equal zero")
    }
  }
  parMean <- apply(resReps, MARGIN=2, FUN=mean)
  parCv <- apply(resReps, MARGIN=2, FUN=function(x) return(sd(x)/mean(x)))
  theor <- Par2freqs3loci(parMode) * sum(obs)
  ssd <- sum((theor-obs)^2)
  parCv[is.nan(parCv)] <- 0
  return(list(parMean=parMean, parCv=parCv, parMode=parMode, ssd=ssd))
}


# @name MaxLikelihoodFit3loci
# @title Connects AnalyzeRecombination() with MaxLikelihoodFit3loci()
MaxLikelihoodFit3loci <- function(Data, mlPdfFile, logFile) {
  pdf(file=mlPdfFile, 15, 10)
  # obs is a vector with numbers of spores observed for each of the 8 classes of fluorescence
  App <- Data$App	# A++
  ABp <- Data$ABp	# AB+
  ABC <- Data$ABC	# ABC
  ApC <- Data$ApC  # A+C
  pBp <- Data$pBp	# +B+
  pBC <- Data$pBC	# +BC
  ppC <- Data$ppC	# ++C
  ppp <- Data$ppp	# +++
  obs <- c( App=App, ABp=ABp, ABC=ABC, ApC=ApC, pBp=pBp, pBC=pBC,ppC=ppC,ppp=ppp)
  Tot <- sum(obs)
  #Distorsion calculation
  FreqABCvsNNN <- ABC / (ppp+ABC)
  FreqBCvsA    <- pBC  / (App+pBC)
  FreqABvsC    <- ABp  / (ppC+ABp)
  FreqACvsB    <- ApC  / (pBp+ApC)
  #Rec rate and CoC without ML
  Naive_rAB  <- (App + pBC + ApC + pBp) / Tot
  Naive_rBC  <- (ABp + ppC + ApC + pBp) / Tot
  Doubl <- (ApC + pBp) / Tot
  CoC   <- Doubl / (Naive_rAB*Naive_rBC)
  id <- Data$sampleName
  resFit <- try(MlFitParams3loci(obs=obs, id=id , NB_REP_FIT=1000, 
                                 FIT_CONV=TRUE), silent=TRUE)
  if (all(is.numeric(as.matrix(resFit$parMode[1:6])))) {
    Data$TotSpores=Tot
    Data$ML_rAB <- resFit$parMode[1]
    Data$ML_rBC <- resFit$parMode[2]
    Data$ML_CoC <- resFit$parMode[3]
    Data$ML_Aext <- resFit$parMode[4]
    Data$ML_Bext <- resFit$parMode[5]
    Data$ML_Cext <- resFit$parMode[6]
    Data$StatusML <- "Ok"
  } else {
    Data$TotSpores <- "Error"
    Data$ML_rAB <- "Error"
    Data$ML_rBC <- "Error"
    Data$ML_CoC <- "Error"
    Data$ML_Aext <- "Error"
    Data$ML_Bext <- "Error"
    Data$ML_Cext <- "Error"
    Data$StatusML <- "ML failed to converge"
    msg <- "\nFit failed to converge in MaxLikelihood3loci\n"
    cat(msg) ; write(msg, file=logFile, append=TRUE)
  }
  dev.off()
  return(Data)
}


#======================  Function SelectionOnDensity
# @name SelectionOnDensity
# Select spores on SSC vs FSC GRAPH based on density of events
# Select Spores on SSC-Aire log  vs de SSC log GRAPH based on density of events
#PARAMETERS
#X = Values on X axe
#Y = Values on Y axe
#Number of bin for X and Y
#binNumberX = 100
#binNumberY = 100
#==> Do you want to see a grph of data, density and selected group (orange circle)
#graphose=T
# SelectCloudGeometry        == > "Circle", "Axis"
# nbDataForCalc ==> Number of data used for calculation. nbDataForCalc=30 000 ==> use 30 000 events to perform. If nbDataForCalc= "ALL" == > Use All data
# slopeFindPeak = 0.2     # Ratio value to stop Ratio when CV decrease rapidly (if 0.2 then scan will stop when CV is 0.2 X upper than before
# EXEMPLE : SelectionOnDensity(X=X, Y=Y, binNumberX=100, binNumberY=100, graphose=T, SelectCloudGeometry = "Circle", nbDataForCalc = 30000, slopeFindPeak = 0.2 )
SelectionOnDensity <- function(X, Y,  binNumberX, binNumberY, graphose,
                               SelectCloudGeometry, nbDataForCalc,
                               slopeFindPeak)  {
  Data <- data.frame (X=X, Y=Y)
  # Kepp only a sample of data to perform
  if(nbDataForCalc == "ALL" ) { SampleData <- Data
  } else {
    SampleData <- Data[sample(nrow(Data), nbDataForCalc), ]
    while (sum(
      (is.numeric(SampleData$X) & is.finite(SampleData$X)) &
      (is.numeric(SampleData$Y) & is.finite(SampleData$Y))
    ) < nbDataForCalc) {
      print("New try for sampling data in SelectionOnDensity()")
      SampleData <- Data[sample(nrow(Data), nbDataForCalc), ]
    }
  }
  MinX <- min (SampleData$X)
  MaxX <- max (SampleData$X)
  MinY <- min (SampleData$Y)
  MaxY <- max (SampleData$Y)
  RangeX <- MaxX - MinX
  RangeY <- MaxY - MinY
  SizeX <- RangeX / binNumberX  #Size of bin for X and Y axes
  SizeY <- RangeY / binNumberY
  BinX <- rep(1:binNumberX, each = binNumberY)  #Calculate Coordinates of bins (min and max for X and Y)
  BinY <- rep(1:binNumberY, binNumberX)
  BinCoord <- data.frame (binNumberX = BinX , binNumberY = BinY, Xmin=NA, Xmax=NA, Ymin=NA, Ymax=NA, ObsNumber=NA)
  for (i in (1: length(BinCoord$binNumberX)))  {
    BinCoord$Xmin[i] <- MinX + (BinCoord$binNumberX[i]-1) * SizeX
    BinCoord$Xmax[i] <- MinX + (BinCoord$binNumberX[i]) * SizeX
    BinCoord$Ymin[i] <- MinY + (BinCoord$binNumberY[i]-1) * SizeY
    BinCoord$Ymax[i] <- MinY + (BinCoord$binNumberY[i]) * SizeY
  }
  for (i in (1: length(BinCoord$binNumberX)))  { 
    #Calculate the number of observations in each bin
    ObservedInBinX  <-  SampleData[SampleData$X >= BinCoord$Xmin[i] &  SampleData$X < BinCoord$Xmax[i] ,]
    ObservedInBinXY <-  ObservedInBinX[ObservedInBinX$Y >= BinCoord$Ymin[i] &  ObservedInBinX$Y < BinCoord$Ymax[i] ,]
    ObsNumber <- length(ObservedInBinXY$X)
    BinCoord$ObsNumber[i] <- ObsNumber
  }
  # Scan Ratio to find the best CV of distance to the highest density bin in case of circle cloud  (Select "spores")
  #Get group of max density :
  #Max density
  MaxObsNumber <- max(BinCoord$ObsNumber)
  #Get X position of max density bin
  MaxDensityBinX <- c(BinCoord[which(BinCoord$ObsNumber==max(BinCoord$ObsNumber)), "Xmin"],
                      BinCoord[which(BinCoord$ObsNumber==max(BinCoord$ObsNumber)), "Xmax"] )
  MaxDensityBinY <- c(BinCoord[which(BinCoord$ObsNumber==max(BinCoord$ObsNumber)), "Ymin"],
                      BinCoord[which(BinCoord$ObsNumber==max(BinCoord$ObsNumber)), "Ymax"] )
  if(SelectCloudGeometry == "Circle") {
    Scan <- NULL
    # par(mfrow = c(4,5))
    for (Ratio in seq(0.1, 0.95, 0.05)) {
      #Select neighbour
      HighObsNumber <- BinCoord[BinCoord$ObsNumber > MaxObsNumber * Ratio, ]
      #And select on the distance to MaxDensityBin
      MaxDensityBinX_Center <- MaxDensityBinX[1] + SizeX/2
      MaxDensityBinY_Center <- MaxDensityBinY[1] + SizeY/2
      # Calculate distance to Max Bin
      HighObsNumber$DistToMaxObsNumber <- sqrt(
        ((HighObsNumber$Xmin + SizeX/2)-MaxDensityBinX_Center)^2
        +((HighObsNumber$Ymin+SizeY/2)-MaxDensityBinY_Center)^2)
      #Sort by distance to Max Bin
      HighObsNumber <- HighObsNumber[order(HighObsNumber$DistToMaxObsNumber), ] 
      #Delete high distance to max bin
      #NbBinToKeep <- length (HighObsNumber$DistToMaxObsNumber) * DistanceRatio
      #HighObsNumber <- HighObsNumber[1:NbBinToKeep, ]
      #plot(SampleData$X, SampleData$Y, col=densCols(SampleData$X, SampleData$Y, nbin=30), cex=0.1, ,main = Ratio, xlab="X", ylab="Y" )
      #points(HighObsNumber$Xmax - SizeX/2 ,  HighObsNumber$Ymax - SizeY/2, col="orange" , pch=0, lwd=2)
      Df <- data.frame(
        Ratio=Ratio, 
        length=length (HighObsNumber$DistToMaxObsNumber), 
        mean=mean(HighObsNumber$DistToMaxObsNumber), 
        var=var(HighObsNumber$DistToMaxObsNumber), 
        CV=sqrt(var(HighObsNumber$DistToMaxObsNumber))/mean(HighObsNumber$DistToMaxObsNumber))
      Scan <- rbind.data.frame(Scan, Df)
    }
    #Get the minimal value of CV and get Ratio value
    # Plot CV vs Ratio ==  >   plot (Scan$Ratio, Scan$CV)
    # Plot CV vs Mean ==  >    plot (Scan$Ratio, Scan$mean)
    # Ignore CV calculated with less than 10 bins
    Scan <- Scan[Scan$length > 10 , ]
    # Order Scan
    Scan <- Scan[order(Scan$Ratio, decreasing=T), ]
    # Get the slope cutOff
    FindCvPeak <- NULL
    FindCvPeakResults <- NULL
    for (Ratio in Scan$Ratio) {
      ThisScanData <- Scan[Scan$Ratio >= Ratio & Scan$Ratio <= Ratio+0.05 , ]
      mean <- mean(ThisScanData$mean )
      FindCvPeak <- c(FindCvPeak, mean)
      # Difference with previous mean Ratio
      Diff <- mean - mean(FindCvPeak)
      if(Diff > mean(FindCvPeak) * slopeFindPeak ) break
      Df <- data.frame(Ratio = Ratio, MeanCV=mean, DiffCV=Diff)
      FindCvPeakResults <- rbind.data.frame(FindCvPeakResults, Df)
    }
    # Get min value of CV
    Ratio <- min(FindCvPeakResults$Ratio )
    #Select neighbour using this Ratio
    HighObsNumber <- BinCoord[BinCoord$ObsNumber > MaxObsNumber * Ratio, ]
  }
  # Scan Ratio to find the best CV of distance to the axis in case of Axis cloud  (Select "single")
  if(SelectCloudGeometry == "Axis") {
    # Get axis parameters of the highets density events
    # Select highest density bins : Get only the 10  hihest density
 #   BinCoordNotNA <- BinCoord[!is.na(BinCoord$Xmax) & !is.na(BinCoord$Ymax),]
    BinCoord <- BinCoord[order(BinCoord$ObsNumber, decreasing = TRUE), ]
    HighObsCloud <- BinCoord[1:10, ]
    # Get axis parameters (y=ax + b)
    Dframe <- data.frame( x=HighObsCloud$Xmax - SizeX/2, y=HighObsCloud$Ymax - SizeY/2)
    Reglin <- lm(y ~ x, Dframe)
    Intercept <- summary(Reglin)$coefficients[1,1]
    Slope <- summary(Reglin)$coefficients[2,1]
    #   plot(SampleData$X, SampleData$Y, cex=0.1, ,main = "X/Y GRAPH", xlab="X", ylab="Y" )
    # 	points(HighObsCloud$Xmax - SizeX/2 ,  HighObsCloud$Ymax - SizeY/2, col="orange" , pch=0, lwd=2)
    #   abline(Reglin, col="orange")
    # GET STD ERROR
    IntercepStdError <- summary(Reglin)$coefficients[1,2]
    SlopeStdError    <- summary(Reglin)$coefficients[2,2]
    MinIntercept <- Intercept - IntercepStdError
    MaxIntercept <- Intercept + IntercepStdError
    #abline(MinIntercept, Slope, col="chartreuse4")
    #abline(MaxIntercept, Slope, col="chartreuse4")
    # Measure the distance to the axis of events for differents Density Ratio
    #par(mfrow = c(4,5))
    Scan <- NULL
    for (Ratio in seq( 0 , 0.95 , 0.05)) {
      #Select neighbour
      HighObsNumber <- BinCoord[BinCoord$ObsNumber > MaxObsNumber * Ratio, ]
      # Calculate distance to the axis
      Num <- abs(Slope*(HighObsNumber$Xmax-SizeX/2)-(HighObsNumber$Ymax-SizeX/2)+Intercept  )
      Denom <-  sqrt (Slope^2 + 1)
      HighObsNumber$DistToAxis <- round (Num/ Denom, 6)
      Df <- data.frame(
        Ratio=Ratio, 
        length=length (HighObsNumber$DistToAxis), 
        mean=mean (HighObsNumber$DistToAxis), 
        var=var(HighObsNumber$DistToAxis), 
        CV=sqrt(var (HighObsNumber$DistToAxis))/mean(HighObsNumber$DistToAxis))
      Scan <- rbind.data.frame(Scan, Df)
      #plot(SampleData$X, SampleData$Y, cex=0.1, ,main = Ratio, xlab="X", ylab="Y" )
      #points(HighObsNumber$Xmax - SizeX/2 ,  HighObsNumber$Ymax - SizeY/2, col="orange" , pch=0, lwd=2)
      #abline(Reglin)
    }
    # Ignore CV calculated with less than 10 bins
    Scan <- Scan[Scan$length >10 , ]
    # Order Scan
    Scan <- Scan[order(Scan$Ratio, decreasing=T), ]
    # Get the slope cutOff
    FindCvPeak <- NULL
    FindCvPeakResults <- NULL
    for (Ratio in Scan$Ratio) {
      ThisScanData <- Scan[Scan$Ratio >= Ratio & Scan$Ratio <= Ratio+0.05 , ]
      mean <- mean(ThisScanData$mean )
      FindCvPeak <- c(FindCvPeak, mean)
      # Difference with previous mean Ratio
      Diff <- mean-mean(FindCvPeak)
      if(Diff > mean(FindCvPeak) * slopeFindPeak) break
      Df <- data.frame(Ratio = Ratio, MeanCV=mean, DiffCV=Diff)
      FindCvPeakResults <- rbind.data.frame(FindCvPeakResults, Df)
    }
    # Get mini value of CV
    Ratio <- min(FindCvPeakResults$Ratio )
    #Select neighbour using this Ratio
    HighObsNumber <- BinCoord[BinCoord$ObsNumber > MaxObsNumber * Ratio, ]
  }
  if 	(graphose==T) {
    plot(SampleData$X, SampleData$Y, cex=0.1, ,main = "X/Y GRAPH", xlab="X", ylab="Y" )
    BinCoord <- BinCoord[order (BinCoord$ObsNumber), ]
    ColorsFunction <- colorRampPalette(c("green", "red"))
    # points(BinCoord$Xmax - SizeX/2 ,  BinCoord$Ymax - SizeY/2, col=ColorsFunction(length(BinCoord$ObsNumber)) , pch=19)
    points(HighObsNumber$Xmax - SizeX/2 ,  HighObsNumber$Ymax - SizeY/2, col="orange" , pch=0, lwd=2)
  }
  # Get events of Data in selected bins
  # Get events of Data in selected bins (not to work with data out of range)
  # Keep only Data which are in range of X and Y bins
  MinX <- min (HighObsNumber$Xmin)
  MaxX <- max (HighObsNumber$Xmax)
  MinY <- min (HighObsNumber$Ymin)
  MaxY <- max (HighObsNumber$Ymax)
  DataInRange <- Data[ Data$X >= MinX & Data$X <= MaxX   &   Data$Y >= MinY & Data$Y <= MaxY, ]
  #Select Data in range of each bin
  # Get indices lignes of data to select
  LignesOK=NULL
  for (i in 1: length(HighObsNumber$Xmax)) {
    #For the i bin
    Xmin <-  HighObsNumber[i , "Xmin"]
    Xmax <-  HighObsNumber[i , "Xmax"]
    Ymin <-  HighObsNumber[i , "Ymin"]
    Ymax <-  HighObsNumber[i , "Ymax"]
    Vec <- which (DataInRange$X >= Xmin & DataInRange$X <= Xmax   &   DataInRange$Y >= Ymin & DataInRange$Y <= Ymax)
    LignesOK <- c (LignesOK, Vec)
  }
  #Delete double data
  LignesOK <- unique (LignesOK)
  # Keep only Data in bins
  SelData <- DataInRange[LignesOK, ]
  return( list (SelData=SelData, HighObsNumber=HighObsNumber, Scan=Scan))
}

# Use the FindPeaksAndValleys function on these data, with these parameters
# intensityValues = a vector of intensity values (LOG)
# NbSeeed = number of random seed on the density function
# graphose = Do you want to see the graph of the density function ? If yes then value = TRUE, if no then value=FALSE
# LineEachOptim = Do you want to see each peaks and valleys on the graph ?  If yes then value = TRUE, if no then value=FALSE (peaks are grey lines and valleys green lines)
# Debug = Do you want to see the seeds course ?  If yes then value = TRUE, if no then value=FALSE (peaks are red dots and valleys green dots)
# R code is for example the following :
# intensityValues = DataSample$FSC1HeightLog
# PeaksValleys = FindPeaksAndValleys(intensityValues=intensityValues , nbSeeds=100, graphose=FALSE, LineEachOptim=FALSE, Debug=FALSE)
#
# @name FindPeaksAndValleys
# @title Find peaks and valleys of a distribution
# @description Used to determine the fluorescence levels of cytometry events
# @param intensityValues Vector of Numerics. Fluorescence intensity values (log)
# @param nbSeeeds Numeric. Number of random seeds for the density function
# @param graphose Logical. TRUE to see the graph of the density function
# @param LineEachOptim Logical. TRUE to see each peaks and valleys on the graph
# (peaks are grey lines and valleys green lines)
# @param Debug Logical. TRUE to see the seeds course (peaks are red dots and
# valleys green dots)
# @return a list with coordinates of peaks and valleys
FindPeaksAndValleys <- function(intensityValues, nbSeeds, graphose
                                , LineEachOptim, Debug )  {
  FSCDensity <- data.frame(FSCIntensity = density(
    intensityValues)$x, Density = density(intensityValues)$y )
  FSCDensitySmooth <- data.frame(
    FSCIntensitySmooth=smooth.spline(FSCDensity, df=50)$x, 
    FSCDensitySmooth=smooth.spline(FSCDensity, df=50)$y)
  DataX <- FSCDensitySmooth$FSCIntensitySmooth
  DataY <- FSCDensitySmooth$FSCDensitySmooth
  ResultsPeaks=NULL
  if (Debug==TRUE) plot (DataX, DataY, cex=0.2)
  for (i in 1:nbSeeds) {
    PosX_Random=NULL
    PosX_Random[1] = sample(5:length(DataX)-1,1)
    ValueY_Random=1
    ValueY_After=2
    ValueY_Before=2
    Iter=1
    OptimProcess="Run"
    while (OptimProcess!="End" & Iter<300) {
      if((ValueY_Random < ValueY_After) |(ValueY_Random < ValueY_Before ) ){
        ValueX_Random <- DataX[PosX_Random[Iter]]
        ValueY_Random <- DataY[PosX_Random[Iter]]
        ValueY_Before <- DataY[PosX_Random[Iter] - 1]
        ValueY_After <-  DataY[PosX_Random[Iter] + 1]
        if (identical(ValueY_Before, numeric(0))) ValueY_Before <- min (DataY)
        if (identical(ValueY_After , numeric(0))) ValueY_After  <- min (DataY)
        if (is.na(ValueY_Before)) {  ValueY_Before <- min (DataY) }
        if (is.na(ValueY_After))  {  ValueY_After  <- min (DataY) }
        if (Debug==TRUE) points (ValueX_Random, ValueY_Random, col="red")
        Df <- data.frame( PosX_Random=PosX_Random[Iter], 
                          ValueX_Random=ValueX_Random, 
                          ValueY_Before=ValueY_Before, 
                          ValueY_Random=ValueY_Random, 
                          ValueY_After=ValueY_After)
        if (ValueY_Random < ValueY_After) 
          PosX_Random[Iter+1] <- PosX_Random[Iter] +1 
        if (ValueY_Random < ValueY_Before) 
          PosX_Random[Iter+1] <- PosX_Random[Iter] - 1 
        if (ValueY_Random >= ValueY_Before & ValueY_Random >= ValueY_After ) 
          OptimProcess="End"
        Iter=Iter+1
      }
    }
    ValueX_Peak <- Df$ValueX_Random
    ValueY_Peak <- Df$ValueY_Random
    Df <- data.frame(ValueX_Peak=ValueX_Peak, ValueY_Peak=ValueY_Peak)
    ResultsPeaks <- rbind.data.frame (ResultsPeaks, Df)
  }
  ResultsPeaks <- unique (ResultsPeaks)
  ResultsPeaks <- ResultsPeaks[order(ResultsPeaks$ValueY_Peak, 
                                     decreasing=TRUE), ]
  #FIND VALLEYS
  ResultsValleys=NULL
  for (i in 1:nbSeeds) {
    PosX_Random=NULL
    PosX_Random[1] <-   sample(5:length(DataX)-1,1)
    ValueY_Random=3
    ValueY_After=2
    ValueY_Before=2
    Iter=1
    OptimProcess="Run"
    while (OptimProcess!="End" & Iter<300) {
      if((ValueY_Random > ValueY_After) |(ValueY_Random > ValueY_Before)  ){
        ValueX_Random <- DataX[PosX_Random[Iter]]
        ValueY_Random <- DataY[PosX_Random[Iter]]
        ValueY_Before <- DataY[PosX_Random[Iter] - 1]
        ValueY_After <-  DataY[PosX_Random[Iter] + 1]
        if (identical(ValueY_Before, numeric(0))) ValueY_Before <- max (DataY)
        if (identical(ValueY_After , numeric(0))) ValueY_After <- max (DataY)
        if (is.na(ValueY_Before)) ValueY_Before <- max (DataY)
        if (is.na(ValueY_After)) ValueY_After  <- max (DataY)
        Df <- data.frame(PosX_Random=PosX_Random[Iter], 
                         ValueX_Random=ValueX_Random, 
                         ValueY_Before=ValueY_Before, 
                         ValueY_Random=ValueY_Random, 
                         ValueY_After=ValueY_After)
        if (Debug==TRUE) points (ValueX_Random, ValueY_Random, col="green") 
        if (ValueY_Random > ValueY_After) 
          PosX_Random[Iter+1]  <- PosX_Random[Iter] +1 
        if (ValueY_Random > ValueY_Before) 
          PosX_Random[Iter+1] <- PosX_Random[Iter] - 1
        if (ValueY_Random <= ValueY_Before & ValueY_Random <= ValueY_After) 
          OptimProcess <- "End"
        Iter <- Iter+1
      }
    }
    ValueX_Valley <- Df$ValueX_Random
    ValueY_Valley <- Df$ValueY_Random
    Df <- data.frame(ValueX_Valley=ValueX_Valley, ValueY_Valley=ValueY_Valley)
    ResultsValleys <- rbind.data.frame (ResultsValleys, Df)
  }
  ResultsValleys <- unique (ResultsValleys)
  ResultsValleys <- ResultsValleys[order(ResultsValleys$ValueY_Valley, 
                                         decreasing = TRUE), ]
  # MAKE A GRAPHIC
  if (graphose==TRUE) {
    if (Debug==FALSE) {plot(DataX  , DataY , col="grey", cex=0.2 , 
                            xlab="X Value", ylab="Density" , 
                            main= "Density Graph" )}

    if (LineEachOptim == TRUE ) {
      abline(v=     ResultsPeaks$ValueX_Peak, lty=2, col="grey")
      abline(v= ResultsValleys$ValueX_Valley, lty=2, col="green")
    }
  }
  PeaksAndValleys <- list(Peaks=ResultsPeaks, Valleys=ResultsValleys)
  return( PeaksAndValleys)
}


#======================  Function IntensityCluster
# @name IntensityCluster
# @title Groups events in clusters based on their fluorescence intensities
# @description
# @details
# @param markerName Character string. Name of the fluorescent marker (e.g. RFP)
# @param markerColor Character string. Color to be used in the graphs for the
# fluorescent marker (e.g. red)
# @param eventsID Numeric. As a unique reference for the event, we take the
# time of the event as outputted by the cytometer. It is taken from
# DataSporeSingle$TIME
# @param eventsIntensity Numeric. Fluorescence intensity of the event in
# log arbitrary units, as outputted in DataSporeSingle$RFPHeightLog
# @param graphose Logical. If TRUE, produces graphs of data and results
# @param nbFluoClasses Numeric. Number of expected fluorescence classes
# @param cutOff Numeric. Minimum ratio between the height of a fluorescence 
# peak and that of the highest peak of the considered marker, to consider 
# the peak as significant. \cr
# Candidate peaks are discarded if Y <= cutOff*max(Y)
# @return List with elements:
#     $Log: Character string. status of the output
#     $AssignResults: Data frame with columns:
#        $eventsID Numeric, same as param eventsID
#        $eventsIntensity Numeric. Same as param eventsIntensity
#        $fluoClass Numeric.
#     $NumberPerClass Data frame with
#        $fluoClass Numeric. Number of the fluorescence class
#        $DataNumber Numeric. Number of events in the class
#     $HighestPeaksDf Data frame with:
#        $HighestPeakX
#        $HighestPeakY
#        $nbFluoClasses
IntensityCluster <- function(
              markerName, markerColor, eventsID, eventsIntensity,
              nbSeeds=100, graphose, nbFluoClasses, 
              cutOff=0.1, DataX, DataY)  {
  Data <- data.frame(eventsID=eventsID, eventsIntensity=eventsIntensity)
  #Find peaks and valleys
  intensityValues <- Data$eventsIntensity
  PeaksValleys <- FindPeaksAndValleys(intensityValues=intensityValues ,
                nbSeeds=100, graphose=FALSE, LineEachOptim=FALSE, Debug=FALSE)
  if (graphose==TRUE) {
    plot(DataX, DataY, cex=0.2, xlab="MarkerIntensity", ylab="Density", 
         main = markerName)
  }
  #Find the highest peaks sorted on X level, with at least a Y value > Ymax*cutOff
  Ymax <- max(PeaksValleys$Peaks$ValueY_Peak)
  HighestPeaks <- NULL
  #Keep only HighestPeak which Yvalue > Ymax*cutOff
  PeaksValleys$Peaks <- PeaksValleys$Peaks[
    which(PeaksValleys$Peaks$ValueY_Peak > Ymax*cutOff), ]
  for (fluoClass in 1:nbFluoClasses) {
    HighestPeakX <- PeaksValleys$Peaks[order((PeaksValleys$Peaks$ValueX_Peak)
                      , decreasing = TRUE), "ValueX_Peak"] [fluoClass]
    HighestPeakY <- PeaksValleys$Peaks[order((PeaksValleys$Peaks$ValueX_Peak)
                      , decreasing = TRUE), "ValueY_Peak"] [fluoClass]
    Df <- data.frame (HighestPeakX=HighestPeakX, HighestPeakY=HighestPeakY)
    HighestPeaks <- rbind.data.frame (HighestPeaks, Df)
  }
  #Give a number to each peak, sorted on X
  HighestPeaks <- HighestPeaks[order(HighestPeaks$HighestPeakX, 
                                     decreasing = TRUE), ]
  for (fluoClass in 1:nbFluoClasses) {
    HighestPeaks$fluoClass[fluoClass] <- fluoClass
  }
  #Send an error message if nbFluoClasses > than Number of finded peak, 
  #ie there is an NA value
  Status <- "Ok"
  for (i in 1:nbFluoClasses) {
    if (is.na(HighestPeaks$HighestPeakX[i]))
      Status="ERROR: Nb of peaks detected < nbFluoClasses"
  }
  if (Status=="Ok") {
    #TRACE PEAKS ON GRAPH
    Color <- c (markerColor, rep("grey",nbFluoClasses-1))
    if (graphose==TRUE) {
      abline(v=HighestPeaks$HighestPeakX, lty=1, col=Color )
      text(HighestPeaks$HighestPeakX, HighestPeaks$HighestPeakY*0.9, 
           labels=c(1:nbFluoClasses))
    }
    #Find valleys surrounding theses peaks
    HighestPeaks <- HighestPeaks[order(HighestPeaks$fluoClass, 
                                       decreasing=FALSE), ]
    SurroundingValleys= NULL
    for (fluoClass in 1:nbFluoClasses) {
      XLeft <- max(PeaksValleys$Valleys[
        which(PeaksValleys$Valleys$ValueX_Valley<HighestPeaks$HighestPeakX[fluoClass])
            , "ValueX_Valley" ] )
      XRight <- min(PeaksValleys$Valleys[
        which(PeaksValleys$Valleys$ValueX_Valley>HighestPeaks$HighestPeakX[fluoClass])
            , "ValueX_Valley" ]    )
      Df <- data.frame (fluoClass=fluoClass, XLeft=XLeft, XRight=XRight)
      SurroundingValleys <- rbind.data.frame (SurroundingValleys, Df)
    }
    HighestPeaks <- merge(HighestPeaks, SurroundingValleys, by="fluoClass")
    if (graphose==TRUE) abline(v=HighestPeaks$XLeft, lty=3, col="grey" )
    if (graphose==TRUE) abline(v=HighestPeaks$XRight, lty=3, col="grey" )
    #Assign a class to events
    Data$fluoClass <- "NA"
    for (fluoClass in 1:nbFluoClasses) {
      Data[(Data$eventsIntensity > HighestPeaks$XLeft[fluoClass])
           & (Data$eventsIntensity < HighestPeaks$XRight[fluoClass])
           , "fluoClass"] <- fluoClass
    }
    # Number of spores in each class
    NumberPerClass=NULL
    for (fluoClass in 1:nbFluoClasses) {
      DataNumber <- length(Data[which(Data$fluoClass==fluoClass), "eventsID"] )
      Df <- data.frame (fluoClass=fluoClass, DataNumber=DataNumber)
      NumberPerClass <- rbind.data.frame(NumberPerClass, Df)
    }
    EventsNonAssignNumber <- length(which(Data$fluoClass=="NA"))
    Df <- data.frame (fluoClass="NA", DataNumber=EventsNonAssignNumber)
    NumberPerClass <- rbind.data.frame (NumberPerClass, Df)
    if (graphose==TRUE) {
      legend("topleft", legend= paste("Class ", NumberPerClass$nbFluoClasses
            ," = ", NumberPerClass$DataNumber, sep="")    , text.col="black"
            , box.lty=0, bg="transparent" ) }
    ret = list(Log=Status,
               AssignResults=Data,
               NumberPerClass=NumberPerClass,
               HighestPeaksDf=HighestPeaks)
  }
  if (Status != "Ok") {
    ret <- list(    Log="ERROR: Nb of peaks detected < nbFluoClasses",
                 AssignResults=NA, NumberPerClass=NA, HighestPeaksDf=NA)
  }
  return( ret)
}


# @name CleanFcs
# @title Cleans cytometry data from invalid values
# @description First removes duplicated events, and then removes all
# events with non-numeric log-values of the relevant channels (FSC.H, SSC.H,
# SSC.A, fluo1.H, fluo2.H, fluo3.H)
# @param fcsData Data frame. As outputted by function ImportFcs()
# @param logFile Character string. File and path name for the log file
# @return Data frame. As outputted by function ImportFcs()
CleanFcs <- function(fcsData, fluo1H, fluo2H, fluo3H,
                     chanFscH, chanSscH, chanSscA, logFile=NULL) {
  initSize <- dim(fcsData)[1]
  fcsData <- fcsData[ ! duplicated(fcsData$Time),]
  noDupSize <- dim(fcsData)[1]
  if (length(duplicated(fcsData$Time))>0) {
    msg <- paste0("Removed ",initSize-noDupSize
                  ," duplicated event IDs (TIME)")
    cat(paste0(msg,"\n"))
    if (!is.null(logFile)) write(msg, file=logFile, append=TRUE)
  }

  fcsData <- fcsData[! is.na(fcsData[, chanFscH]), ]
  fcsData <- fcsData[! is.na(fcsData[, chanSscH]), ]
  fcsData <- fcsData[! is.na(fcsData[, chanSscA]), ]
  fcsData <- fcsData[! is.na(fcsData[, fluo1H[2]]), ]
  fcsData <- fcsData[! is.na(fcsData[, fluo2H[2]]), ]
  fcsData <- fcsData[! is.na(fcsData[, fluo3H[2]]), ]

  fcsData <- fcsData[is.numeric(fcsData[, chanFscH]), ]
  fcsData <- fcsData[is.numeric(fcsData[, chanSscH]), ]
  fcsData <- fcsData[is.numeric(fcsData[, chanSscA]), ]
  fcsData <- fcsData[is.numeric(fcsData[, fluo1H[2]]), ]
  fcsData <- fcsData[is.numeric(fcsData[, fluo2H[2]]), ]
  fcsData <- fcsData[is.numeric(fcsData[, fluo3H[2]]), ]

  fcsData <- fcsData[is.finite(fcsData[, chanFscH]), ]
  fcsData <- fcsData[is.finite(fcsData[, chanSscH]), ]
  fcsData <- fcsData[is.finite(fcsData[, chanSscA]), ]
  fcsData <- fcsData[is.finite(fcsData[, fluo1H[2]]), ]
  fcsData <- fcsData[is.finite(fcsData[, fluo2H[2]]), ]
  fcsData <- fcsData[is.finite(fcsData[, fluo3H[2]]), ]

  fcsData <- fcsData[fcsData[, chanFscH] > 0, ]
  fcsData <- fcsData[fcsData[, chanSscH] > 0, ]
  fcsData <- fcsData[fcsData[, chanSscA] > 0, ]
  fcsData <- fcsData[fcsData[, fluo1H[2]] > 0, ]
  fcsData <- fcsData[fcsData[, fluo2H[2]] > 0, ]
  fcsData <- fcsData[fcsData[, fluo3H[2]] > 0, ]
  endSize <- dim(fcsData)[1]
  if (noDupSize-endSize>0) {
    msg <- paste0("Removed ",noDupSize-endSize
                  ," invalid raw data in channels")
    cat(paste0(msg,"\n"))
    if (!is.null(logFile)) write(msg, file=logFile, append=TRUE)
  }
  return(fcsData)
}


# @name ImportFcs
# @title
# @description Import FCS data from MoFlo or CytExpert
# @param fcsDataPath path to Fcs files (.fcs)
# @param truncateMaxRange if you don't want to import all data
ImportFcs <- function(fcsDataPath, fluo1H, fluo2H, fluo3H,
                      chanFscH, chanSscH, chanSscA,
                      truncateMaxRange, logFile) {
  
  if (!file.exists(fcsDataPath)) stop(paste0(" Cytometry file not found at: ", fcsDataPath))
  
  RawData = tryCatch(
    suppressWarnings(
      flowCore::read.FCS(fcsDataPath, alter.names=TRUE, truncate_max_range=truncateMaxRange)
    ),
    error = function(err) err
  )
  # When an exception is raised, the class of RawData is "error" and contains the error message.
  if(is(RawData,"error")) {
    stop('Reading this FCS file failed: "',fcsDataPath,'"\n', RawData$message, domain=NA)
  }
  
  kw <- flowCore::keyword(RawData)
  cyt <- kw[['$CYT']]
  Data <- as.data.frame(flowCore::exprs(RawData))
  
  if(is.null(Data$Time)) Data$Time <- Data$TIME
  if(is.null(Data$TIME)) Data$TIME <- Data$Time
  
  if( any(c(fluo1H[2], fluo2H[2], fluo3H[2], chanFscH, chanSscH, chanSscA) == "auto") ){
    if(!is.null(cyt) && cyt == "MoFlo Astrios"){
      FcsType <- "MoFlo Astrios"
      if (fluo1H[2]=="auto") fluo1H[2] <- "FL8.Height"
      if (fluo2H[2]=="auto") fluo2H[2] <- "FL20.Height"
      if (fluo3H[2]=="auto") fluo3H[2] <- "FL17.Height"
      if (chanFscH=="auto") chanFscH <- "FSC1.Height"
      if (chanSscH=="auto") chanSscH <- "SSC2.Height"
      if (chanSscA=="auto") chanSscA <- "SSC2.Area"
      Data$Time <- Data$TIME
    } else if (!is.null(cyt) && cyt == "CytoFLEX S") {
      if("FL8.H" %in% colnames(Data)){
        FcsType <- "CytoFLEX after 2021"
        if (fluo1H[2]=="auto") fluo1H[2] <- "FL1.H"
        if (fluo2H[2]=="auto") fluo2H[2] <- "FL8.H"
        if (fluo3H[2]=="auto") fluo3H[2] <- "FL5.H"
        if (chanFscH=="auto") chanFscH <- "FSC.H"
        if (chanSscH=="auto") chanSscH <- "SSC.H"
        if (chanSscA=="auto") chanSscA <- "SSC.A"
        Data$TIME <- Data$Time
      } else if("FL6.H" %in% colnames(Data)){
        FcsType <- "CytoFLEX before 2021"
        if (fluo1H[2]=="auto") fluo1H[2] <- "FL6.H"
        if (fluo2H[2]=="auto") fluo2H[2] <- "FL10.H"
        if (fluo3H[2]=="auto") fluo3H[2] <- "FL1.H"
        if (chanFscH=="auto") chanFscH <- "FSC.H"
        if (chanSscH=="auto") chanSscH <- "SSC.H"
        if (chanSscA=="auto") chanSscA <- "SSC.A"
        Data$TIME <- Data$Time
      } else { # unknown CytoFLEX type
        msg <- "Unknown format of CytoFLEX data."
        stop(msg, domain=NA)
      }
    } else { # cyt is null or unknown cytometer name.
      msg <- "Unknown cytometer type.\nPlease specify manually all channel names."
      stop(msg, domain=NA)
    }
  } else {
    msg <- "Info: Channels names manually set by user."
    cat(paste0(msg,"\n"))
    if (!is.null(logFile)) write(msg, file=logFile, append=TRUE)
    if (!is.null(cyt)) { FcsType <- cyt } else FcsType <- "Unknown"
  }

  msg <- paste0("Type of cytometer detected: ", FcsType)
  cat(paste0(msg,"\n"))
  if (!is.null(logFile)) write(msg, file=logFile, append=TRUE)
  
  Data <- CleanFcs(fcsData=Data, fluo1H, fluo2H, fluo3H,
                   chanFscH, chanSscH, chanSscA, logFile=logFile)
  
  Data$FSC1HeightLog <- log10(Data[, chanFscH])
  Data$SSC1HeightLog <- log10(Data[, chanSscH])
  Data$SSC1AreaLog   <- log10(Data[, chanSscA])
  Data$CFPHeightLog <- log10(Data[, fluo1H[2]])  # CFP
  Data$RFPHeightLog <- log10(Data[, fluo2H[2]])  # RFP
  Data$YFPHeightLog <- log10(Data[, fluo3H[2]])  # YFP
  
  msg <- paste0("Valid events imported: ", nrow(Data))
  cat(paste0(msg,"\n"))
  if (!is.null(logFile)) write(msg, file=logFile, append=TRUE)
  
  FcsData <- list(Data=Data, StatusFcs="OK", FcsType=FcsType,
                  fluo1H, fluo2H, fluo3H, chanFscH, chanSscH, chanSscA)
  return(FcsData)
}


# @name ParsePlateDesign
# @title Parse Plate Design file
# @description Reads plate design file, adds two additional columns 'Status'
# and 'Changed_Params' (initially filled with NAs), and saves to the same file
# name, but extended with "_PARAMS" before the ".txt" extension. This new file
# may then be edited manually to replace the NA in the column 'Changed_Params'
# by a character string indicating specific values that should be given to
# some parameters of the function AnalyzeRecombination() for that specific 
# sample. For instance, if the 3rd sample of the plate design file must have
# the specific values 0.3 for parameter 'cutOff' and 1.1 for parameter
# 'widthAroundPeak', the string "cutOff=0.3_widthAroundPeak=1.1" must be put
# at the 3rd line and column 'Changed_Params'. \cr
# Then, if this xxx_PARAMS.txt file exists in the plate design file directory,
# the particular parameters values indicated in its column 'Changed_Params' will 
# be used only for the corresponding sample by the function 
# AnalyzeRecombination().
# @param plateDesignFile Character string. Name with complete path of the
# plate design file
# @param logFile Character string. Name with complete path of the log file
ParsePlateDesign <- function(plateDesignFile, logFile) {

  if (length(grep(x=plateDesignFile, pattern="\\_PARAMS.txt$"))==0) {
    plateParamsFile <- sub(x=plateDesignFile, pattern="\\.txt$"
                           , replacement="_PARAMS.txt")
  } else plateParamsFile <- plateDesignFile
  if ( ! file.exists(plateParamsFile)) {
    plateDesign <- read.table (plateDesignFile, sep="\t", header=TRUE
                               , blank.lines.skip=TRUE, skipNul=T)
    colnames(plateDesign) <- c("Sample","fcs_ID")
    Re_Do <- rep(1, nrow(plateDesign))
    Changed_Params <- rep(NA, nrow(plateDesign))
    plateParams <- cbind.data.frame(plateDesign, Re_Do, Changed_Params)
    write.table(x=plateParams, file=plateParamsFile, sep="\t",quote=F, row.names=F)
  } else {
    plateParams <- read.table(file=plateParamsFile, sep="\t", header=TRUE
                              , blank.lines.skip=TRUE, skipNul=T)
  }
#  plateParams <- plateParams[plateParams$Re_Do!=0,]
  plateParams <- plateParams[plateParams$Sample!="" 
                             & plateParams$fcs_ID!="",]
    
  parStrings <- c("outDir","fcsDir","plateDesignFile","pdfGraphs")
  parNums <- c("minNbSpores","nbUsedForGating",
    "minSporeFscHLog","maxSporeFscHLog","minSporeSscHLog","maxSporeSscHLog=6",
    "cutOff", "binNumberX","binNumberY","slopeFindPeak","widthAroundPeak")
  parVecs <- c("tokensDateChrMarkers","fluo1H","fluo2H","fluo3H",
               "chanFscH","chanSscH","chanSscA")
  changedParams <- list()
  iSamp <- 1
  for (iSamp in 1:nrow(plateParams)) {
    samplePars <- NULL
    if ( ! is.na(plateParams[iSamp,]$Changed_Params)) {
      parStr <- plateParams[iSamp,]$Changed_Params
      parVec <- unlist(strsplit(parStr,"_"))
      iPar <- 1
      for (iPar in 1:length(parVec)) {
        parName <- unlist(strsplit(parVec[iPar],"="))[1]
        parVal <- unlist(strsplit(parVec[iPar],"="))[2]
        if (parName %in% parNums) {
          parVal <- as.numeric(parVal)
          df <- data.frame(parName=parName, parVal=parVal)
          samplePars <- rbind.data.frame(samplePars, df)
        } else if (parName %in% parStrings) {
          msg <- paste0("\nError in reading changed parameter ", parName
                        ,"\nThis parameter (string) cannot ",
                        "be changed in file ",plateParamsFile)
          cat(msg) ; write(msg, file=logFile, append=TRUE)
        } else if (parName %in% parVecs) {
          msg <- paste0("\nError in reading changed parameter ", parName
                        ,"\nThis parameter (vector) cannot ",
                        "be changed in file ",plateParamsFile)
          cat(msg) ; write(msg, file=logFile, append=TRUE)
        } else {
          msg <- paste0("\nError in reading changed parameter ", parName
                    ,"\nThis is not a valid parameter of AnalyzeRecombination()")
          cat(msg) ; write(msg, file=logFile, append=TRUE)
        }
      }
    }
    changedParams <- c(changedParams, list(samplePars))
  }
  return(list(plateDesign=plateParams, lst=changedParams
              , str=plateParams$Changed_Params))
}


#' @name AnalyzeRecombination
#' @title Automatic gating and calculation of recombination rates and CoC
#' @description This functions reads a plate design file containing a list of 
#' samples, and then for each of them, reads the corresponding FCS output file 
#' from the cytometer, sets automatically the gates on fluorescence values 
#' to count yeast spores according to their phenotype for two or three
#' fluorescent markers, and analyses segregation to compute recombination
#' fractions between markers as well as (in the case of three markers) the
#' coefficient of coincidence (CoC) which indicates the effect of crossover
#' interference. The markers must be hemizygous in the parental diploids, 
#' located on the same chromosome, and genetically linked 
#' (see Raffoux et al. 2018 Yeast(6):431). Based on the same paper, the 
#' function also infers recombination and interference 
#' parameters by Maximum-Likelihood based on a model including partial 
#' fluorescence extinction of some of the markers.
#' @param outDir Character string. Path to the folder where all output folders
#' and files will be written
#' @param fcsDir Character string. Path to the folder containing raw data
#' files (.fcs) from the flow cytometer
#' @param plateDesignFile Character string. Name (must end with ".txt") with
#' complete path of the file containing the sample list (or plate design). 
#' This tab-delimited text file (with headers) must have the following two 
#' columns:
#' \itemize{
#'   \item 'Sample' containing the name of the sample. This name should be made 
#'   of several tokens separated by underscores. This will be used to collect
#'   mandatory information concerning the date of the experiment, the 
#'   chromosome, and the fluorescent markers present in the sample with their 
#'   order along the chromosome. The respective positions of these tokens in 
#'   the sample name can be specified in the argument 'tokensDateChrMarkers'. 
#'   For instance, for the sample name 'VAR-PLATE-2B_VI_C1Y2R3_20230502', 
#'   corresponding to chromosome VI, date 2023/05/02, and markers 'C', 'Y', 
#'   and 'Y' in that order on the chromosome, the parameter 
#'   'tokensDateChrMarkers' must be set to c(4,2,3). 
#'   \item 'fcs_ID' containing a unique alphanumeric sample ID which must 
#'   also be present at the end of the FCS file just before the '.fcs' 
#'   extension, and separated from the beginning of the file name by '_' 
#'   or '-' (e.g. the ID 'A1' will be in the FCS file 'Exp1_A1.fcs'). 
#'   This ID can (but does not have to) be for instance the well coordinates 
#'   in the form "A1".."H12".
#'  }
#'  When the file 'plateDesignFile' is read for the first time, CAYSS will
#'  write a file containing the same columns with two additional ones:
#' \itemize{
#'   \item 'Re_Do' initially filled with '1'. If some '1' are manually turned 
#'   to '0' in that file for some samples, if the same 'plateDesignFile' 
#'   is read another time to re-analyze the same data, then the samples 
#'   corresponding to the lines with Re_Do=='0' will be skipped.
#'   \item 'Changed_Params' initially filled with 'NA'. If some samples
#'   need to be re-analyzed with some different parameters, then in the column
#'   'Changed_Params' of the corresponding lines, the 'NA' can be replaced by 
#'   a character string indicating which parameters must be modified for that
#'   particular sample, and which new values they can take. For instance, if 
#'   'cutOff' must be turned to 0.3 and 'minNbSpores' must be turned to 800,
#'   then the column 'Changed_Params' must contain 'cutOff=0.3_minNbSpores=800'
#'   for that line.
#'  }
#' @param fluo1H Vector of Character strings. Named vector with elements:
#'  \itemize{
#'    \item $name: Character string. Name of the fluorescent marker (e.g. "Cerulean"
#'    or "CFP")
#'    \item $channel: Character string. Corresponding name of the channel of the
#'    cytometer (e.g. "FL1")
#'    \item $code: Character. Single-letter code of the marker (e.g. "C").
#'    This code must be used in the sample name of the plateDesign File column 
#'    to encode the order of the marker positions along the chromosome 
#'    (e.g. "CYR"), indicating how recombination rates should be calculated.
#'  }
#'@param fluo2H Vector of Character strings. Named vector with elements:
#'  \itemize{
#'    \item $name: Character string. Name of the fluorescent marker (e.g. 
#'    "mCherry" or "RFP")
#'    \item $channel: Character string. Corresponding name of the channel of the
#'    cytometer (e.g. "FL8")
#'    \item $code: Character. Single-letter code of the marker (e.g. "r").
#'    This code must be used in the sample name of the plateDesign File column 
#'    to encode the order of the marker positions along the chromosome 
#'    (e.g. "CYR"), indicating how recombination rates should be calculated.
#'  }
#'@param fluo3H Vector of Character strings. Named vector with elements:
#'  \itemize{
#'    \item $name: Character string. Name of the fluorescent marker (e.g. 
#'    "Venus" or "YFP")
#'    \item $channel: Character string. Corresponding name of the channel of the
#'    cytometer (e.g. "FL5")
#'    \item $code: Character. Single-letter code of the marker (e.g. "y").
#'    This code must be used in the sample name of the plateDesign File column 
#'    to encode the order of the marker positions along the chromosome 
#'    (e.g. "CYR"), indicating how recombination rates should be calculated.
#'  }
#' @param chanFscH Character string. Name of the output channel of the
#' cytometer, that should be used to measure the height of Forward SCattering
#' (FSC) peaks, indicating the size of the objects e.g. "FSC.H"
#' @param chanSscH Character string. Name of the output channel of the
#' cytometer, that should be used to measure the height of Side SCattering
#' (SSC) peaks, indicating the surface texture of the objects e.g. "SSC.H"
#' @param chanSscA Character string. Name of the output channel of the
#' cytometer, that should be used to measure the area under the time-curve of
#' Side SCattering (SSC) peaks, indicating the surface texture of the objects
#' e.g. "SSC.A"
#' @param minNbSpores Numeric. Minimum number of spores singlets to continue
#' the analysis. Otherwise skip that sample.
#' @param nbUsedForGating Numeric or Character string.
#' - If Numeric: Number of spores (events that have been previously selected 
#' as Spores and Singlets) used to automatically determine the boundaries of 
#' gates to be applied to fluorescence values. This is to reduce computation 
#' time, while all data will be used to calculate recombination rates. 
#' - If "all" then all spores are used to determine the boundaries of gates.
#' @param minSporeFscHLog Numeric. Minimum threshold of FSC (Forward SCatter) 
#' peak height value (log scale) to preselect events as being putative spores
#' @param maxSporeFscHLog Numeric. Maximum threshold of FSC (Forward SCatter) 
#' peak height value (log scale) to preselect events as being putative spores
#' @param minSporeSscHLog Numeric. Minimum threshold of SSC (Side SCatter) 
#' peak height value (log scale) to preselect events as being putative spores
#' @param maxSporeSscHLog Numeric. Maximum threshold of SSC (Side SCatter) 
#' peak height value (log scale) to preselect events as being putative spores
#' @param cutOff Numeric. Minimum ratio between the height of a fluorescence 
# peak and that of the highest peak for the considered marker, to consider 
# that peak as significant.
#' @param binNumberX Numeric. Number of bins for X scans: size of the grid 
#' used to locate density peaks
#' @param binNumberY Numeric. Number of bins for Y scans: size of the grid 
#' used to locate density peaks
#' @param slopeFindPeak Numeric. Threshold value (between 0 and 1) used to
#' detect the extent of density peaks. When this value is higher, events in a 
#' wider region around the peak summit are considered as belonging to the peak. 
#' This is used at two levels: 
#' - to determine the (circular) gate to be applied to events in the graph
#'   SporeSscHLog vs SporeFscHLog, to further select spores after 
#'   pre-selecting events based on 'minSporeFscHLog', 'maxSporeFscHLog', 
#'   'minSporeSscHLog', and 'maxSporeSscHLog'
#' - to determine the (linear) gate to be applied to events in the graph
#'   SporeSscHeightLog vs SporeFscAreaLog to select the spore singlets from 
#'   possible remaining of spore doublets if the spore preparation protocol
#'   was not efficient enough
#' @param widthAroundPeak Numeric. Factor applied to the standard deviation 
#' of a peak, to determine the boundaries for events that should be assigned 
#' to that fluo class. This is used to identify the fluorescence phenotype of 
#' spores. 
#' @param tokensDateChrMarkers Vector of Numerics. Token position (tokens are
#' separated by underscores) in the sample names listed in the Plate Design 
#' file, for Date, Chromosome, and Markers, respectively. For instance, 
#' tokensDateChrMarkers will be set to c(5,3,4) if the sample name has the form
#' Exp1_Validation_ChrIV_CRY_20240805 and the order of the fluorescent markers 
#' on the chromosome is C-R-Y
#' @param pdfGraphs Character string. Determines how many pdf files will be
#' generated with the analysis output graphs:
#' - if "separated", a distinct pdf file will be generated for each sample
#' - if "merged_only", a single multi-page pdf file will be generated with all 
#' samples
#' - if "both", all single pdf files will be kept, and also the multi-page 
#' merged file will be generated and named "Date_Time_ALL_CYTO_GRAPHS.pdf"
#' - if "none", no pdf files will be written for the cytometry graphs, and 
#' the graphics will be sent to the default graphical device
#' @return Data.frame with columns:
#' \itemize{
#'   \item $SampleName Character string. Complete sample name merged with the
#'   fcs_ID found in the Plate Design file
#'   \item $DateExp Character string. Date, taken from the Plate Design file 
#'    (token 'tokensDateChrMarkers\[1\]')
#'   \item $Chr Character string. Chromosome name, taken from the Plate Design 
#'   file (token 'tokensDateChrMarkers\[2\]')
#'   \item $Markers Character string. Marker order, taken from the Plate Design 
#'   file (token 'tokensDateChrMarkers\[3\]'). E.g. "CYR"
#'   \item $NbEvents Numeric. Total number of valid events read from the 
#'   FCS file
#'   \item $NbPreselected Numeric. Number of events pre-selected based on 
#'   thresholds 'minSporeFscHLog', 'maxSporeFscHLog', 'minSporeSscHLog', and 
#'    'maxSporeSscHLog'
#'   \item $NbSpores Numeric. Number of events considered as being spores, 
#'   after pre-selection based on thresholds (see below) and gating of the main 
#'   peak based on 'slopeFindPeak'
#'   \item $NbSingletSpores Numeric. Number of events considered as spores 
#'   (see below) and being considered as singlets based on selection with a 
#'   linear gate in the graph SporeSscHeightLog vs SporeFscAreaLog
#'   \item $Freq_NameFluo1 Numeric. Frequency of singlet spores expressing the 
#'   fluo1 marker. In this column name, NameFluo1 will be replaced by the name 
#'   of the 1st fluorescent marker, as entered in 'fluo1H\["name"\]'.
#'   \item $Freq_NameFluo2 Numeric. Frequency of singlet spores expressing the 
#'   fluo2 marker. In this column name, NameFluo2 will be replaced by the name 
#'   of the 2nd fluorescent marker, as entered in 'fluo2H\["name"\]'.
#'   \item $Freq_NameFluo3 Numeric. Frequency of singlet spores expressing the 
#'   fluo3 marker. In this column name, NameFluo3 will be replaced by the name 
#'   of the 3rd fluorescent marker, as entered in 'fluo3H\["name"\]'.
#'   \item $FreqABCvsPPP Numeric. Ratio between the frequency of spores 
#'   expressing markers A, B, and C (A, B, and C indicate the three fluorescent 
#'   markers in the order of their position on the chromosome) and the 
#'   frequency of non-fluorescent spores
#'   \item $FreqBCvsA Numeric. Ratio between the frequency of spores expressing 
#'   only markers B and C and the frequency of spores expressing only marker A
#'   \item $FreqABvsC Numeric. Ratio between the frequency of spores expressing 
#'   only markers A and B and the frequency of spores expressing only marker C
#'   \item $FreqACvsB Numeric. Ratio between the frequency of spores expressing 
#'   only markers A and C and the frequency of spores expressing only marker B
#'   \item $Naive_rAB Numeric. Recombination fraction between markers A and B, 
#'   calculated as (fA+fAC+fB+fBC), fX being the frequency of spores expressing 
#'   the fluorescent phenotype X
#'   \item $Naive_rBC Numeric. Recombination fraction between markers B and C, 
#'   calculated as (fB+fAB+fC+fAC), fX being the frequency of spores expressing 
#'   the fluorescent phenotype X
#'   \item $Naive_rDouble Numeric. Fraction of double recombinant spores,
#'   calculated as (fAC+fB), fX being the frequency of spores expressing the
#'   fluorescent phenotype X
#'   \item $Naive_CoC Numeric. Coefficient of coincidence used to assess 
#'   crossover interference between intervals AB and BC, calculated as 
#'   (fAC+fB)/((fA+fAC+fB+fBC)*(fB+fAB+fC+fAC)). CoC equals 1 if there is no
#'   interference, and CoC becomes lower as interference becomes stronger
#'   \item $StatusFcs Character string. Status of reading the FCS file. Can 
#'   return 'ok' or 'Not enough events in FCS file'
#'   \item $StatusGating Character string. Status of the automatic gating 
#'   procedure. Can return 'Not enough preselected events before spore gating', 
#'   'Not enough gated spores', 'Not enough single spores', 'Only one Fluo 
#'    peak detected', or 'Data number ok'
#'   \item $App Numeric. Number of spores expressing only fluo marker A
#'   \item $ABp Numeric. Number of spores expressing only fluo markers A and B
#'   \item $ABC Numeric. Number of spores expressing fluo markers A, B, and C
#'   \item $ApC Numeric. Number of spores expressing only fluo markers A and C
#'   \item $pBp Numeric. Number of spores expressing only fluo marker B
#'   \item $pBC Numeric. Number of spores expressing only fluo markers B and C
#'   \item $ppC Numeric. Number of spores expressing only fluo marker C
#'   \item $ppp Numeric. Number of spores expressing non of the fluo markers
#'   \item $TotSpores Numeric. Total number of spores 
#'   \item $ML_rAB Numeric. Recombination fraction between markers A and B, 
#'   inferred by Maximum-Likelihood with a model including partial extinction
#'   of fluorescence for each of the markers
#'   \item $ML_rBC Numeric. Recombination fraction between markers B and C, 
#'   inferred by Maximum-Likelihood with a model including partial extinction
#'   of fluorescence for each of the markers
#'   \item $ML_CoC Numeric. Coefficient of coincidence inferred by 
#'   Maximum-Likelihood with a model including partial extinction of 
#'   fluorescence for each of the markers
#'   \item $ML_Aext Numeric. Frequency of fluorescence extinction for marker A, 
#'   inferred by Maximum-Likelihood
#'   \item $ML_Bext Numeric. Frequency of fluorescence extinction for marker B, 
#'   inferred by Maximum-Likelihood
#'   \item $ML_Cext Numeric. Frequency of fluorescence extinction for marker C, 
#'   inferred by Maximum-Likelihood
#'   \item $StatusML Character string. Status of the ML estimation of parameters
#'   rAB, rBC, Coc, extA, extB, and extC. Possible values:
#'    - "ok" if the ML inference converged correctly and the results are close
#'    to the naive estimates
#'    - "Warning: Naive-ML difference > 10%" if the inference converged but the 
#'    results are quite different from the naive estimates. This may
#'    indicate that fluorescence extinction is significant on some of the 
#'    markers, or that the gating was not accurate (e.g. if there are 
#'    unexpected fluo peaks)
#'    - "ML failed to converge" if the observed numbers of spores of each
#'    fluorescent phenotype did not allow the model to converge
#'   \item $ChangedParams Character string or NA. 
#'   - if NA, the sample was run with all
#'   parameters used to call the function AnalyzeRecombination()
#'   - if character string, list of parameters that have been given specific 
#'   values for a particular sample, separated by an underscore 
#'   (e.g. 'cutOff=0.2_widthAroundPeak=1.0')
#'  }
#' @examples
#' plate <- ExampleFile("PlateDesign2")
#' fcs <- system.file("extdata", package="CAYSS")
#' out <- getwd()
#' res <- AnalyzeRecombination(
#'    outDir=out, 
#'    fcsDir=fcs, 
#'    plateDesignFile=plate, 
#'    tokensDateChrMarkers=c(2,3,4)
#'  ) 
#' @export
AnalyzeRecombination <- function (
      outDir, fcsDir, plateDesignFile,
      fluo1H=c(name="CFP",channel="auto",code="C"),
      fluo2H=c(name="RFP",channel="auto",code="R"),
      fluo3H=c(name="YFP",channel="auto",code="Y"),
      chanFscH="auto", chanSscH="auto", chanSscA="auto",
      minNbSpores=1000, nbUsedForGating=5000,
      minSporeFscHLog=4.9, maxSporeFscHLog=5.3, 
      minSporeSscHLog=0,   maxSporeSscHLog=6,
      cutOff=0.1, binNumberX=100, binNumberY=100, 
      slopeFindPeak=0.4, widthAroundPeak=1.6,
      tokensDateChrMarkers=c(1,2,3), pdfGraphs="both") {

  argsList <- c(as.list(environment()))
  firstSelectOnFSC <- FALSE
  discardNFSpores <- TRUE
  halfDistrib <- FALSE
  savNbUsedForGating <- nbUsedForGating
  minFluo1Value=0 ; minFluo2Value=0 ; minFluo3Value=0
  
  requireNamespace("stringr")
  requireNamespace("flowCore")
  setwd(outDir)
  allCytoPdfFiles <- allMlPdfFiles <- c()

## 10 Create output folders to save graphic, results, and log file----
  Date <- Sys.Date()
  Time <-  stringr::str_replace_all(format(Sys.time(), "%X"), ":", "")
  PathCytoGraphs  <- paste0(Date,"_",Time,"_GRAPHICS")
  PathResults <- paste0(Date,"_",Time,"_RESULTS")
  PathMlGraphs <- paste0(Date,"_",Time,"_MAX_LIKELIHOOD")
  if ( ! file.exists(PathCytoGraphs)) dir.create(PathCytoGraphs)
  if ( ! file.exists(PathResults)) dir.create(PathResults)
  if ( ! file.exists(PathMlGraphs)) dir.create(PathMlGraphs)
  log <- "ARGUMENTS:\n"
  if (length(argsList)>0) for (iArg in 1:length(argsList)) {
    log <- paste0(log, names(argsList)[iArg], ": ",
                  paste(argsList[[iArg]], collapse=" "), "\n")
  }
  logFile <- file.path(PathResults, paste0(Date,"_",Time,"_LOG_FILE.txt"))
  write(log, file=logFile)

  AllResults=NULL

## 15 Read PlateDesign and ChangedParams file----
  
  changedParams <- ParsePlateDesign(plateDesignFile=plateDesignFile
                                    , logFile=logFile)
  changedParStrs <- changedParams$str
  plateDesign <- changedParams$plateDesign
  write.table(x=plateDesign
              , file=file.path(PathResults, "Sample_List_PARAMS.txt")
              , sep="\t",quote=F, row.names=F)
  
  allFcsFiles <- unique(list.files(fcsDir, pattern=".fcs"))
  allCoords <- sub(x=allFcsFiles, pattern=".*[-_ ]([^-_ ]+)\\.fcs", replacement="\\1")
  
## 20 LOOP on samples =======================----
  iFcs <- 5
  for (iFcs in 1:dim(plateDesign)[1]) {   ##  MAIN LOOP  ##
    
    nbUsedForGating <- savNbUsedForGating
    if (plateDesign[iFcs,]$Re_Do==0) next
    
    Continue <- TRUE
    set.seed(111)
    sampleName <- plateDesign[iFcs,]$Sample
    plateCoord <- plateDesign[iFcs,]$fcs_ID
    sampleName <- paste0(sampleName,"_", plateCoord)

## 25 Initialize Graphics window----
    cytoPdfFile <- file.path(PathCytoGraphs, paste0(sampleName,".pdf"))
    if (pdfGraphs != "none") {
      pdf(file=cytoPdfFile, width = 21/2.54, height = 29.7/2.54)
      par(mfrow=c(3,2))
    }
    dev1 <- dev.cur()
    mlPdfFile <- file.path(PathMlGraphs, paste0(sampleName,"_Max_Likelihood_Fit.pdf"))

## 30 Parse sample information----
    vec <- unlist(strsplit(sampleName,"_"))
    expDate <- vec[tokensDateChrMarkers[1]]
    expDate <- as.Date(expDate,
      tryFormats=c("%d/%m/%y%y","%d-%m-%y%y","%d%m%y%y","%y%y%m%d","%y%y-%m-%d",""))
    Chr <- vec[tokensDateChrMarkers[2]]
    Markers <- vec[tokensDateChrMarkers[3]]
#    sample <- paste(vec[-tokensDateChrMarkers], collapse="_")
    a <- unlist(strsplit( Markers ,""))		#Get marker order
    suppressWarnings(b <- as.numeric(a))
    a <- a[is.na(b)]
    if (length(a)%in%c(2,3)
        && sum(a %in% c(fluo1H[3],fluo2H[3],fluo3H[3]))%in%c(2,3)) {
      a[a==fluo1H[3]] <- "C"
      a[a==fluo2H[3]] <- "R"
      a[a==fluo3H[3]] <- "Y"
      Order <- paste(a, collapse = "")
    } else stop(paste0(" Invalid 'Markers' token: ",Markers))

    iFile <- which(allCoords==plateCoord)
    if (length(iFile)>1) stop(paste0(
            " Two different samples have the coordinate: ", plateCoord))
    if (length(iFile)<1) {
      msg <- paste0("No FCS file found for sample ",sampleName, " Skipping")
      cat(paste0(msg,"\n")) ; write(msg, file=logFile, append=TRUE)
      dev.off()
      next
    } else {
      msg <- paste0(sampleName," => ",allFcsFiles[iFile])
      cat(paste0(msg,"\n")) ; write(msg, file=logFile, append=TRUE)
    }
    
## 35 Initializes output data frame Df----
    res <- data.frame (
      SampleName=sampleName, DateExp=expDate, Chr=Chr, Markers=Markers, 
      NbEvents=NA, NbPreselected=NA, NbSpores=NA, NbSingletSpores=NA, 
      FreqCFP=NA, FreqRFP=NA, FreqYFP=NA,
      FreqABCvsPPP=NA, FreqBCvsA=NA, FreqABvsC=NA, FreqACvsB=NA,
      Naive_rAB=NA, Naive_rBC=NA, Naive_rDouble=NA, Naive_CoC=NA, 
      StatusFcs=NA, StatusGating=NA,
      App=NA, ABp=NA, ABC=NA, ApC=NA, pBp=NA, pBC=NA, ppC=NA, ppp=NA,
      TotSpores=NA, ML_rAB=NA, ML_rBC=NA, ML_CoC=NA, ML_Aext=NA, ML_Bext=NA,
      ML_Cext=NA, StatusML=NA, ChangedParams=NA)
    res$ChangedParams <- changedParStrs[iFcs]
    colnames(res)[colnames(res)=="FreqCFP"] <- paste0("Freq_",fluo1H[1])
    colnames(res)[colnames(res)=="FreqRFP"] <- paste0("Freq_",fluo2H[1])
    colnames(res)[colnames(res)=="FreqYFP"] <- paste0("Freq_",fluo3H[1])
    
## 38 Changes params specified in plate design file----
    pars <- changedParams$lst[[iFcs]]
    if ( ! is.null(pars)) {
      for (iPar in 1:length(pars)) {
        assign(x=pars[iPar,]$parName, value=pars[iPar,]$parVal)
        msg <- paste0("Changed param '",
                      pars[iPar,]$parName,"' to ",pars[iPar,]$parVal)
        cat(paste0(msg,"\n")) ; write(msg, file=logFile, append=TRUE)
      }
    }
    
## 40 Import FCS Data----
    fcsDataPath <- file.path(fcsDir, allFcsFiles[iFile])
    Data <- ImportFcs(fcsDataPath=fcsDataPath,
                      fluo1H=fluo1H, fluo2H=fluo2H, fluo3H=fluo3H,
                      chanFscH=chanFscH, chanSscH=chanSscH, chanSscA=chanSscA,
                      truncateMaxRange=FALSE,
                      logFile=logFile)
    StatusFcs <- Data$StatusFcs
    Data <- Data$Data
    res$NbEvents <- nrow(Data)
    RawData <- Data
    if (nrow(RawData) < minNbSpores) {
      msg <- paste0("Nb of events loaded: ",nrow(RawData),
                    " < 'minNbSpores' (", minNbSpores,")")
      cat(paste0(msg,"\n")) ; write(msg, file=logFile, append=TRUE)
      res$StatusFcs <- "Not enough events in FCS file"
      Continue <- FALSE
    }

## 45 Sub-sample nbUsedForGating events----
    if (Continue && nrow(RawData) > nbUsedForGating) {
      RawDataSample <- RawData[sample(nrow(RawData), nbUsedForGating), ]
    } else RawDataSample <- RawData

## 50 Pre-select spores on FSC thresholds----
    ##### TO ADJUST PARAMETERS	IF NECESSARY
    # hist(RawDataSample$YFPHeightLog, n=100)
    # hist(RawDataSample$RFPHeightLog, n=100)
    # plot(RawDataSample$FSC1HeightLog, RawDataSample$SSC1HeightLog
#          , col=densCols(RawDataSample$FSC1HeightLog,RawDataSample$SSC1HeightLog
#          , nbin=50), cex=0.1, ,main = Name
#          , xlab="FSC1HeightLog", ylab="SSC1HeightLog" )
    if (Continue) {
      Data <- Data[Data$CFPHeightLog > minFluo1Value, ]
      Data <- Data[Data$RFPHeightLog > minFluo2Value, ]
      Data <- Data[Data$YFPHeightLog > minFluo3Value, ]
      Data <- Data[Data$FSC1HeightLog > minSporeFscHLog, ]
      Data <- Data[Data$FSC1HeightLog < maxSporeFscHLog, ]
      Data <- Data[Data$SSC1HeightLog > minSporeSscHLog, ]
      Data <- Data[Data$SSC1HeightLog < maxSporeSscHLog, ]
      DataNumber <- nrow(Data)
      res$NbPreselected <- DataNumber
    }

## 60 Sub-sample pre-selected events----
    if (Continue) {
      DataNumberComment <- "OK"
      if(DataNumber < minNbSpores) {
        DataNumberComment <- "Low"
        msg <- paste0("Not enough preselected events (",DataNumber
                      ,") before spore gating")
        cat(paste0(msg,"\n")) ; write(msg, file=logFile, append=TRUE)
        Comment <- "-"
        res$StatusGating <- "Not enough preselected events before spore gating"
        main <- paste0("Selecting spores    ",sampleName)
        plot (0, 0, type='n', main=main, #sub=sub,
                              xlab="FSCHeightLog", ylab="SSCHeightLog" )
        text (0,0,"Not enough preselected events before spore gating", cex=1.4)
        Continue <- FALSE
      } else {
        if (DataNumber < nbUsedForGating) nbUsedForGating <- DataNumber
        DataSample <- Data[sample(nrow(Data), nbUsedForGating), ]
      }
    }

## 70 Plot FSC/SSC to selects Spores based on peak of FSC density----
    if (Continue) {
      DataSampleSpore <- DataSample
      DataSpore <- Data
                  if (firstSelectOnFSC) {  ###  NOT USED ANY MORE
                    FSCDensity <- data.frame(
                      FSCIntensity=density(DataSample$FSC1HeightLog)$x,
                      Density = density(DataSample$FSC1HeightLog)$y )
                    FSCDensitySmooth <- data.frame(
                      FSCIntensitySmooth=smooth.spline(FSCDensity, df=50)$x,
                      FSCDensitySmooth = smooth.spline(FSCDensity, df=50)$y)
                    PeaksValleys <- FindPeaksAndValleys(
                         intensityValues=FSCDensitySmooth$FSCDensitySmooth,
                         nbSeeds=100, graphose=FALSE, LineEachOptim=FALSE, Debug=FALSE)
                    HigherPeakX <- PeaksValleys$Peaks$ValueX_Peak[1]
                    Valleys <- PeaksValleys$Valleys[
                            order(PeaksValleys$Valleys$ValueX_Valley, decreasing=TRUE), ]
                    LeftLimit <- max(Valleys[
                            which(Valleys$ValueX_Valley < HigherPeakX),"ValueX_Valley" ])
                    RightLimit <- min(Valleys[
                            which(Valleys$ValueX_Valley > HigherPeakX),"ValueX_Valley" ])
                    DataSampleSpore <- DataSample[which(
                        DataSample$FSC1HeightLog > LeftLimit
                      & DataSample$FSC1HeightLog < RightLimit) , ]
                # Consider only data in widthAroundPeak
                    MeanFSC <- mean  (DataSampleSpore$FSC1HeightLog)
                    NbData  <- nbUsedForGating
                    VarFSC  <- var  (DataSampleSpore$FSC1HeightLog)
                    EctFSC  <- (VarFSC)^(1/2)
                    FSCCiSup <- HigherPeakX + widthAroundPeak * EctFSC
                    FSCCiInf <- HigherPeakX - widthAroundPeak * EctFSC
                # Select Data in this Interval
                    DataSampleSpore <- DataSampleSpore[which(
                           DataSampleSpore$FSC1HeightLog > FSCCiInf
                         & DataSampleSpore$FSC1HeightLog < FSCCiSup) , ]
                    DataSpore <- Data[which(
                           Data$FSC1HeightLog > FSCCiInf
                         & Data$FSC1HeightLog < FSCCiSup) , ]
                  }
    # Select spores based on FSC and SSC density gates
      DataSporeNumber <- nrow(DataSpore)
      X <- DataSpore$FSC1HeightLog
      Y <- DataSpore$SSC1HeightLog
      graphose=F
      SelectCloudGeometry <- "Circle"
      nbDataForCalc <- nbUsedForGating
      SelectSporesResult <- SelectionOnDensity(X, Y, binNumberX, binNumberY,
                                               graphose, SelectCloudGeometry,
                                               nbDataForCalc, slopeFindPeak)
  # Select spores in DataSampleSpore
      DataSampleSpore <- DataSampleSpore[
            DataSampleSpore$FSC1HeightLog %in% SelectSporesResult$SelData$X , ]
      DataSampleSpore <- DataSampleSpore[
            DataSampleSpore$SSC1HeightLog %in% SelectSporesResult$SelData$Y , ]
  # Select spores in DataSpore
      DataSpore <- DataSpore[
            DataSpore$FSC1HeightLog %in% SelectSporesResult$SelData$X , ]
      DataSpore <- DataSpore[
            DataSpore$SSC1HeightLog %in% SelectSporesResult$SelData$Y , ]
      DataSporeNumber <- nrow(DataSpore)
  # Plot FSC.H / SSC.H
      main <- paste0("Selecting spores    ",sampleName)
      plot(RawDataSample$FSC1HeightLog, RawDataSample$SSC1HeightLog,
           col=densCols(RawDataSample$FSC1HeightLog,
                        RawDataSample$SSC1HeightLog, nbin=30),
           cex=0.1, ,main=main,
           xlab="FSC Height Log", ylab="SSC Height Log" )
      points(DataSampleSpore$FSC1HeightLog,
             DataSampleSpore$SSC1HeightLog,
             col="green", pch=0, cex=0.2)
      NbSelected <- nrow(DataSpore)
      PercentSelected <- round (NbSelected *100 / DataNumber, 1)
      bg <- "transparent"
      legend("topleft", legend=paste0("Total events: ", DataNumber),
             text.col="red" , box.lty=0, bg=bg)
      legend("topright", legend=paste0("Spores: ", NbSelected),
             text.col="red" , box.lty=0, bg=bg, xjust=1)
      legend("bottomright", legend=paste0("Selected: ",
                                         PercentSelected, "%"),
             text.col="red" , box.lty=0, bg=bg, xjust=1)

  # Do not calculate anything if not enough spores
      if(DataSporeNumber < minNbSpores) {
        res$StatusGating <- "Not enough gated spores"
        msg <- paste0("Not enough gated spores (",DataSporeNumber,")")
        cat(paste0(msg,"\n")) ; write(msg, file=logFile, append=TRUE)
        plot(0,0, type='n', xaxt='n', yaxt='n', xlab="", ylab="")
        text (0,0,"Not enough gated spores", cex=1.4)
        Continue <- FALSE
        DataSporeSingleNumber <- 0
      }
      res$NbSpores <- DataSporeNumber
      if(DataSporeNumber < nbDataForCalc) {
        nbDataForCalc <- nbUsedForGating <- DataSporeNumber
        msg <- paste0("Only ",DataSporeNumber,
                      "single spores. Set 'nbUsedForGating' to that value")
        cat(msg) ; write(msg, file=logFile, append=TRUE)
      }
    }

## 80 Plot SSC.A/SSC.H and select singlet spores----
    if(Continue) {  # Continue if enough spores
      X <- DataSpore$SSC1AreaLog
      Y <- DataSpore$SSC1HeightLog
      graphose <- F
      SelectCloudGeometry <- "Axis"
      nbDataForCalc <- nbUsedForGating
      SelectSporesSingleResult <- SelectionOnDensity (
           X=X, Y=Y, binNumberX=binNumberX, binNumberY=binNumberY,
            graphose=graphose , SelectCloudGeometry=SelectCloudGeometry,
            nbDataForCalc=nbDataForCalc, slopeFindPeak=slopeFindPeak)
    # Select Single in DataSampleSpore
      DataSampleSporeSingle <- DataSampleSpore[
        DataSampleSpore$SSC1AreaLog %in% SelectSporesSingleResult$SelData$X , ]
      DataSampleSporeSingle <- DataSampleSporeSingle[
        DataSampleSporeSingle$SSC1HeightLog %in% SelectSporesSingleResult$SelData$Y , ]
      DataSporeSingle <- DataSpore[
        DataSpore$SSC1AreaLog %in% SelectSporesSingleResult$SelData$X , ]
      DataSporeSingle <- DataSporeSingle[
        DataSporeSingle$SSC1HeightLog %in% SelectSporesSingleResult$SelData$Y , ]
      DataSporeSingleNumber <- length(DataSporeSingle$SSC1HeightLog)
      DataSporeSinglePercent <- round(DataSporeSingleNumber/DataSporeNumber *100, 1)
  # Plots Spores Singlets SSC.A / SSC.H
      main <- "Selecting spore singlets"
      if (nrow(DataSampleSpore)>1) {
        plot(DataSampleSpore$SSC1AreaLog, DataSampleSpore$SSC1HeightLog,
                  col=densCols(DataSampleSpore$SSC1AreaLog,
                  DataSampleSpore$SSC1HeightLog, nbin=30),
             main=main, #sub=sub,
             cex=0.1, , xlab="SSC Area Log", ylab="SSC Height Log" )
        points(DataSampleSporeSingle$SSC1AreaLog,
               DataSampleSporeSingle$SSC1HeightLog,
               col="green", pch=0, cex=0.1)
        bg <- "transparent"
        legend("topleft",
            legend=paste0("Total spores: ",DataSporeNumber),
            text.col="red" , box.lty=0, bg=bg)
        legend("topright",
            legend=paste0("Single spores: ",DataSporeSingleNumber),
            text.col="red", box.lty=0, bg=bg)
        legend("bottomright",
            legend=paste0("Selected: ",DataSporeSinglePercent,"%"),
            text.col="red" , box.lty=0, bg=bg)
      } else stop(" Only one data to plot SSC.A / SSC.H")
  # Do not calculate anything if not enough single spores
      if(DataSporeSingleNumber < minNbSpores) {
        res$StatusGating <- "Not enough single spores"
        msg <- paste0("Not enough single spores (",DataSporeSingleNumber,")")
        cat(msg) ; write(msg, file=logFile, append=TRUE)
        
        plot(0,0, type='n', xaxt='n', yaxt='n', xlab="", ylab="")
        text (0,0,"Not enough single spores", cex=1.4)
        Continue <- FALSE
      }
      res$NbSingletSpores <- DataSporeSingleNumber
      if (DataSporeSingleNumber < nbUsedForGating) 
                               nbUsedForGating <- DataSporeSingleNumber
    }

## 90 Copy before removing  spores from NFxNF diploids----
    if(Continue) {
      RawDataSporeSingle <- DataSporeSingle
      if(! discardNFSpores) {
        Results <- NULL
        Results$Log <- "Ok"
      }
    }

## 100 Discard NF spores outside fluo peaks (spores from NFxNF diploids)----
    if(Continue && discardNFSpores) {   ##  discardNFSpores ALWAYS TRUE
  # Remove NA events corresponding to non-valid YFP peaks
      threeMarkers <- c("CFPHeightLog", "RFPHeightLog", "YFPHeightLog")
      threeCodes <- c("C", "R", "Y")
      threeColors <- c("green","red","orange")
      threeFluos <- c(fluo1H[1], fluo2H[1], fluo3H[1])
      allMarkers <- allColors <- allFluos <- c()
      iM <- 1
      for (iM in 1:length(unlist(strsplit(Order,"")))) {
        sel <- threeCodes==unlist(strsplit(Order,""))[iM]
        allMarkers <- c(allMarkers, threeMarkers[sel])
        allColors <- c(allColors, threeColors[sel])
        allFluos <- c(allFluos, threeFluos[sel])
      }
  # Loop on markers ================-
      iMarker <- 1
      for (iMarker in 1:length(unlist(strsplit(Order,"")))) {
        Marker <- allMarkers[iMarker]
        markerColor <- allColors[iMarker]
        ThisData <- DataSporeSingle[ , c("TIME", Marker)]
        colnames(ThisData) <- c("TIME","Intensity")
        eventsID <- ThisData$TIME
        eventsIntensity <- ThisData$Intensity
        nbSeeds=100
        graphose=FALSE
        nbFluoClasses <- 2
        Results <- IntensityCluster(markerName=markerName, markerColor=markerColor
                                  , eventsID=eventsID, eventsIntensity=eventsIntensity
                                  , nbSeeds=nbSeeds, graphose=F, nbFluoClasses=nbFluoClasses
                                  , cutOff=cutOff, DataX=DataX, DataY=DataY)
        if(Results$Log == "ERROR: Nb of peaks detected < nbFluoClasses") {
          res$StatusGating <- "Only one Fluo peak detected"
          msg <- paste0("Only one Fluo peak detected")
          cat(paste0(msg,"\n")) ; write(msg, file=logFile, append=TRUE)
          
          ThisRawData <- RawDataSporeSingle[ , c("TIME", Marker) ]
          colnames(ThisRawData) <- c("TIME","Intensity")
          eventsID <- ThisRawData$TIME
          eventsIntensity <- ThisRawData$Intensity
          RawData <- data.frame(eventsID=eventsID,
                                eventsIntensity=eventsIntensity)
          RawDensityData <- data.frame(
            MarkerIntensity=density(RawData$eventsIntensity)$x,
            Density = density(RawData$eventsIntensity)$y )
          rawX <- RawDensityData$MarkerIntensity
          rawY <-RawDensityData$Density
          xLim <- range(c(X, rawX))
          yLim <- range(c(Y, rawY)*1.1)
          letter <- c("A","B","C")[iMarker]
          if (Marker=="CFPHeightLog") main <- paste0(
                              "Marker ",letter,": ", fluo1H[1] -> trueMarker)
          if (Marker=="RFPHeightLog") main <- paste0(
                              "Marker ",letter,": ", fluo2H[1] -> trueMarker)
          if (Marker=="YFPHeightLog") main <- paste0(
                              "Marker ",letter,": ", fluo3H[1] -> trueMarker)
          plot(rawX, rawY, cex=0.2, type='l', col="red", xlim=xLim, ylim=yLim,
               xlab="Marker intensity", ylab="Density", main=main)
      # Displays message in empty plot
          plot(0,0, type='n', xaxt='n', yaxt='n', xlab="", ylab="")
          text (0,0,paste0("Only one peak detected in ",allFluos[iMarker]), cex=1.4)
          Continue <- FALSE
        } else if(Results$Log == "Ok") {  # then delete NA events
          MonoDistorsionResult <- NULL
 #         DataSporeSingle <- DataSporeSingle[order(DataSporeSingle$TIME), ]
          AssignResults <- data.frame(
                    TIME=DataSporeSingle$TIME,
                    ThisClass=rep("NA",length(DataSporeSingle$TIME)))
          AssignResults$ThisClass <- Results$AssignResults[, "fluoClass"]
 #         AssignResults <- AssignResults[order(AssignResults$TIME), ]
          a <- merge (DataSporeSingle, AssignResults, by="TIME")
          # Removes all data outside the 2 identified peaks
          b <- a[which( a$ThisClass != "NA"), ]
          # Keep only events of YFP class 1 or 2 (from right to left !!!)
          # removes the column
          DataSporeSingle <- b[, which(colnames(b)!="ThisClass")]
          NbYFPDeletedData <- nrow(RawDataSporeSingle)-nrow(DataSporeSingle)
        } else { stop(" Wrong output in Results$Log")}
      }
    }

## 120 Corresp. ABC <=> CRY----
    if(Continue) {
      noFluo3 <-  ! (is.character(fluo3H[3]) || fluo3H[3]==""
                                             || is.na(fluo3H[3])
                                             || is.null(fluo3H[3]))
      only2Markers <- ( length(unlist(strsplit(Order,""))) == 2 )
      if ( ! only2Markers && noFluo3)
        stop(paste0(" No fluo3H marker declared, but markers ",
                                          Markers," found in sample name"))
      if (only2Markers) {
        DfFluo <- data.frame(Marker=c("A","B"), Fluo=unlist(strsplit(Order,"")))
      } else {   ##  DfFluo$Marker is NEVER USED ?
        DfFluo <- data.frame(Marker=c("A","B","C"), Fluo=unlist(strsplit(Order,"")))
      }
      DfFluo$FluoCode <- paste0(DfFluo$Fluo,"FPHeightLog")
      MarkersToUse <- c(DfFluo$FluoCode)  # Choose fluo colors to analyse
    }

## 125 Plot 2D fluo intensity if only 2 markers used----
    if (Continue && length(MarkersToUse) == 2) {
      DataX <- unlist(DataSampleSporeSingle[
            which(colnames (DataSampleSporeSingle)==MarkersToUse[1])])
      DataY <- unlist(DataSampleSporeSingle[
            which(colnames (DataSampleSporeSingle)==MarkersToUse[2])])
      trueMarker <- c(NA,NA)
      for (i in 1:2) {
        if (MarkersToUse[i]=="CFPHeightLog") trueMarker[i] <- fluo1H[1]
        if (MarkersToUse[i]=="RFPHeightLog") trueMarker[i] <- fluo2H[1]
        if (MarkersToUse[i]=="YFPHeightLog") trueMarker[i] <- fluo3H[1]
      }
      plot(DataX, DataY, cex=0.2, xlab=paste0("Marker A: ",trueMarker[1])
                                , ylab=paste0("Marker B: ",trueMarker[2])
                                , main = "Fluorescence intensities")
    }

## 130 Gets fluo classes of events----
    if(Continue)  {
      MonoDistorsionResult <- NULL
 #     DataSporeSingle <- DataSporeSingle[order(DataSporeSingle$TIME), ]
      AssignResults <- data.frame(TIME=DataSporeSingle$TIME,
                      CFP_Class=rep("NA",length(DataSporeSingle$TIME)),
                      RFP_Class=rep("NA",length(DataSporeSingle$TIME)),
                      YFP_Class=rep("NA",length(DataSporeSingle$TIME)))
      Marker <- MarkersToUse[1]
      for (Marker in MarkersToUse) {
        msg <- paste0 ("Marker: ", Marker)
        cat(paste0(msg,"\n")) ; write(msg, file=logFile, append=TRUE)
        markerName <- paste0(unlist(strsplit(Marker,""))[1],
                             unlist(strsplit(Marker,""))[2],
                             unlist(strsplit(Marker,""))[3])
        if (markerName=="CFP") { markerColor="green"}
        if (markerName=="RFP") { markerColor="red"}
        if (markerName=="YFP") { markerColor="orange"}
        ThisData <- DataSporeSingle[ , c("TIME", Marker) ]
        colnames (ThisData) <- c("TIME","Intensity")
        eventsID <- ThisData$TIME
        eventsIntensity <- ThisData$Intensity
        nbSeeds <- 100
        graphose <- FALSE
        nbFluoClasses <- 2
        Results <- IntensityCluster(markerName, markerColor, eventsID,
                                    eventsIntensity, nbSeeds, graphose=F,
                                    nbFluoClasses, cutOff, 
                                    DataX=DataX, DataY=DataY)
        if(Results$Log == "ERROR: Nb of peaks detected < nbFluoClasses") {
          res$StatusGating <- "Only one Fluo peak detected"
          plot(0,0, type='n', xaxt='n', yaxt='n', xlab="", ylab="")
          text (0,0,paste0("Only one Fluo peak detected in ",trueMarker), cex=1.4)
          Continue <- FALSE
        }

## 135 Plot 1D densities----
        if(Continue) {
      # Density histogram of all singlet spores (raw data)
          ThisRawData <- RawDataSporeSingle[ , c("TIME", Marker) ]
          colnames(ThisRawData) <- c("TIME","Intensity")
          eventsID <- ThisRawData$TIME
          eventsIntensity <- ThisRawData$Intensity
          RawData <- data.frame(eventsID=eventsID,
                                         eventsIntensity=eventsIntensity)
          RawDensityData <- data.frame(
                MarkerIntensity=density(RawData$eventsIntensity)$x,
                Density = density(RawData$eventsIntensity)$y )
          rawX <- RawDensityData$MarkerIntensity
          rawY <-RawDensityData$Density
      # Density histogram of spores after discarding NF spores
          ThisData <- DataSporeSingle[ , c("TIME", Marker) ]
          colnames(ThisData) <- c("TIME","Intensity")
          eventsID <- ThisData$TIME
          eventsIntensity <- ThisData$Intensity
          Data <- data.frame(eventsID=eventsID,
                             eventsIntensity=eventsIntensity)
          DensityData <- data.frame(
            MarkerIntensity=density(Data$eventsIntensity)$x,
            Density = density(Data$eventsIntensity)$y )
          X <- DensityData$MarkerIntensity
          Y <- DensityData$Density
          if (length(MarkersToUse)==3) allLetters <- c("A","B","C")
          if (length(MarkersToUse)==2) allLetters <- c("A","B")
          letter <- allLetters[MarkersToUse==Marker]
          if (markerName=="CFP") trueMarker <- paste0(
                                      "Marker ",letter,": ", fluo1H[1])
          if (markerName=="RFP") trueMarker <- paste0(
                                      "Marker ",letter,": ", fluo2H[1])
          if (markerName=="YFP") trueMarker <- paste0(
                                      "Marker ",letter,": ", fluo3H[1])
          xLim <- range(c(X, rawX))
          yLim <- range(c(Y, rawY)*1.1)
      # Plot density without spores from NFxNF diploids (first peak RFP/YF)
          plot(X, Y, cex=0.2, type='l', col="red", xlim=xLim, ylim=yLim,
               xlab="Marker intensity", ylab="Density", main=trueMarker)
      # Then adds line with all singlet spores
          points(rawX, rawY, cex=0.2, col="black", type='l')

## 137 Peak detect FULL distribution----
          if ( ! halfDistrib) {
            # SELECT ONLY DATA WHICH ARE IN STD * widthAroundPeak
            # For Class 1
            MeanIntensityClass1 <- mean(Results$AssignResults$eventsIntensity[
                          Results$AssignResults$fluoClass=="1"])
            NbDataClass1 <- length(Results$AssignResults$eventsIntensity[
                          Results$AssignResults$fluoClass=="1"])
            VarIntensityClass1 <- var(Results$AssignResults$eventsIntensity[
                          Results$AssignResults$fluoClass=="1"])
            EctIntensityClass1 <- (VarIntensityClass1)^(1/2)
            # Must be centered on the highest peak
            XposHighestPeakClass1 <- Results$HighestPeaksDf$HighestPeakX[1]
            YposHighestPeakClass1 <- Results$HighestPeaksDf$HighestPeakY[1]
            abline(v= XposHighestPeakClass1, lty=1,  col="purple")
            CiSupIntensityClass1 <- XposHighestPeakClass1+widthAroundPeak*EctIntensityClass1
            CiInfIntensityClass1 <- XposHighestPeakClass1-widthAroundPeak*EctIntensityClass1
            abline(v= CiSupIntensityClass1, lty=2,  col="purple")
            abline(v= CiInfIntensityClass1, lty=2,  col="purple")
            # For Class 2
            MeanIntensityClass2 <- mean (Results$AssignResults$eventsIntensity[
                          Results$AssignResults$fluoClass=="2"])
            NbDataClass2  <- length(Results$AssignResults$eventsIntensity[
                          Results$AssignResults$fluoClass=="2"])
            VarIntensityClass2  <- var(Results$AssignResults$eventsIntensity[
                          Results$AssignResults$fluoClass=="2"])
            EctIntensityClass2  <- (VarIntensityClass2)^(1/2)
            # Must be centered on the highest peak
            XposHighestPeakClass2 <- Results$HighestPeaksDf$HighestPeakX[2]
            YposHighestPeakClass2 <- Results$HighestPeaksDf$HighestPeakY[2]
            abline(v= XposHighestPeakClass2, lty=1,  col="purple")
            CiSupIntensityClass2 <- XposHighestPeakClass2+widthAroundPeak*EctIntensityClass2
            CiInfIntensityClass2 <- XposHighestPeakClass2-widthAroundPeak*EctIntensityClass2
            abline(v= CiSupIntensityClass2, lty=2,  col="purple")
            abline(v= CiInfIntensityClass2, lty=2,  col="purple")
            #Discard info on the class of Fluo to Re-assign class
            Results$AssignResults$fluoClass <- "NA"
            #Assign Data which are in Class 1
            Results$AssignResults$fluoClass[
                Results$AssignResults$eventsIntensity > CiInfIntensityClass1
              & Results$AssignResults$eventsIntensity < CiSupIntensityClass1] <- "1"
            #Assign Data which are in Class 2
            Results$AssignResults$fluoClass[
                Results$AssignResults$eventsIntensity > CiInfIntensityClass2
              & Results$AssignResults$eventsIntensity < CiSupIntensityClass2] <- "2"
        # Add widthAroundPeak value on graph
        # Add Class of Fluo to Data
            if (markerName=="CFP") AssignResults$CFP_Class <- Results$AssignResults[,"fluoClass"]
            if (markerName=="RFP") AssignResults$RFP_Class <- Results$AssignResults[,"fluoClass"]
            if (markerName=="YFP") AssignResults$YFP_Class <- Results$AssignResults[,"fluoClass"]
            # Print the number of selected data
            NbDataClass1 <- sum(Results$AssignResults$fluoClass=="1")
            NbDataClass2 <- sum(Results$AssignResults$fluoClass=="2")
            text(x=XposHighestPeakClass1*1.03, y=YposHighestPeakClass1*1.11,
                 labels=paste0("Nb Class1: ",NbDataClass1), pos=1)
            text(x=XposHighestPeakClass2, y=YposHighestPeakClass2*1.00,
                 labels=paste0("Nb Class2: ",NbDataClass2), pos=2)
            # Segregation distortion
            FreqClass1 <- round(NbDataClass1/(NbDataClass1+NbDataClass2),2)
            FreqClass2 <- round(NbDataClass2/(NbDataClass1+NbDataClass2),2)
            text(x=XposHighestPeakClass1*1.03, y=YposHighestPeakClass1*1.16,
                            labels=paste0("Freq Class1: ",FreqClass1), pos=1)
            text(x=XposHighestPeakClass2, y=YposHighestPeakClass2*1.05,
                            labels=paste0("Freq Class2: ",FreqClass2), pos=2)
            myDf <- data.frame (markerName=markerName, FreqClass1 = FreqClass1)
            MonoDistorsionResult <- rbind.data.frame (MonoDistorsionResult, myDf)
          }

## 138 Peak detect HALF distribution----
          if (halfDistrib) {
            # SELECT ONLY HALF DATA WHICH ARE IN STD * widthAroundPeak
            # For Class 1
            MeanIntensityClass1 <- mean(Results$AssignResults$eventsIntensity[
                                  Results$AssignResults$fluoClass=="1"])
            NbDataClass1  <- length(Results$AssignResults$eventsIntensity[
                                  Results$AssignResults$fluoClass=="1"])
            VarIntensityClass1  <- var(Results$AssignResults$eventsIntensity[
                                  Results$AssignResults$fluoClass=="1"])
            EctIntensityClass1  <- (VarIntensityClass1)^(1/2)
            #Il faut centrer sur le pic le plus haut !
            XposHighestPeakClass1 <- Results$HighestPeaksDf$HighestPeakX[1]
            abline(v= XposHighestPeakClass1, lty=1,  col="Red")
            CiSupIntensityClass1 <- XposHighestPeakClass1+widthAroundPeak*EctIntensityClass1
            abline(v= CiSupIntensityClass1, lty=2,  col="purple")
            #For Class 2
            MeanIntensityClass2 <- mean(Results$AssignResults$eventsIntensity[
                              Results$AssignResults$fluoClass=="2"])
            NbDataClass2  <- length(Results$AssignResults$eventsIntensity[
                              Results$AssignResults$fluoClass=="2"])
            VarIntensityClass2  <- var(Results$AssignResults$eventsIntensity[
                              Results$AssignResults$fluoClass=="2"])
            EctIntensityClass2  <- (VarIntensityClass2)^(1/2)
            # Center on highest peak
            XposHighestPeakClass2 <- Results$HighestPeaksDf$HighestPeakX[2]
            abline(v= XposHighestPeakClass2, lty=1,  col="Red")
            CiInfIntensityClass2 <- XposHighestPeakClass2-widthAroundPeak*EctIntensityClass2
            abline(v= CiInfIntensityClass2, lty=2,  col="purple")
            # Discard info on the class of Fluo to Re-assign class
            Results$AssignResults$fluoClass <- "NA"
            # Assign Data which are in Class 1
            Results$AssignResults$fluoClass[
                Results$AssignResults$eventsIntensity > XposHighestPeakClass1
              & Results$AssignResults$eventsIntensity < CiSupIntensityClass1] <- "1"
            # Assign Data which are in Class 2
            Results$AssignResults$fluoClass[
                Results$AssignResults$eventsIntensity > CiInfIntensityClass2
              & Results$AssignResults$eventsIntensity < XposHighestPeakClass2] <- "2"
            # Add widthAroundPeak value on graph
            # Add Class of Fluo to Data
            if (markerName=="CFP") AssignResults$CFP_Class <- Results$AssignResults[,"fluoClass"]
            if (markerName=="RFP") AssignResults$RFP_Class <- Results$AssignResults[,"fluoClass"]
            if (markerName=="YFP") AssignResults$YFP_Class <- Results$AssignResults[,"fluoClass"]
            # Print the number of selected data
            XposHighestPeakClass1 <- Results$HighestPeaksDf$HighestPeakX[1]
            YposHighestPeakClass1 <- Results$HighestPeaksDf$HighestPeakY[1]
            XposHighestPeakClass2 <- Results$HighestPeaksDf$HighestPeakX[2]
            YposHighestPeakClass2 <- Results$HighestPeaksDf$HighestPeakY[2]
            NbDataClass1 <- sum(Results$AssignResults$fluoClass=="1")
            NbDataClass2 <- sum(Results$AssignResults$fluoClass=="2")
            text(x=XposHighestPeakClass1*1.03, y=YposHighestPeakClass1*1.11,
                 labels=paste0("Nb Class1: ",NbDataClass1), pos=1)
            text(x=XposHighestPeakClass2, y=YposHighestPeakClass2*1.00,
                 labels=paste0("Nb Class2: ",NbDataClass2), pos=2)
            # Segregation distortion
            FreqClass1 <- round(NbDataClass1/(NbDataClass1+NbDataClass2),2)
            FreqClass2 <- round(NbDataClass2/(NbDataClass1+NbDataClass2),2)
            text(x=XposHighestPeakClass1*1.03, y=YposHighestPeakClass1*1.16,
                 labels=paste0("Freq Class1: ",FreqClass1), pos=1)
            text(x=XposHighestPeakClass2, y=YposHighestPeakClass2*1.05,
                 labels=paste0("Freq Class2: ",FreqClass2), pos=2)
            myDf <- data.frame (markerName=markerName, FreqClass1 = FreqClass1)
            MonoDistorsionResult <- rbind.data.frame (MonoDistorsionResult, myDf)
          }
        }
      }
    }

## 148 Segregation Distortion----
    if(Continue) {
      YFPDistor <- MonoDistorsionResult[MonoDistorsionResult$markerName == "YFP", "FreqClass1"]
      RFPDistor <- MonoDistorsionResult[MonoDistorsionResult$markerName == "RFP", "FreqClass1"]
      CFPDistor <- MonoDistorsionResult[MonoDistorsionResult$markerName == "CFP", "FreqClass1"]
      # make a data frame with the number of spores in each fluorescent class
      # If three markers are used
    }

## 150 Rec rates for 3 markers----
    if(Continue && length(unlist(strsplit(Order,"")))==3) {
      msg <- paste0("Computing naive recombination rates for "
                    ,length(unlist(strsplit(Order,"")))," markers")
      cat(paste0(msg,"\n"))
      write(msg, file=logFile, append=TRUE)
      RYC <- RCY <- YRC <- YCR <- CYR <- CRY <- length(which(
              AssignResults$CFP_Class=="1"
              & AssignResults$RFP_Class=="1"
              & AssignResults$YFP_Class=="1"))
      NNN <- length(which(
              AssignResults$CFP_Class=="2"
              & AssignResults$RFP_Class=="2"
              & AssignResults$YFP_Class == "2"))
      R   <- length(which(
              AssignResults$CFP_Class == "2"
              & AssignResults$RFP_Class == "1"
              & AssignResults$YFP_Class == "2"))
      YC  <- CY <- length(which(
              AssignResults$CFP_Class == "1"
              & AssignResults$RFP_Class == "2"
              & AssignResults$YFP_Class == "1"))
      RC  <- CR <- length(which(
              AssignResults$CFP_Class == "1"
              & AssignResults$RFP_Class == "1"
              & AssignResults$YFP_Class == "2"))
      Y   <- length(which(
              AssignResults$CFP_Class == "2"
              & AssignResults$RFP_Class == "2"
              & AssignResults$YFP_Class == "1"))
      RY  <- YR <- length(which(
              AssignResults$CFP_Class == "2"
              & AssignResults$RFP_Class == "1"
              & AssignResults$YFP_Class == "1"))
      C   <- length(which(
              AssignResults$CFP_Class == "1"
              & AssignResults$RFP_Class == "2"
              & AssignResults$YFP_Class == "2"))
      Tot <- RYC + NNN + R + YC + RC + Y + RY + C
      NbSpores <- c(RYC,RCY,YRC,YCR,CYR,CRY,NNN,R,YC,CY,RC,CR,Y,RY,YR,C,Tot)
      Class    <- c("RYC","RCY","YRC","YCR","CYR","CRY","NNN","R","YC","CY","RC","CR","Y","RY","YR","C","Tot")
      DfData  <- data.frame (Class, NbSpores)
      #Attribute each class to recombinant or parental class in case of ABC
      MarqA <- strsplit (Order, "")[[1]] [1]
      MarqB <- strsplit (Order, "")[[1]] [2]
      MarqC <- strsplit (Order, "")[[1]] [3]
      MarqN <- "N"
      ABC <- DfData[which(DfData$Class== paste (MarqA,MarqB,MarqC, sep="")), "NbSpores" ]
      NF  <- DfData[which(DfData$Class== paste (MarqN,MarqN,MarqN, sep="")), "NbSpores" ]
      A   <- DfData[which(DfData$Class== MarqA), "NbSpores" ]
      BC  <- DfData[which(DfData$Class== paste (MarqB,MarqC, sep="")), "NbSpores" ]
      AB  <- DfData[which(DfData$Class== paste (MarqA,MarqB, sep="")), "NbSpores" ]
      C   <- DfData[which(DfData$Class== MarqC), "NbSpores" ]
      AC  <- DfData[which(DfData$Class== paste (MarqA,MarqC, sep="")), "NbSpores" ]
      B   <- DfData[which(DfData$Class== MarqB), "NbSpores" ]
      #Distorsion calculation
      FreqABCvsPPP <- ABC / (NF+ABC)
      FreqBCvsA    <- BC  / (A+BC)
      FreqABvsC    <- AB  / (C+AB)
      FreqACvsB    <- AC  / (B+AC)
      #Rec rate and Naive_CoC in case of markers ABC
      Rec1  <- (A + BC + AC + B) / Tot
      Rec2  <- (AB + C + AC + B) / Tot
      Doubl <- (AC + B) / Tot
      Naive_CoC   <- Doubl / (Rec1*Rec2)
      if (is.nan(Naive_CoC)) Naive_CoC <- -999

    # Writes results in res table
      res[,paste0("Freq_",fluo1H[1])] <- CFPDistor
      res[,paste0("Freq_",fluo2H[1])] <- RFPDistor
      res[,paste0("Freq_",fluo3H[1])] <- YFPDistor
      res$FreqABCvsPPP=FreqABCvsPPP
      res$FreqBCvsA=FreqBCvsA
      res$FreqABvsC=FreqABvsC
      res$FreqACvsB=FreqACvsB
      res$Naive_rAB=Rec1
      res$Naive_rBC=Rec2
      res$Naive_rDouble=Doubl
      res$Naive_CoC=Naive_CoC
      res$StatusFcs=StatusFcs
      res$StatusGating="Data number OK"
      res$App=A
      res$ABp=AB
      res$ABC=ABC
      res$ApC=AC
      res$pBp=B
      res$pBC=BC
      res$ppC=C
      res$ppp=NF
    #   
    #   #Write on graphical window
      plot(0, 0, type="n", main="Recombination results", yaxt="n", xaxt="n",
           xlab="", ylab="", xlim=c(0,10), ylim=c(0,15) )
      text(x=4, y=14, labels="Naive estimates", adj=1)
      text(x=rep(4,8), y=(12:4),
           labels=c(paste0("Naive_rAB: ", round(Rec1,2)),
                    paste0("Naive_rBC: ", round(Rec2,2)),
                    paste0("Naive_CoC: ", round(Naive_CoC,2)),
                    paste0("DoubleRec: ", round(Doubl,2)),
                    "",
                    paste0("FreqABCvsPPP: ",round(FreqABCvsPPP,2)),
                    paste0("FreqBCvsA: ", round(FreqBCvsA,2)),
                    paste0("FreqABvsC: ", round(FreqABvsC,2)),
                    paste0("FreqACvsB: ", round(FreqACvsB,2)))
           , adj=1)
    }

## 160 Rec rates for 2 markers----
    if(Continue && length(unlist(strsplit(Order,"")))==2) {
      msg <- paste0("Computing naive recombination rates for "
                    ,length(unlist(strsplit(Order,"")))," markers")
      cat(paste0(msg,"\n")) ; write(msg, file=logFile, append=TRUE)
      NN <- length (which(AssignResults$CFP_Class == "2"  & AssignResults$YFP_Class == "2"))
      CY <- length (which(AssignResults$CFP_Class == "1"  & AssignResults$YFP_Class == "1"))
      Y  <- length (which(AssignResults$CFP_Class == "2"  & AssignResults$YFP_Class == "1"))
      C  <- length (which(AssignResults$CFP_Class == "1" & AssignResults$YFP_Class == "2"))
      Tot <- CY + Y + C + NN
      NbSpores <- c(CY,NN,Y,C,Tot)
      Class <- c("CY","NN","Y","C","Tot")
      DfData <- data.frame (Class, NbSpores)
      #Attribute each class to recombinant or parental class in case of ABC
      MarqA <- strsplit (Order, "")[[1]] [1]
      MarqB <- strsplit (Order, "")[[1]] [2]
      MarqN <- "N"
      AB  <- DfData[which(DfData$Class== paste (MarqA,MarqB, sep="")), "NbSpores" ]
      NF  <- DfData[which(DfData$Class== paste (MarqN,MarqN, sep="")), "NbSpores" ]
      A   <- DfData[which(DfData$Class== MarqA), "NbSpores" ]
      B   <- DfData[which(DfData$Class== MarqB), "NbSpores" ]
      # Segregation distorsion
      FreqABvsPP <- AB / (NF+AB)
      FreqAvsB    <- A  / (A+B)
      Rec1  <- (A + B) / Tot

    # Writes results in res table
      res[,paste0("Freq_",fluo1H[1])] <- CFPDistor
      res[,paste0("Freq_",fluo2H[1])] <- RFPDistor
      res[,paste0("Freq_",fluo3H[1])] <- YFPDistor
      res$Naive_rAB=Rec1
      res$StatusFcs=StatusFcs
      res$StatusGating="Data number OK"
      res$App=A
      res$ABp=AB
      res$pBp=B
      res$ppp=NF
      
      plot(0, 0, type="n", main="Recombination results", yaxt="n", xaxt="n", xlab="",
             ylab="", xlim=c(0,10), ylim=c(0,15)) # Writes in graphical window
      text(x=4, y=14, labels="Naive estimates", adj=1)
      text(x=rep(4,8), y=(12:4),
           labels = c(paste( "Naive_rAB = ", round(Rec1,2)),
                      "",
                      "",
                      "",
                      "",
                      paste("FreqABCsPP= ", round(FreqABvsPP,2)),
                      paste("FreqAvsB = ", round(FreqAvsB,2)),
                      "",
                      "")
           , adj=1)
    }

## 170 Max Likelihood inference----
    if (Continue && ! only2Markers) {
      msg <- paste0("Computing Max Likelihood recombination rates with 3 markers")
      cat(msg) ; write(msg, file=logFile, append=TRUE)
      res <- MaxLikelihoodFit3loci(Data=res, mlPdfFile=mlPdfFile, logFile=logFile)
      if (is.numeric(c(res$ML_rAB, res$ML_rBC, res$ML_CoC,
                           res$ML_Aext, res$ML_Bext, res$ML_Cext))) {
        dev.set(dev1)
        text(x=9, y=14, labels="ML estimates", adj=1)
        text(x=rep(9,8), y=(12:4),
             labels=c(paste0("ML_Naive_rAB: ",round(res$ML_rAB,2)),
                      paste0("ML_Naive_rBC: ", round(res$ML_rBC,2)),
                      paste0("ML_CoC: ", round(res$ML_CoC,2)),
                      "",
                      "",
                      paste0("ML_extA: ", round(res$ML_Aext,2)),
                      paste0("ML_extB: ", round(res$ML_Bext,2)),
                      paste0("ML_extC: ", round(res$ML_Cext,2)),
                      "")
             , adj=1)
      }
    }
    # if (Continue && only2Markers) {
    #   plot(0, 0, type="n", main="Recombination results", yaxt="n", xaxt="n", xlab="",
    #        ylab="", xlim=c(0,10), ylim=c(0,15)) # Writes in graphical window
    #   text(x=4, y=14, labels="Naive estimates", adj=1)
    #   text(x=rep(4,8), y=(12:4),
    #        labels = c(paste( "Naive_rAB = ", round(Rec1,2)),
    #                   "",
    #                   "",
    #                   "",
    #                   "",
    #                   paste("FreqABCsPP= ", round(FreqABvsPP,2)),
    #                   paste("FreqAvsB = ", round(FreqAvsB,2)),
    #                   "",
    #                   "")
    #        , adj=1)
    # }

## 175 Compares naive vs ML estimates of rec rates and CoC----
    if (Continue && ! only2Markers && res$StatusML=="Ok") {
      okMl <- TRUE
      for (this in 1:3) {
        if (this==1) { nar <- res$Naive_rAB ; mlr <- res$ML_rAB ; ms <- "rAB" }
        if (this==2) { nar <- res$Naive_rBC ; mlr <- res$ML_rBC ; ms <- "rBC" }
        if (this==3) { nar <- res$Naive_CoC ; mlr <- res$ML_CoC ; ms <- "CoC" }
        if (abs(nar-mlr)/nar > 0.1) {
          msg <- paste0("\nWarning: Naive_",ms,"=",nar," but ML_",ms,"=",mlr)
          okMl <- FALSE
          cat(msg) ; write(msg, file=logFile, append=TRUE)
        }
      }
      if ( ! okMl) res$StatusML <- "Warning: Naive-ML difference > 10%"
    }

## 180 Write results file----
    AllResults <- rbind.data.frame (AllResults, res)
    resFile <- file.path(PathResults, paste0(Date,"_",Time,"_ALL_RESULTS.txt"))
    write.table(AllResults, file=resFile, sep="\t", row.names=FALSE)

## 190 Merge CYTO PDF files----
    if (pdfGraphs!="none") {
      dev.off()
      allCytoPdfFiles <- c(allCytoPdfFiles, cytoPdfFile)
      mergedCytoPdfFile <- file.path(PathCytoGraphs,
                                paste0(Date,"_",Time,"_ALL_CYTO_GRAPHS.pdf"))
      
      if (pdfGraphs=="merged_only") {
        if (iFcs==1) {
          qpdf::pdf_combine(input=cytoPdfFile, output=mergedCytoPdfFile)
           file.remove(cytoPdfFile)
        } else {
          file.copy(from=mergedCytoPdfFile, to="aaa.pdf")
          qpdf::pdf_combine(input=c("aaa.pdf", cytoPdfFile), output=mergedCytoPdfFile)
          file.remove("aaa.pdf")
          file.remove(cytoPdfFile)
        }
      } else if (pdfGraphs!="separated") {
       qpdf::pdf_combine(input=allCytoPdfFiles, output=mergedCytoPdfFile)
      }
    }

## 200 Merge ML fit PDF files----
#    if (Continue) {
    if (Continue && ! only2Markers) {
      allMlPdfFiles <- c(allMlPdfFiles, mlPdfFile)
      mergedMlPdfFile <- file.path(PathMlGraphs,
                                 paste0(Date,"_",Time,"_ALL_ML_GRAPHS.pdf"))
      if (pdfGraphs=="merged_only") {
        if (iFcs==1) {
          qpdf::pdf_combine(input=mlPdfFile, output=mergedMlPdfFile)
          file.remove(mlPdfFile)
        } else {
          file.copy(from=mergedMlPdfFile, to="aaa.pdf")
          qpdf::pdf_combine(input=c("aaa.pdf", mlPdfFile), output=mergedMlPdfFile)
          file.remove("aaa.pdf")
          file.remove(mlPdfFile)
        }
      } else if (pdfGraphs!="separated") {
        qpdf::pdf_combine(input=allMlPdfFiles, output=mergedMlPdfFile)
      }
    }

    cat("\n\n") ; write("", file=logFile, append=TRUE)
  }    #  END OF MAIN LOOP
  return(AllResults)
}
```
